# Integrated quantitative imaging and biomechanical modeling of early gastrulation in *C. elegans*

**DOI:** 10.64898/2026.03.30.715391

**Authors:** Wim Thiels, Michiel Vanslambrouck, Casper Van Bavel, Keyu Xiao, Jef Vangheel, Bart Smeets, Rob Jelier

## Abstract

The stereotyped internalization of two endodermal precursors during early *Caenorhabditis elegans* gastrulation enables quantitative dissection of cell ingression mechanics. Experimental work has shown that apical constriction drives Ea and Ep ingression, and several molecular features involved have been identified. Yet, no integrative mechanical analysis has assessed how these elements collectively produce the observed behavior. To address this, we combined biomechanical simulations with a comprehensive dataset of 3D-segmented cell meshes, some with cortical protein distributions, to analyze the mechanics of ingression in its in-vivo context. Our analysis shows the process starts shortly after birth of the ingressing cells. A cortical flow drives the formation of an E-cadherin-rich structure at the apical Ea-Ep interface, which contributes to localizing the buildup of apical tension. Simulations show that medioapical actomyosin contraction can reproduce the observed ingression movements and suggest force transmission to neighboring cells via a friction-based ‘molecular clutch’ at the apical ring of contact. A series of concurrent cell divisions facilitates ingression, and their stereotyped planar orientation also contributes. Furthermore, we observe an embryo-wide movement of cells during gastrulation. This movement resembles a flow, suggesting that local force generation leads to global rearrangements via internal pressure changes. Finally, at the end of ingression, detailed microscopy shows that neighboring cells actively close the gastrulation cleft by forming a rosette-like configuration and extending actin-rich protrusions. In conclusion, our integrated mechanical description of gastrulation shows that successful ingression is driven by apical constriction and supported by localized friction-based force transmission, coordinated stereotyped cell divisions, and the resulting global tissue flow.

## 2 Introduction

Gastrulation is a conserved morphogenetic process in embryonic development that establishes the fundamental body plan by rearranging cells into distinct germ layers (Martindale, 2005; Keller et al., 2003). In *C. elegans*, gastrulation begins at the 26 cell stage with the stereotyped internalization of two endodermal precursor cells, Ea and Ep (Nance, 2005), driven by apical constriction of their contact-free surfaces (Goldstein and Nance, 2020). This process provides a tractable system for studying conserved cellular ingression, as it can be readily imaged and experimentally perturbed. Apical constriction is generated by locally heightened actomyosin contractility in the apical domain of these cells (Roh-Johnson et al., 2012).

The signaling cascade leading up to these cortical contractions depends on HMR-1/E-cadherin, which is basolaterally positioned at cell-cell contacts (Klompstra et al., 2015). HMR-1 recruits the RhoGAP PAC-1, which locally inactivates the Rho GTPase CDC-42, limiting active CDC-42 to the apical surface (Klompstra et al., 2015). Apical CDC-42 then recruits the myosin light-chain kinase MRCK-1, which activates non-muscle myosin II (NMY-2, hereafter myosin) by phosphorylating regulatory light chains, increasing cortical tension and driving apical constriction (Marston et al., 2016). Depletion of CDC-42, MRCK-1, or myosin inhibits apical constriction and ingression (Marston et al., 2016). Depletion of HMR-1 results in delayed ingression, and ingression fails entirely when the partially redundant adhesion protein SAX-7 is simultaneously knocked down (Grana et al., 2010).

Beyond its signaling role, HMR-1 also serves as the mechanical anchor for force transmission. The HMR-1/catenin complex anchors to the actomyosin cortex, physically coupling adjacent cells and enabling force transmission (Borghi et al., 2012; Clarke and Martin). The strength of this anchoring is modulated by additional proteins that bind to this complex, referred to as a ‘molecular clutch’ (Elosegui-Artola et al., 2016; Fortunato and Sunyer, 2022; Blanchard et al., 2018). During Ea/Ep ingression, this clutch regulates the coupling between the actomyosin cortex and cell-cell junctions. Live-embryo microscopy showed that, initially, the actomyosin network flows centripetally (toward the center) across the apical surface while junctions remain stationary, producing cortical slippage (Slabodnick et al., 2023). As the molecular clutch engages, junctions and cortex move in concert, and ZYX-1 is a mediator in this mechanical coupling (Slabodnick et al., 2023).

Several molecular drivers of gastrulation have been identified, but the biomechanics of the process have not been addressed in a quantitative, systems-level mechanical analysis. Key questions remain: What forces are generated during ingression, and is apical constriction alone sufficient to drive the observed movements? How do concurrent processes such as cell divisions contribute mechanically? Could divisions in the AB lineage facilitate ingression by promoting tissue unjamming, a transition from jammed to fluid-like behavior that reduces mechanical resistance (Tarannum et al., 2022; Mao and Wickström, 2024)? Computational models have provided insight into gastrulation biomechanics (Monier et al., 2015; Gracia et al., 2019; Yamashita et al., 2025; Dries et al., 2025). However, most existing models focus on epithelial sheet bending, such as ventral furrow invagination in *Drosophila*, and are often restricted to 2D representations. In contrast, *C. elegans* gastrulation involves individual cells ingressing into a 3D aggregate confined by a rigid eggshell, requiring models that explicitly represent individual cell mechanics, division dynamics, and spatial constraints.

In this work, we analyze the ingression of the endodermal precursor cells Ea and Ep from their birth to first division, when ingression is complete, through direct observation and mechanical modeling. This analysis is based on a dataset of 50 segmented 3D time-lapse recordings of wild-type embryos (section 5.1), including detailed 3D cell geometries and, for a subset, cortical localization of HMR-1, myosin, and F-actin. This dataset allows us to infer the embryo’s mechanical state. In particular, the angles at triple junctions, where three cells meet, provide a readout of the force balance at cell interfaces under the assumption of mechanical quasi-static equilibrium (Brodland et al., 2010; Vanslambrouck et al., 2024).

We complement this experimental analysis with biomechanical simulations based on a validated force model, the Deformable Cell Model (DCM) (section 5.2.2). This framework is an implementation of the Discrete Element Method (DEM), which represents the system as an assembly of interacting particles (Smeets et al., 2015). In the ‘active bubble’ formulation, each cell acts as a shell filled with fluid and with active surface tension (Smeets et al., 2019). The rigid eggshell is also modeled and imposes physical confinement. We initialize simulations with measured 3D cell shapes, division timings, and orientations from 17 embryos to capture in vivo context and natural variability. We incorporate active forces from cell division (Cuvelier et al., 2023), and explicitly model apical constriction in Ea and Ep to capture tension generation and force transmission. This integrated experimental and computational approach enables us to quantify, for the first time, the force distributions during *C. elegans* gastrulation and to dissect the mechanical contributions of apical constriction, cell division, and embryo-level rearrangements to Ea/Ep ingression.

## 3 Results

### 3.1 A timeline of Ea-Ep ingression

In *C. elegans*, gastrulation begins at the 26-cell stage with the ingression of the endodermal precursor cells Ea and Ep. In this paper, we analyzed the ∼45 minute interval spanning their lifetime (fig. 1A). These cells have a characteristic, extended cell cycle compared to the 22-minute cycle of their sisters MSa and MSp (Sulston et al., 1983), with Ep dividing on average 2.6 minutes later than Ea (fig. A14).

**Figure 1.**
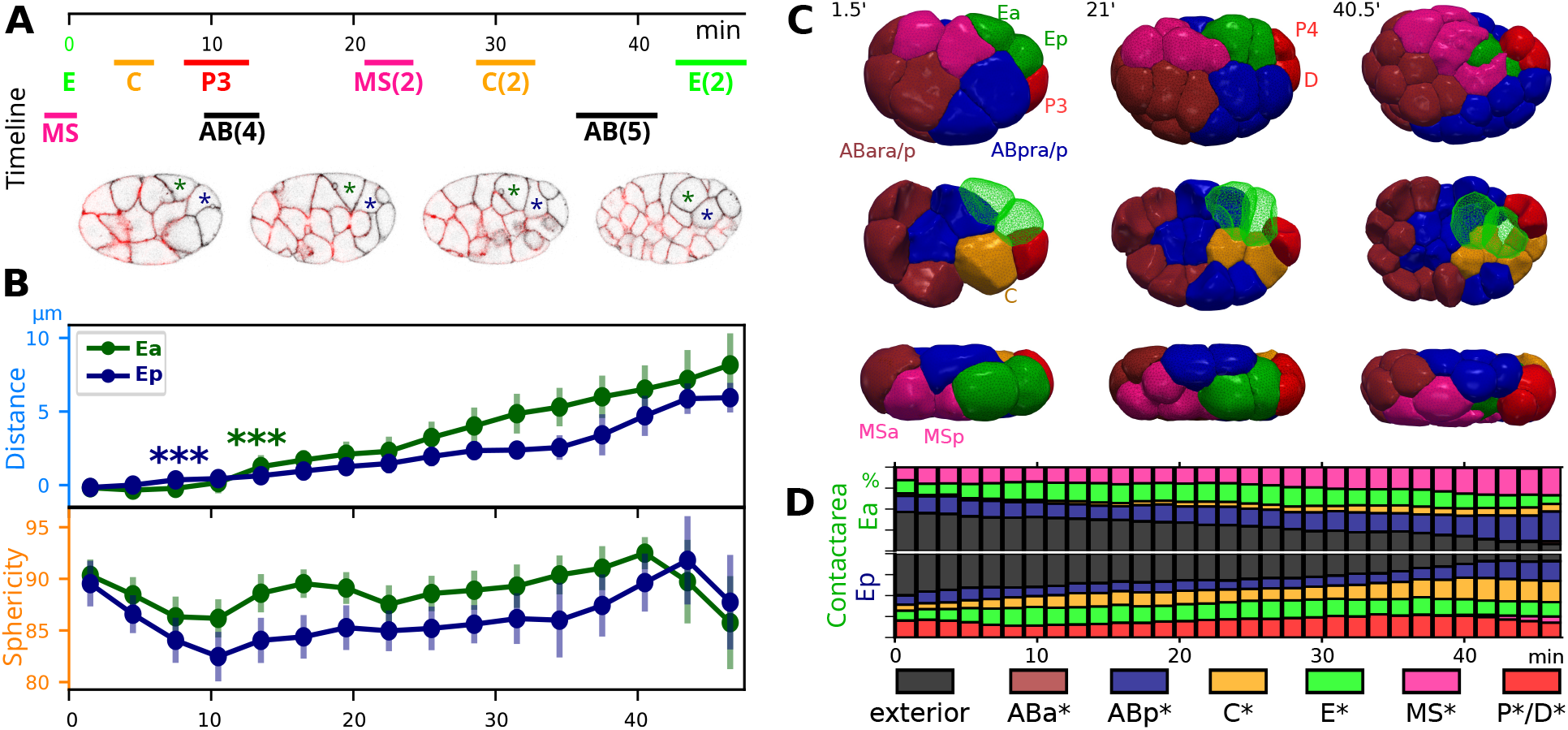
**(A)** Timeline (min) from E cell division. The horizontal bars represent the mean division times ± 1SD for a given cell or within a cell division round (round brackets) . Time is normalized by Ep lifetime (mean = 46.8 min). (strain RJ013, appendix A.1, actin in red, membrane in black). Asterisks highlight the ingressing Ea (left) and Ep (right) cell. **(B)** Top panel: Mean ingression distance [*µ*m] measured along the E-axis (appendix B.2). (*** indicates p *<* 0.001, t-test) indicates the first time bin where the mean ingression distance is significantly greater than zero (7.5 min for Ep, 13.5 min for Ea). Bottom panel: Mean sphericity over time. For both panels, error bars are ±1 SD (*n* = 26). **(C)** Detailed spatial context, showing segmented cells from replicate ‘crm04’. Top row: view from the right. Middle row: same view with the outer layer of cells removed to show the cells on the left. Bottom row: side view. **(D)** Contact areas of Ea and Ep during their lifetime. The free exterior area of both cells decreases in a fairly linear fashion over time, while the contact area between Ea and Ep remains largely constant throughout ingression.

Ingression of Ea and Ep, characterized by inward movement and reduced free surface area, begins shortly after division of their progenitor, the E cell, at 9.0± 5.1 minutes (mean ± standard deviation, SD) for Ep and 10.2 ±3.8 minutes for Ea (fig. 1B,fig. A14). At the time of their division, Ea and Ep often still have some remaining apical surface area, indicating that the covering by neighboring cells is not yet complete (fig. A13).

Ea ingresses further into the embryo, reaching a mean depth of 7.4±2.1 µm (fig. A14) along the ingression axis (appendix B.2), whereas Ep ingresses 6.0±1.1 µm. Their speed profiles also slightly differ: Ea maintains a relatively constant ingression speed throughout, while Ep temporarily plateaus around 25 minutes before accelerating at around 35 minutes (fig. 1B, fig. A13). Despite differences in depth and speed, Ea and Ep maintain a constant contact area throughout gastrulation, and appear to ingress as a mechanically coupled cell pair (fig. 1D). Both cells gradually become more spherical as they move inwards, a process that initiates approximately 20 minutes after their birth and continues until division (fig. 1B, fig. A13). They initiate rounding early in their extended cell cycle, whereas their sister cells, MSa and MSp, begin to round at the same time, but divide shortly after rounding (fig. 1A).

Several cell divisions punctuate this 45-minute interval (fig. 1A, fig. A5). Two synchronized AB lineage divisions occur during this time: the fourth, producing 16 ABxxxx daughter cells, at approximately 11 minutes after E-division, and the fifth at around 39 minutes. Division rounds are indicated with round brackets in this paper (e.g., AB(4)). Additional divisions include those of the C cell shortly before AB(4), P3 towards the end of AB(4), MS(2) around 22 minutes, and C(2) around 31 minutes (for the spatial context, see fig. 1C).

### 3.2 Role of apical constriction in ingression

Several observations point to a build-up of apical cortical tension in Ea and Ep. First, we observe dynamic membrane blebbing on their apical surfaces (fig. 2A, arrowhead, table A2), a phenomenon consistent with high cortical tension (Tinevez et al., 2009) that is not seen on the MS daughter cells. Blebbing is also prominent on the P4 cell as it moves over Ep (fig. 2B) and may contribute to its rapid movement (Pohlet al., 2012; Charras and Paluch, 2008) (fig. A11). Second, the geometry at triple junctions, which reflects the balance of cortical tensions, shows a distinct inflection point around 9 minutes post E-division, coinciding with the onset of ingression (fig. 2C.1). Apical constriction is the established driver of ingression for these endodermal precursor cells (Goldstein and Nance, 2020), but earlier studies placed its onset a few minutes after the MS(2) division (Sullivan-Brown et al., 2016; Marston et al., 2016). This timing (∼25 min. after the E-division) is considerably later than the ∼9 minutes onset we observe.

**Figure 2.**
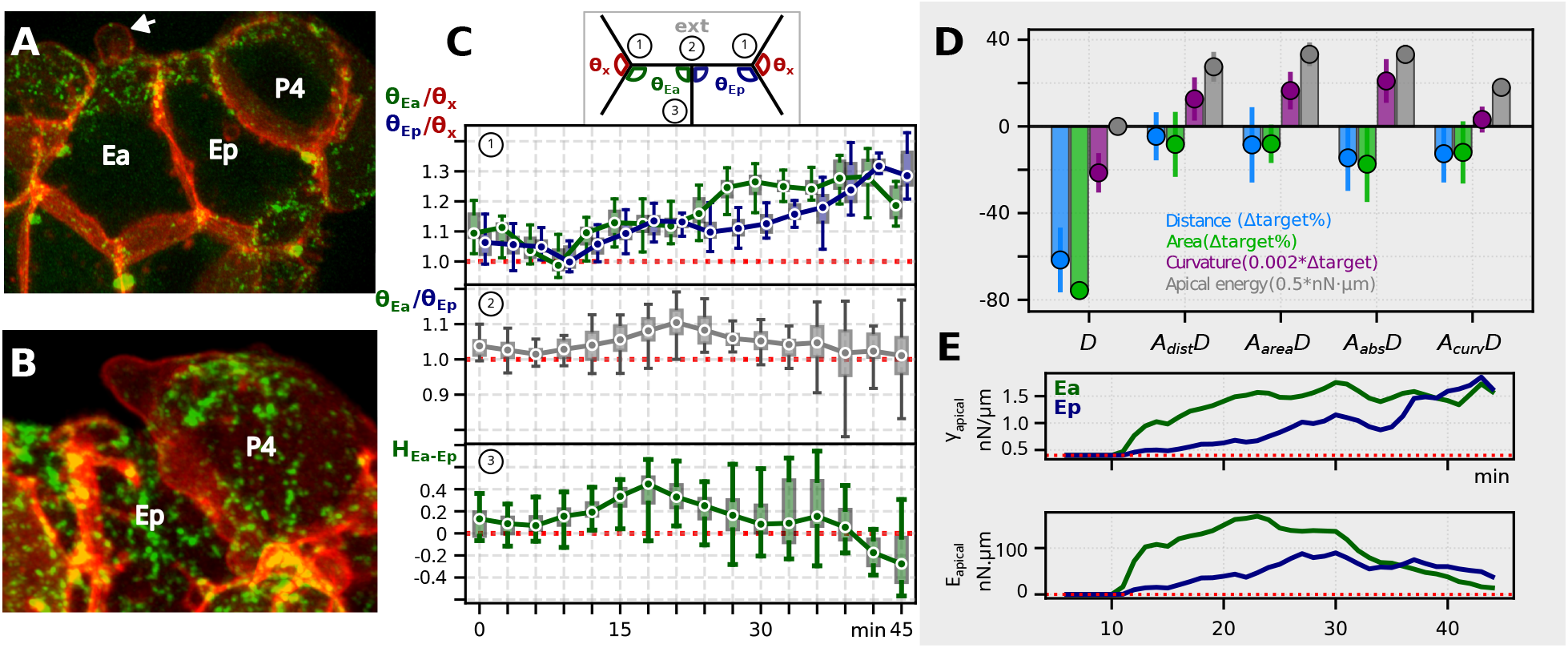
Apical constriction during E-cell ingression. **(A)** Z-maximum projection of ingressing Ea and Ep (strain RJ013, appendix A.1, myosin II in green and membrane in red). The image shows the even apical distribution of myosin, and an arrow indicates a membrane bleb on the apical surface. **(B)** Z-max projection of P4 cells showing pronounced membrane blebbing (arrow). **(C)** Temporal evolution of apical trijunction angles and interface curvature. 1. Apical trijunction angle ratio between E-cells and their neighbors (*θ*_*Ea*_*/θ*_*x*_ and *θ*_*Ep*_*/θ*_*x*_). 2. Apical trijunction angle ratio at the Ea-Ep interface (*θ*_*Ea*_*/θ*_*Ep*_). 3. Curvature of the Ea-Ep interface (*H*_*Ea−Ep*_*/H*_0_, where *H*_0_ = 10^5^). A positive value indicates a convex interface where Ea pushes into Ep. **(D)** Simulated ingression metrics (final ingression distance, final apical area, average E-cell curvature, average apical energy) for varying apical tension models. Y-axis: Deviation from target (ΔTarget). For ingression distance and apical area this deviation is expressed as a percentage. For average curvature, *Y* = (⟨*H*_*sim*_−*H*_*target*_⟩)×0.002[*m*^*−*1^] For apical energy, *Y* = ⟨*E*_*apical*_⟩×0.5[*nN* ·*µm*]. More details in Appendix appendix B.3.6. Scenarios: *D* (no apical tension), *A*_*dist*_*D* (apical tension tracks ingression distance), *A*_*area*_*D* (apical tension tracks apical area), *A*_*abs*_*D* (absolute tension), *A*_*curv*_*D* (apical tension tracks E-curvature). Matching ingression distance or area leads to excessive E-cell convexity, while matching curvature reduces ingression distance and area. Error bars: SD (n=17 replicates). **(E)** Apical tension and apical energy over the course of simulation, for Ea and Ep for the *A*_*dist*_*D* scenario (n=17). Lines show the mean value per one-minute time bin.

To characterize the mechanism of this early apical tension buildup, we analyzed F-actin and myosin distribution on the apical surfaces of Ea and Ep. Both F-actin and myosin are relatively uniform with no significant radial asymmetry (fig. A25). This property is captured by the moment of inertia metric, *prot*_*moi*_ (see appendix B.5.2), which indicates whether protein is concentrated centrally or dispersed peripherally. This distribution aligns with the architecture proposed by Zhang et al. (2023), where contractile forces are generated by a diffuse medioapical actomyosin network across the entire surface, rather than being confined to the periphery or center (fig. B7).

We next evaluated the trijunction angle ratio between E-cells and their neighbors to examine how the tensions in Ea and Ep change. First, the apical angle ratio indicates a lower tension in Ep compared to Ea between 21 and 36 minutes (fig. 2C.1). The small size of the neighboring P cell may imply a higher cortical tension, which could confound this measurement and lead to an underestimation of the tension in Ep. However, excluding P from analysis confirms lower Ep tension (fig. A16).

Second, the angle ratio at the apical Ea-Ep trijunction shows a wider angle for Ea, which is consistent with higher tension at the Ea apical interface compared to Ep, with the largest difference observed at 21 minutes (fig. 2C.2). Finally, from approximately 8 minutes post-division, Ea bulges into Ep, which reflects a greater internal pressure in Ea, with the maximal difference observed between 18 and 21 minutes (fig. 2C.3). Taken together, these observations suggest that while Ea and Ep ingress as a mechanically coupled pair, they exhibit different tension profiles, with Ea generating greater apical tension throughout most of the ingression, although Ep’s tension increases notably in the last 9 minutes.

We next used mechanical simulations to test whether apical constriction can account for the observed cellular movements (section 5.2). Simulations with a constant apical tension (*A*_*abs*_*D*) show that this mechanism can drive a substantial degree of ingression (fig. 2D), but this often results in an unbalanced ingression where one cell progresses significantly more than its partner (fig. A54a). To better reproduce the observed dynamics, we then explored more complex scenarios where the applied tension was varied over time. Of these, the scenario that dynamically adjusted tension to track the observed ingression distance (*A*_*dist*_*D*) reproduced the in vivo dynamics more closely and also mitigated this imbalance (fig. A52), while requiring slightly less apical energy compared to a constant tension approach (fig. 2D). The simulation’s inferred tension profile is broadly consistent with our in-vivo analysis, showing a higher apical tension for Ea between 10 and 30 minutes, and a late boost in apical tension for Ep in the last stage of ingression (fig. 2E). We return to this ingression imbalance when we discuss the role of apical HMR-1 accumulation at the Ea-Ep interface. Overall, our simulations confirm that apical constriction is sufficient to drive ingression. However, the overly convex shape of the E-cells in simulations that match the target ingression distance suggests that the required apical tension is higher than expected from the in-vivo situation (fig. 2D).

### 3.3 Differential effects of cell divisions

During Ea-Ep ingression, we observe that the timing of two major synchronized division rounds, AB(4) and AB(5) (fig. A5), correlates with peaks in the inward volumetric flux of the E-cells (fig. 3A). Since cell divisions can generate strong mechanical forces that drive substantial cell rearrangements in embryos (Atia et al., 2021), we used our simulation framework to examine how divisions influence ingression.

**Figure 3.**
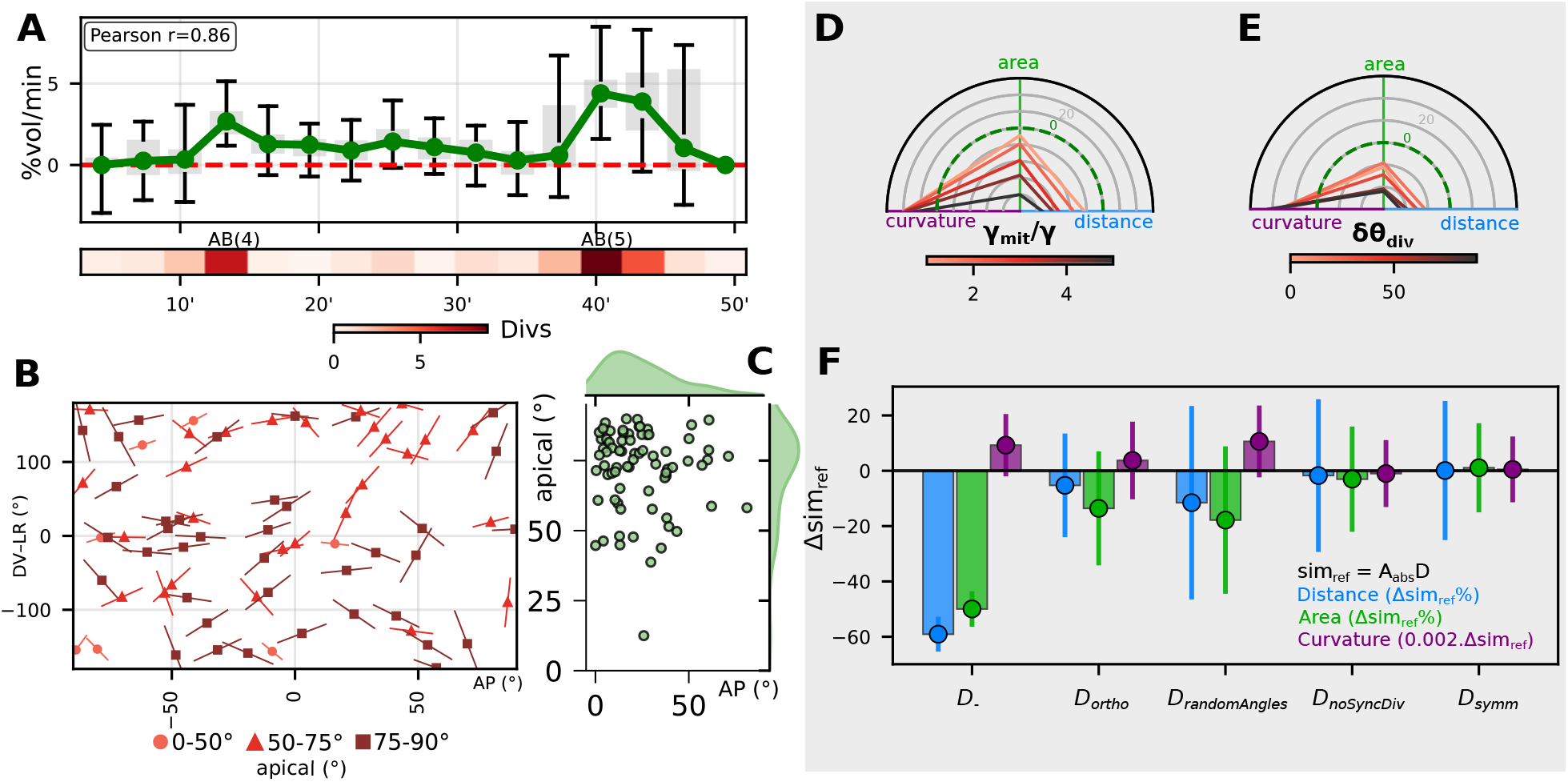
Cell division during E-cell ingression. **A.** Correlation between cell division events and E-cell ingression, quantified by the volumetric flux (%vol/min, see appendix B.3.7) of Ea/p into the embryonic interior. The bar plot, subdivided into 3-minute intervals, shows an increase in volume flux concomitant with the AB(4) and AB(5) division rounds. The color bar indicates the average number of cell divisions within each 3-minute interval. (Boxplot every 3 minute interval, every box represents IQR between Q1 and Q3, whiskers at min and max values) **B**. Division angles for all cell divisions occurring during E-ingression. The apical angle (degrees) represents the orientation relative to the eggshell normal (see appendix B.4.1), where 0° indicates a division orthogonal to the eggshell, towards the interior. Most divisions occur parallel to the eggshell and are oriented along the AP direction (see fig. A6 for more detail). **C**. Distribution of division angles (apical angle, see B, versus angle to the AP axis) for all divisions during E-cell ingression, with each point representing the average for a specific dividing cell. **D**. Simulated effect of mitotic tension (*γ*_*mit*_) on ingression metrics. *γ*_*mit*_ represents the surface tension during mitosis, relative to interphase surface tension (*γ*). The default simulation condition is *γ*_*mit*_ = 3*γ*. (Metrics as in fig. 2E, n=5, constant apical tension of 1.2 nN/*µ*m). **E**. Simulated effect of rotating the division angle in the AP-DV plane. The sweep applies incremental rotations (18°, 36°, 54°, 72°, and 90°) relative to the original division angles. Increasing deviation from the original angles leads to a worsening of ingression metrics. (n=17, constant apical tension of 1.2 nN/*µ*m) **F**. Impact of different simulated cell division scenarios on ingression, shown as Δ relative to the reference scenario *A*_*abs*_*D*. Scenarios include: no division (*D*_*−*_), 90° shift in division angle in the AP-DV plane (*D*_*ortho*_), randomized division angles (*D*_*randomAngles*_), unsynchronized AB divisions (*D*_*noSyncDiv*_, see fig. A41), and volume-symmetric divisions (*D*_*symm*_). (Metrics as in fig. 2E, n=17, constant apical tension of 1.2 nN/*µ*m)

We simulated different scenarios in which the occurrence, orientation, mitotic tension, and synchronicity of cell divisions varied, while the applied apical tension on Ea and Ep was kept constant (fig. 3F, fig. A37). First, completely removing cell divisions from simulations (*D_*), leading to fewer and larger cells, severely hampers ingression, requiring significantly more apical tension to drive the process (fig. A38). This shows that smaller cells are easier to displace during cell rearrangements. Next, we assessed the role of division orientation. Cell divisions during gastrulation are highly stereotyped, i.e. they consistently follow the same orientation with little variability (fig. A6, fig. A7). Divisions predominantly occur parallel to the eggshell surface (fig. 3C), with their axes aligned in the anteroposterior direction (fig. 3B). Simulations in which division angles were altered, either by randomization (*D*_*randomAngles*_) or progressive rotation, hampered ingression (fig. 3E, fig. A39), which indicates that the stereotyped orientation contributes functionally to gastrulation. We then examined other division properties, such as symmetry and synchronicity. In contrast to the critical role of orientation, simulations with equalized daughter cell volumes (*D*_*symm*_) or with unsynchronized AB divisions (*D*_*noSyncDiv*_) showed no significant or systematic effect on ingression (fig. 3F, fig. A41). Finally, we investigated the effect of forces generated during mitosis itself. In our simulations, a default cortical tension during mitosis was set to three times the base interphase tension, a value consistent with the 2-4 fold increase in cortical stiffness observed in mitotic mammalian cells (Taubenberger et al., 2020). Increasing this mitotic tension in simulations progressively worsened ingression metrics (fig. 3D, fig. A40).

These results indicate that cell divisions facilitate ingression mainly by producing smaller cells, which can reposition more easily than larger cells during internalization. The stereotyped orientation of these divisions also contributes, whereas mechanical perturbations from mitotic rounding have a counteracting effect.

### 3.4 The onset of apical constriction is marked by E-cadherin polarization

Initially, the E cell is born with low E-cadherin (HMR-1) levels, as cortical flows during the asymmetric EMS division move most HMR-1 to the MS sister cell (Caroti et al., 2021). This E–MS asymmetry persists into its daughters, Ea and Ep(fig. A19, fig. A21, fig. A20, fig. A23). Subsequently, coinciding with the onset of apical constriction at around 9 minutes post E-division (section 3.2), we observe the formation of a distinct apical HMR-1 enrichment at the Ea-Ep interface (fig. A28, fig. A27).

As discussed in the introduction, HMR-1 reduces basolateral contractility, while the HMR-1-free apical surface recruits MRCK-1, driving higher myosin activity. Such a contractility gradient can drive cortical flows. Marston et al. (2016) previously observed the apical HMR-1 accumulation at the Ea-Ep interface and found it to be myosin-dependent, consistent with such a flow. We therefore hypothesized that a basal-to-apical cortical flow drives this apical HMR-1 enrichment. Spatio-temporal measurements show an apical shift of HMR-1 (fig. 4B), confirmed by a polarization metric (fig. 4C), and detailed microscopy observations support the existence of such a flow at the Ea-Ep interface (fig. 4A). The same pattern of cadherin relocation was not observed at the interfaces with neighboring cells (fig. A29). These observations support a myosin-driven cortical flow as the mechanism of apical HMR-1 enrichment (fig. 6.1-3).

**Figure 4.**
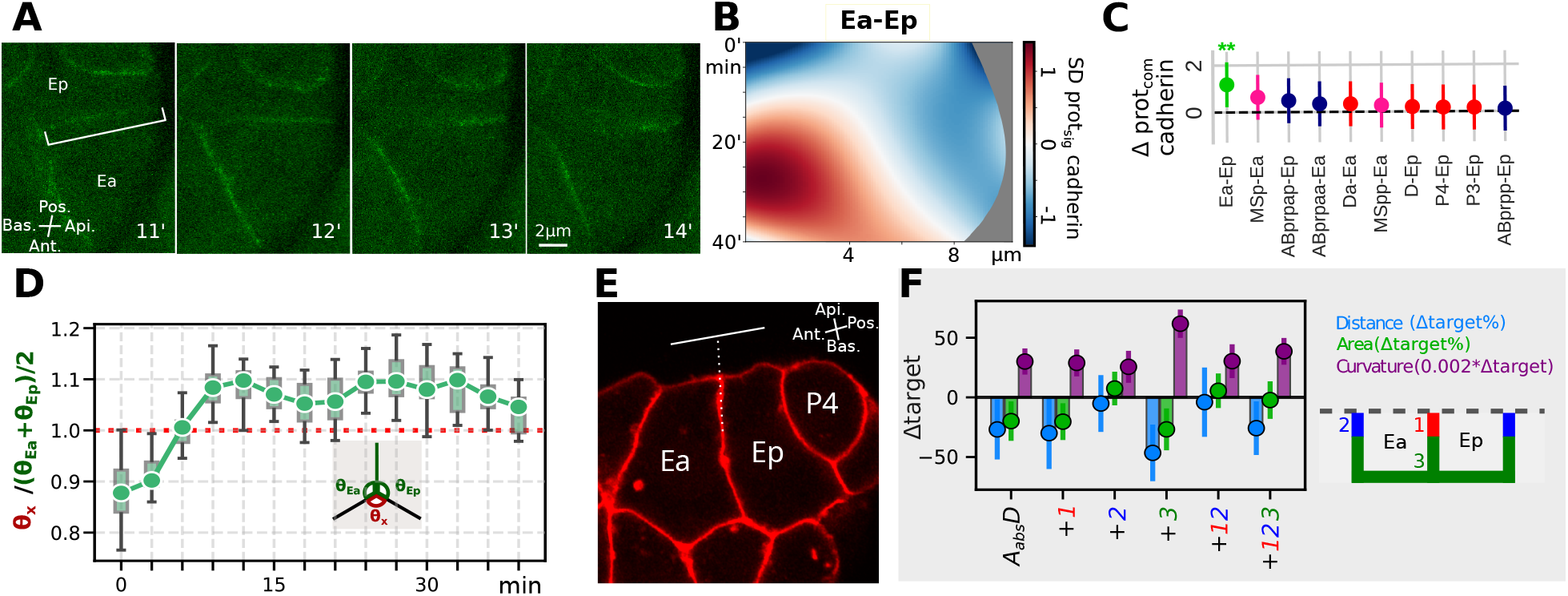
**(A)** Left: Maximum intensity Z-projection of endogenously tagged E-cadherin (strain RJ014, appendix A.1), illustrating its apical shift at the Ea-Ep interface. **(B)** Kymographs showing the spatio-temporal distribution of E-cadherin at the Ea-Ep interface. The y-axis represents time from E-division, and the x-axis indicates distance from the apical side. E-cadherin shows a progressive apical accumulation, initiating around 15 minutes post E-division. **(C)** Statistical analysis of cadherin asymmetry (*prot*_*com*_) at E-cell interfaces (see appendix A.9.8 for details). The plot shows the time-averaged deviation (Δ), which compares *prot*_*com*_ at an interface contacting an E-cell with the mean value across all other non-apical and non-basal interfaces of that same cell. The top 10 interfaces with the most significant enrichment are shown. This analysis reveals a significant asymmetric build-up of cadherin exclusively at the Ea-Ep interface, a pattern not observed at interfaces with neighboring cells. **(D)** Temporal evolution of triple junction angles involving Ea and Ep. The plot shows the ratio *θ*_*x*_*/*((*θ*_*Ea*_ + *θ*_*Ep*_)*/*2), where *θ*_*x*_ is the angle spanned by the non-E cell. From 9 minutes post E-division, this ratio is significantly greater than one, consistent with the proposition that the Ea-Ep interface experiences reduced tension compared to surrounding interfaces. **(E)** Zoomed in membrane image (strain RJ013, appendix A.1, membrane in red). ∼20 minutes before E(2) division. Apical surface angles approach 90 degrees as indicated by the white lines. **(F)** Simulated ingression metrics evaluating the effect of increasing cell-cell friction at specific regions of the E-cells (see appendix A.11.7 for details). Three regions are tested: at the apical Ea-Ep interface (marked as +1), at the apical ring contacting neighboring cells (marked as +2), and at the basolateral region (marked as +3). Metrics show the deviation (Δ) from the target, with the reference simulation (*A*_*abs*_*D*) included for comparison (n=17, constant apical tension of 1.2 nN/*µ*m).

As the apical cadherin accumulation forms, around 9 minutes post E-division, the trijunction angles at the basolateral side of the Ea and Ep contact indicate a drop in the relative interfacial tension (fig. 4D). This suggests that the higher apical tension of Ea and Ep is not transmitted across this interface. This tension drop is also observed at the apical side, where the angles between Ea and Ep and their interface approaches 90 degrees, and the two apical surfaces align (fig. 4E, fig. A1). The local tension drop is consistent with HMR-1’s known role in reducing cortical contractility (Klompstra et al., 2015; Marston et al., 2016).

We next evaluate a potential mechanical role of the apical HMR-1 accumulation using our simulation framework. In simulation, the HMR-1 accumulation is represented as a local increase in cell-cell friction at the apical Ea-Ep interface (fig. 4F, scenario ‘+1’). This addition had little effect on ingression dynamics or distance, but led to a more complete reduction of the apical free area (fig. 4F, fig. A50). Further, in simulations with constant apical tension (*A*_*abs*_*D*), the ingression imbalance between Ea and Ep was reduced (fig. A52,fig. A56). In the distance-tracking scenario (*A*_*dist*_*D*), apical tensions of Ea and Ep rose more linearly and in tandem (fig. A55). This simpler, more synchronized behavior is consistent with the expectation that the sister cells operate in the same way and have comparable force profiles. The simulations suggest that the apical HMR-1 accumulation acts as a mechanical anchor, coupling the cell pair and mediating force transmission, which facilitates their synchronized ingression.

### 3.5 A frictional clutch transmits apical constriction forces

How ingression proceeds depends on the physical properties of the interfaces between the E-cells and their neighbors. Our simulations show that differential adhesion, increasing the adhesion strength of the E-cells (fig. A47, fig. A49), can facilitate ingression, and effectively promotes their internalization (fig. A44). However, this mechanism requires an enrichment of adhesion molecules at the E-cell interfaces with their neighbors, which is not supported by our protein distribution data (fig. 4F, fig. A24, fig. A29). While we observed a trend toward HMR-1 apical polarization at the MSp-Ea interface (fig. 4C), consistent with observations by Marston et al. (2016), this was not statistically significant in our analysis and was absent at other neighbor contacts.

Next, we consider how the apical constriction forces generated by the E-cells are transmitted to pull neighboring cells over them. This requires adhesion proteins connecting the cells to each other, and to the actomyosin cortex. Slabodnick et al. (2023) proposed that force transmission is regulated during apical constriction through a frictional clutch that strengthens cadherin-actin linkage (see introduction). We used our simulations to test the spatial effects of such a clutch, modeled as a drastic localized increase (1000-fold) in cell-cell friction. Increasing friction at the apical ring, where the E-cells contact their neighbors, facilitated ingression, with a modest increase in ingression distance and a larger decrease in the final apical free area (fig. 4F, scenario +2), regardless of changes to global cell-cell friction (fig. A53). Conversely, increasing friction at the basolateral region resists downward movement and severely hinders ingression (scenario +3). Together, our findings support an apical-ring coupling mechanism that transmits constriction forces to neighboring cells. Since we observe no significant cadherin enrichment at these interfaces, this coupling is likely established by the recruitment of clutch-mediating proteins to the high-tension apical ring as described by Slabodnick et al. (2023). However, Slabodnick et al. (2023) characterize the early stage of gastrulation (3–7 minutes post-MSxx birth) as an ‘ineffective’ phase of cortical slippage where actomyosin is uncoupled from junctions. In contrast, our observation of early apical force generation (section 3.2) suggests that significant friction is already being sustained and transferred during this initial period.

### 3.6 Embryo-wide flow during AB(5)

During the AB(5) division round, coordinated embryo-wide cell movements are apparent. To characterize these, we reconstructed 3D velocity fields from the cells’ centroid trajectories of all embryos using Stokes flow (see appendix A.12). In this approach, the cell collective is modeled as a viscous medium with free-slip boundary conditions along the eggshell. This aggregate analysis shows a distinct rotational pattern, or curl, emerging specifically during the late phase of ingression (fig. A58). We found no evidence of additional active force sources beyond the AB(5) division round. Notably, this curl is absent in our simulations (fig. A58). We attribute this discrepancy to overdamping in the DCM, where local dissipation at cell-cell interfaces limits long-range force transmission. This interpretation is further supported by the slightly elevated E-cell curvature in our simulations noted above (fig. 2D, *A*_*dist*_*D*), which suggests higher than expected apical overpressure even when ingression distance is matched.

### 3.7 Actin-rich protrusions with localized E-cadherin mediate the closure

At the end of ingression, neighboring cells converge to cover the gastrulation site, forming rosette-like configurations (fig. 5A,B), a recurring motif in *C. elegans* gastrulation whose stability and organization depend on cadherin-based adhesion (Serre et al., 2023). Cells of the ABp and C lineage on the dorsal side are repositioned (fig. A8, fig. A12), while on the ventral side, neighboring cells converge to cover the gastrulation cleft.

**Figure 5.**
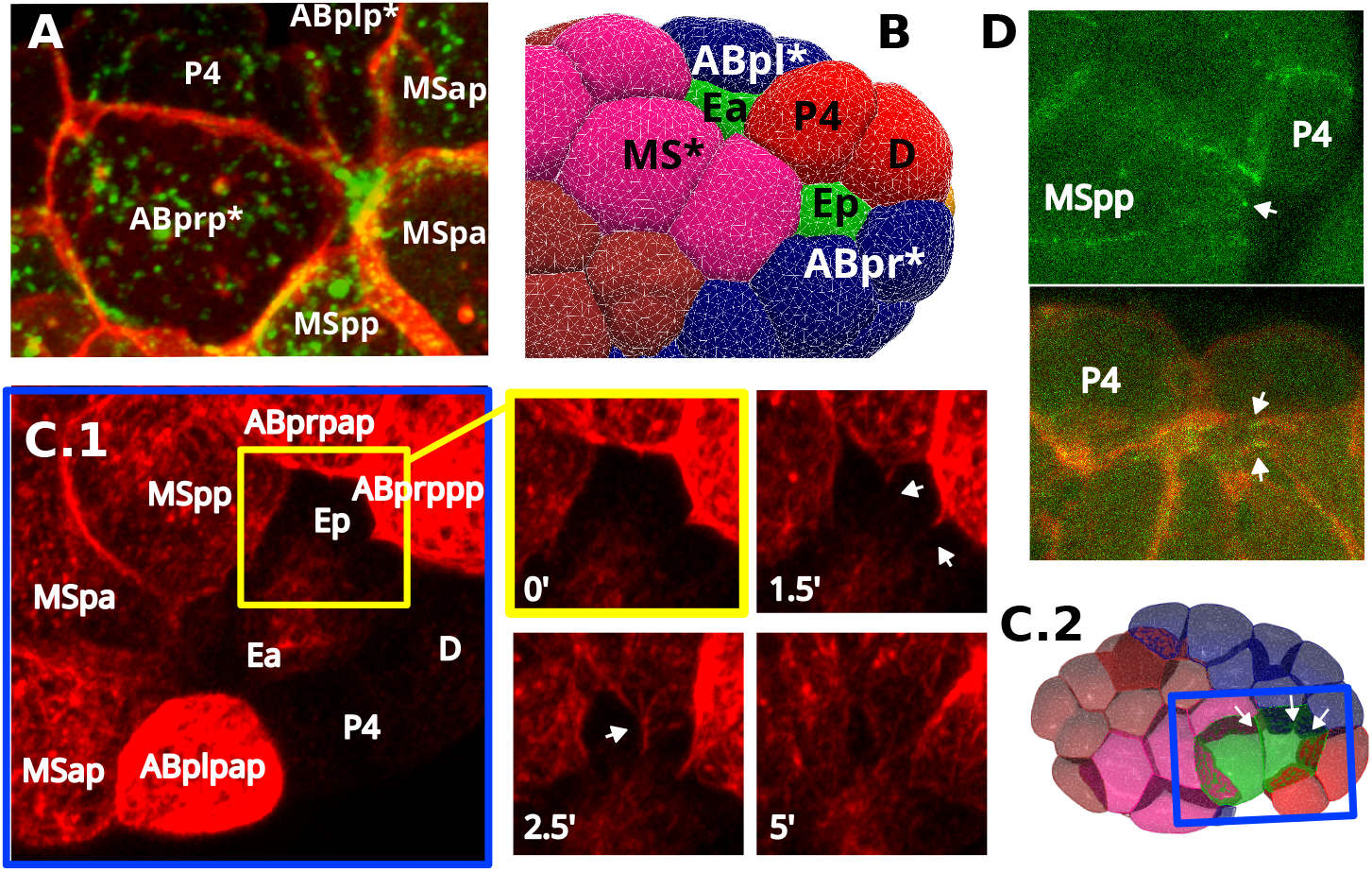
Covering cells at the end of E-cell ingression. (**A**) Maximum intensity Z-projection showing the rosette-like arrangement of cells covering the ingressing endodermal precursor cells (strain RJ013, appendix A.1, myosin in green and membranes in red). (**B**) Covering cells (right view) form rosette configurations around Ea (P4, MSpp, MSpa, MSap, ABplpap) and around Ep (P4, D, ABprppp, ABprpap, MSpp). (**C**) 1. Maximum intensity Z-projection of F-actin in covering cells during the final stages of rosette closure (strain RJ012, appendix A.1). The yellow boxed inset provides a temporal zoom illustrating the formation of transient, linear actin protrusions (white arrows) extending from the leading edges of converging cells, including ABprpap, MSpp, and ABprppp. t=0 matches approximately 44.5 min post-E division. 2. Spatial context for the blue boxed inset. (**D**) Maximum intensity Z-projection showing E-cadherin enrichment at the tip of covering cells (strain RJ004, appendix A.1, E-cadherin in green, membrane in red).

In the final stages of closure, distinct actin protrusions extend from the leading edges of covering cells toward the closing center (fig. 5C). These structures are consistent with the “dynamic filopodia” observed by Pohl et al. (2012), although they appear in our observations as transient extensions persisting for several minutes (fig. 5C.1) rather than on a sub-minute timescale.

These late-stage, filopodial structures are distinct from the earlier, lamellipodia-like actin enrichments reported by Roh-Johnson and Goldstein (2009), which are restricted to the MS-lineage and appear much earlier (22-28 minutes post E-division). We further observe enrichment of E-cadherin at the tips of these protrusions, beginning around 38 minutes post E-division (fig. 5D). In other morphogenetic systems, cadherin enrichment at protrusion tips establishes localized adhesive anchor points for cell migration and tissue remodeling (Adams et al., 1996; Almagro et al., 2010). The actin protrusions, with HMR-1 enrichment at their tips, indicate localized, directed force generation. We found no evidence in our images of a supracellular actin ring (fig. A22, fig. A30), making a purse-string closure, driven by such a ring as in embryonic wound healing (Garcia-Fernandez et al., 2009), unlikely (Pohl et al., 2012).

This late, protrusion-driven sealing mechanism is not represented in our simulations, which apply apical tension only to the free apical surface of the E-cells.

## 4 Discussion

In this paper, we combined quantitative microscopy with computational modeling to capture the mechanical processes of early gastrulation in *C. elegans*. Our results establish a multi-stage mechanical program in which the ingressing E-cells, their direct neighbors, and other cells throughout the embryo each contribute to successful ingression (fig. 6).

**Figure 6.**
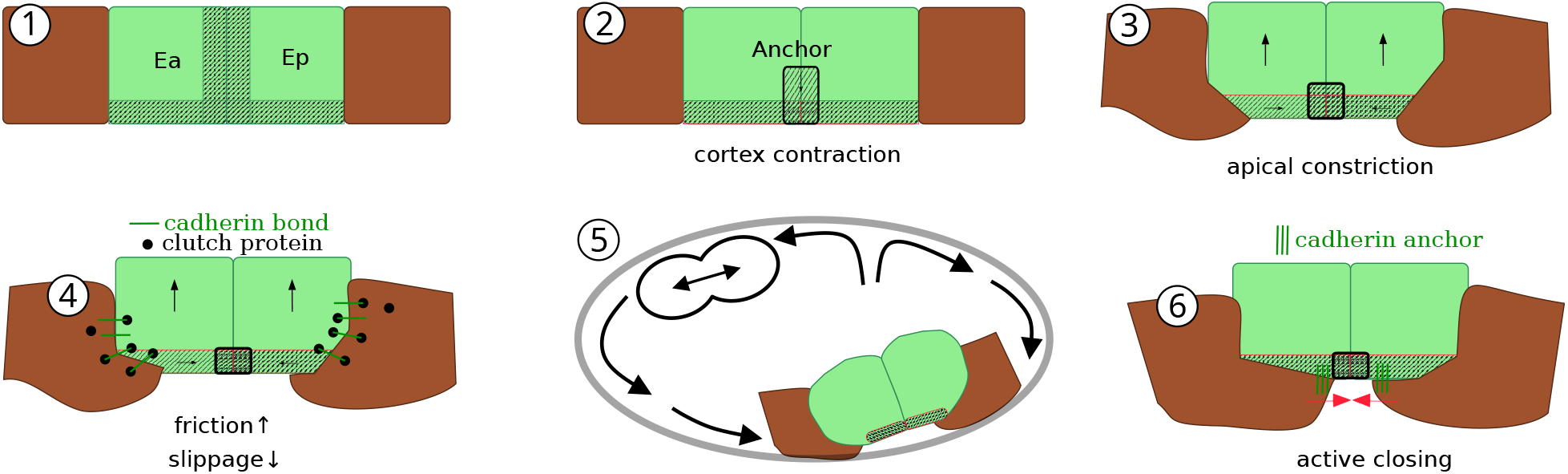
A schematic model of the proposed phases in Ea-Ep ingression. (1) Initial state. (*<*∼9 min post E-division): No cortical activation present at the point. (2) Anchor formation (∼9 min): Cortical contractions drive a basal-to-apical flow, creating a E-cadherin enrichment at the apical side of the Ea-Ep interface that functions as a mechanical anchor. (3) Start apical constriction (>∼9 min): With the anchor in place, the medioapical actomyosin network can generate tension, leading to constriction of the apical surface, which initiates the inward ingression of the cell pair. (4) Ingression and covering phase (∼9-40 min): As ingression proceeds, a molecular clutch mechanism increases cell-cell friction at the apical ring where E-cells contact their neighbors, transmitting the apical constriction forces to the neighboring cells and reducing cortical slippage. (5) Ingression is simultaneously aided by a passive, embryo-wide flow, which is generated by the combined active forces (apical contractions and cell divisions) exerted on cells within the confined space of the embryo. (6) Active closure (>40 min): In the final phase, the covering cells actively seal the gastrulation cleft by extending transient, actin-rich protrusions. E-cadherin enrichment at the tips of these protrusions creates localized anchor points, allowing the cells to pull themselves over the ingressing E-cells.

First, approximately 9 minutes after E-division, at the onset of apical constriction, a cortical flow along the Ea-Ep interface generates an apical enrichment of HMR-1 (fig. 6.2, fig. 4A,B). We propose this flow is driven by a basal-to-apical contractility gradient resulting from apical myosin activation. This is consistent with the myosin- and catenin-dependence of HMR-1 enrichment as reported by Marston et al. (2016), as it provides both the motive force (myosin) and the mechanical linkers (catenins) necessary to transport HMR-1 along the moving cortex.

Our simulations indicate that apical constriction is sufficient to drive ingression. Further, they suggest the HMR-1 enrichment acts as a mechanical anchor that enables tension buildup at the apical surface, which drives apical constriction and ingression (fig. 6.3). The anchor also couples the two sister cells at the apical side, a mechanical link that transmits forces and stabilizes their synchronized ingression.

Force transmission from the E-cells is spatially modulated: while our simulations show that locally increased friction at the apical ring facilitates ingression, friction at other locations, including the basolateral region, impedes it (fig. 6.4). Since we observe that ingression and apical tension generation begins at approximately 9 minutes post E-division, force transmission must already be active well before the late-stage ZYX-1-mediated clutch engagement described by Slabodnick et al. (2023).

Beyond the E-cells and their direct neighbors, ingression coincides with embryo-wide cell division rounds, most notably AB(4) and AB(5), which correlate with peaks in E-cell inward flux (fig. 6.5). Our simulations show that these divisions facilitate ingression by producing smaller, more numerous cells that are more easily displaced, and by their conserved orientation parallel to the eggshell. During AB(5), we observe a distinct embryo-wide rotational cell flow. We consider this flow a passive reorientation induced by a division-driven transition to a more fluid-like collective cell state (Mao and Wickström, 2024).

During final closure, covering cells extend actin-rich protrusions with cadherin-enriched tips (fig. 6.6, fig. 5B,D), consistent with an active sealing mechanism driven by localized adhesion and force generation (Serre et al., 2023; Zilberman et al., 2017).

Our biomechanical model captures the core mechanics of ingression but several modeling limitations remain. First, the model overestimates local force dissipation, a known limitation of the DCM formulation. This prevents reproduction of the embryo-wide rotational flow observed during AB(5). Direct integration of a Computational Fluid Dynamics (CFD) solver into the simulation framework would provide a more faithful representation of long-range force transmission. Second, cadherin is treated as a friction modifier, but biologically we propose it is a prerequisite for force transmission. Future work could model cadherin localization explicitly, for example conditioned on cortical flow. Third, the actomyosin cortex is abstracted as a homogeneous shell with static friction parameters, which does not capture dynamic processes such as cortical turnover or tension-dependent mechanofeedback, for example the strengthening of cadherin-actin linkages under load (Arslan et al., 2021). Fourth, active protrusion-driven closure is not modeled.

In summary, we show that apical constriction is sufficient to drive Ea-Ep ingression, but that the process depends on a broader mechanical program. A cortical flow-driven cadherin anchor couples the cell pair and enables force transmission at the onset of constriction. Spatially localized friction at the apical ring transmits constriction forces to neighboring cells, while directed divisions in the AB lineage facilitate ingression by reducing mechanical resistance. Together, these results provide a quantitative, integrated account of local and embryo-wide mechanical contributions to *C. elegans* gastrulation.

## 5 Methods and materials

### 5.1 Dataset

This study is based on a total of 50 replicate datasets (see appendix A.1, table A1). Of these, 38 capture the Ea and Ep gastrulation phase: 26 cover the process completely, while 12 cover it partially. For mechanical simulations, 17 of these replicates were utilized. Additionally, specific molecular markers were examined: 8 replicates were enriched for E-cadherin, 7 for NMYII, and 7 for F-actin. Cell segmentation from the time-lapse microscopy data followed the methodology outlined by Thiels et al. (2021) (appendix A.2.2). Following segmentation, a range of features were determined for each cell, including volume (fig. A2), sphericity (fig. A3), division timing (fig. A5), cell lifetime (fig. A4), trajectories (fig. A10), cell velocities (fig. A11, cortical protein content (fig. A19) and distributions (fig. A24).

### 5.2 A mechanical framework for modeling ingression

#### 5.2.1 Simulation setup

In this work, we employ mechanical simulations to explore the apical constriction mechanism during *C. elegans* gastrulation and investigate other potentially important factors (fig. 7). We utilized segmented 3D time-lapse recordings from 17 wild-type replicate embryos, focusing on the gastrulation of endodermal precursor cells (Ea and Ep), which serve as the ‘target’ or ground-truth data to initialize, guide, and evaluate the simulations (table A1). Simulations were initialized using the 3D segmented cell configurations from a timepoint after E-cell division but before the AB4 division round. A rigid eggshell boundary, modeled as a repulsively interacting convex hull with negligible friction, was introduced (appendix C). The simulation period spanned approximately 37 minutes, starting just before the AB4 division and concluding 1.5 minutes before the Ea division. This timeframe captures the complete ingression movements of both Ea and Ep cells, but excludes their subsequent divisions and the final closure of the gastrulation cleft, which occurs post-division. The primary metrics used to evaluate ingression in simulations, compared to the target data, are the final ingression distance, the final apical area, and the average curvatures of the E-cells (appendix B.3).

**Figure 7.**
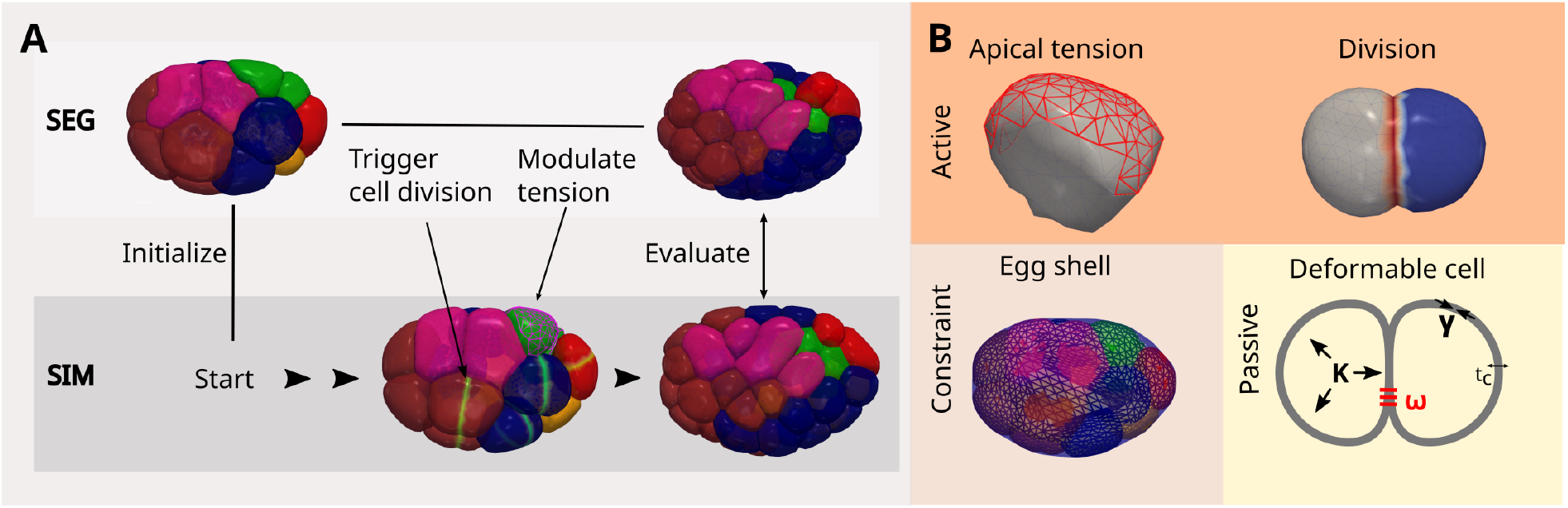
**A.**Schematic illustrating the mechanical simulation setup. Simulations (SIM) are initialized using segmented 3D data (SEG) from a specific timepoint (post-E-division, pre-AB(4) division round) from one of 17 replicate embryos (label ‘crm’, see table A1). During the simulations, cell divisions occur according to ground-truth segmentation data, accurately matching timing, division orientation, and volume asymmetry. Apical tension on the E cells is either constant, or modulated by tracking metrics such as ingression distance or free apical area. The simulation outcomes are evaluated by comparison to ground-truth using metrics including final ingression distance, final apical area, average E-cell curvature, and centroid positioning of all embryonic cells. **B**. Overview of the force model employed in mechanical simulations (appendix C). Active forces include the application of apical tension to the E cells and the forces generated by cell divisions. Mechanical constraints include the rigid convex hull approximating the egg shell. Passive forces are described by the Deformable Cell model, where the cell cortex is represented as a viscous thin layer (thickness *t*_*c*_) under active tension (*γ*). Given that the timescale of gastrulation significantly exceeds the cortical viscoelastic relaxation time (*τ*_*M*_ ≈ 5 s) (Saha et al., 2016), elastic effects are considered negligible.. Interfacial tension between cells is modeled as the sum of their cortical tensions minus adhesion energy (*ω*), with cytoplasmic pressure accounted for by a volumetric modulus (*K*).

#### 5.2.2 Model rationale and parameterization

Our simulation framework is built upon the active foam model implemented in the Mpacts software. This particle-based, deformable cell model, which represents cells as detailed 3D meshes, has been extensively validated in previous studies (Smeets et al., 2015, 2019; Cuvelier et al., 2021, 2023). It is particularly well-suited for simulating cell aggregates with irregular and varied cell shapes.

The parameters for this base model were chosen by starting with values established in the literature (appendix C.1.1). Key parameters such as surface tension, cell-cell adhesion, and effective cortex thickness were adopted from previous work (Cuvelier et al., 2023).

For friction parameters, where experimental data is scarce, we used the hydrodynamic length scale *L*_*h*_ as a guide. Prior literature utilizing laser ablation indicates a regime where cortical viscosity dominates stress propagation, characterized by *L*_*h*_ ≈ 14 *µ*m (Saha et al., 2016). We adopted *η* = 11 kPa·s, *t* = 300 nm, and *ξ* = 0.12 kPa·s·*µ*m^*−*1^ (appendix C.1.1), yielding 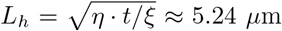, comparable to the average cell radius in our model (*R*_0_ ≈ 6.9 *µ*m). We further confirmed that scaling of cortical viscosity, medium viscosity, and cell-cell friction in tandem, does not alter ingression metrics (fig. A45). The specific influence of modifying cell-cell friction will be further explored in fig. 4.

To this passive mechanical model, we added active forces. First, cell division was incorporated using the model from Cuvelier et al. (2023) (appendix C.1.2). Second, to model apical constriction, we applied a uniform tension across the apical surface of the E-cells (appendix C.1.3). This simple approach is justified by two key observations: the distribution of myosin on the apical surface appears uniform (section 3.2), and myosin activation is understood to occur in regions lacking E-cadherin, which would exclude cell-cell interfaces. This is also consistent with our findings that the Ea-Ep interface, in particular, exhibits low tension (section 3.4). Therefore, a uniform increase in tension across the free apical surface represents the most parsimonious yet biologically plausible implementation.

## 6 Acknowledgments

This work was supported by FWO aspirant grants to W.T., M.V., C.v.B. and J.V.: [11I2921N, 1194222N, 11L0923N, 11D9923N], and FWO research grant G008423N and KU Leuven internal funds grant [C14/24/109] to R.J..

## Supplementary materials

The appendices are available as separate PDF documents: Appendix A (general), Appendix B (definitions), and Appendix C (force models).

