## Appendix A for "Integrated quantitative imaging and biomechanical modeling of early gastrulation in *C. elegans*"

### A Appendix

#### A.1 C. elegans strains and maintenance

*C. elegans* strains were cultured on nematode growth medium (NGM) plates at 20°C as described by Brenner (1974).

- **OD70**: Genotype `ltIs44 [pie-1p::mCherry::PH(PLC1δ1) + unc-119(+)]`. This strain expresses a fluorescent marker (`mCherry::PH(PLC1δ1)`) under the control of the `pie-1` promoter, which predominantly labels the cell membrane.
- **RJ004**: Genotype `hmr-1(cp21[hmr-1::GFP + LoxP])I; unc-119(ed3)III; ltIs44V`. This strain combines a cadherin marker (`HMR-1::GFP`) with the membrane marker from OD70 (`mCherry::PH(PLC1δ1)`).
- **RJ012**: A cross of `LP162 (nmy-2(cp13[nmy-2::GFP + LoxP])I)` and `SWG001 (gesIs001[Pmex-5::Lifeact::mKate2::nmy-2UTR + unc-119(+)])`. This strain expresses a myosin marker (`NMY-2::GFP`) and an actin-associated marker (`Lifeact::mKate2`).
- **RJ013**: A cross between `LP306 (cpIs53[mex-5p::GFP-C1::PH(PLCδ)::tbb-2 3'UTR + unc-119(+)]II)` and `SWG001 (gesIs001[Pmex-5::Lifeact::mKate2::nmy-2UTR + unc-119(+)])` (Reymann et al., 2016). This strain combines a membrane marker (`GFP-C1::PH(PLCδ)`) with an actin-associated marker (`Lifeact::mKate2`).

Additionally, we used microscopy data from Cao et al. (2020). These replicates are labeled as "crm".

Live embryos were mounted on slides with M9 buffer and compressed using Polybead Microspheres (20 µm, Polysciences) as spacers.

Embryos were imaged on either a Zeiss LSM 880 confocal microscope or a Nikon A1 confocal microscope, using Plan-Apochromat 60x or 63x oil immersion objectives. Z-stacks were acquired with a spacing of 1–1.2 µm, an *xy* pixel resolution ranging from 0.0977 to 0.1295 µm, and a temporal resolution of 1.5 to 3 minutes per stack.

#### A.2 Data

##### A.2.1 Overview replicates

table A1.

Table A1: Summary of replicates. 3D+t time lapses of these replicates were used in segmentation and subsequent quantitative analysis. 'crm' replicates specifically were used to run the simulations. Microscopy data capturing zoomed-in images (instead of full-embryo images) are not listed here.

| replicate | strain | sim | gast | t E interval | t EMS interval | #t | res t | res z | res xy | protein | #cells start | #cells end |
|---|---|---|---|---|---|---|---|---|---|---|---|---|
| crm04 | Cao et al. 2020 | True | True | [0.0, 48.0] | NaN | 33 | 90.0 | 0.2 | 0.2 | NaN | 13 | 49 |
| crm05 | Cao et al. 2020 | True | True | [0.0, 48.0] | NaN | 33 | 90.0 | 0.2 | 0.2 | NaN | 13 | 51 |
| crm06 | Cao et al. 2020 | True | True | [0.0, 48.0] | NaN | 33 | 90.0 | 0.2 | 0.2 | NaN | 13 | 47 |
| crm07 | Cao et al. 2020 | True | True | [0.0, 48.0] | NaN | 33 | 90.0 | 0.2 | 0.2 | NaN | 13 | 48 |
| crm08 | Cao et al. 2020 | True | True | [0.0, 48.0] | NaN | 33 | 90.0 | 0.2 | 0.2 | NaN | 13 | 46 |
| crm09 | Cao et al. 2020 | True | True | [0.0, 48.0] | NaN | 33 | 90.0 | 0.2 | 0.2 | NaN | 12 | 46 |
| crm10 | Cao et al. 2020 | True | True | [0.0, 48.0] | NaN | 33 | 90.0 | 0.2 | 0.2 | NaN | 12 | 47 |
| crm11 | Cao et al. 2020 | True | True | [0.0, 48.0] | NaN | 33 | 90.0 | 0.2 | 0.2 | NaN | 13 | 47 |
| crm12 | Cao et al. 2020 | True | True | [0.0, 48.0] | NaN | 33 | 90.0 | 0.2 | 0.2 | NaN | 12 | 47 |
| crm13 | Cao et al. 2020 | True | True | [0.0, 48.0] | NaN | 33 | 90.0 | 0.2 | 0.2 | NaN | 5 | 47 |
| crm14 | Cao et al. 2020 | True | True | [0.0, 48.0] | NaN | 33 | 90.0 | 0.2 | 0.2 | NaN | 12 | 47 |
| crm15 | Cao et al. 2020 | True | True | [0.0, 48.0] | NaN | 33 | 90.0 | 0.2 | 0.2 | NaN | 12 | 46 |
| crm16 | Cao et al. 2020 | True | True | [0.0, 48.0] | NaN | 33 | 90.0 | 0.2 | 0.2 | NaN | 12 | 47 |
| crm17 | Cao et al. 2020 | True | True | [0.0, 55.5] | NaN | 38 | 90.0 | 0.2 | 0.2 | NaN | 12 | 49 |
| crm18 | Cao et al. 2020 | True | True | [0.0, 48.0] | NaN | 33 | 90.0 | 0.2 | 0.2 | NaN | 13 | 47 |
| crm19 | Cao et al. 2020 | True | True | [0.0, 48.0] | NaN | 33 | 90.0 | 0.2 | 0.2 | NaN | 12 | 46 |
| crm20 | Cao et al. 2020 | True | True | [0.0, 48.0] | NaN | 33 | 90.0 | 0.2 | 0.2 | NaN | 13 | 47 |
| 7cell01 | OD70 | False | False | NaN | [3.0, 15.0] | 5 | 90.0 | 1.1 | 0.1 | NaN | 7 | 8 |
| 7cell02 | OD70 | False | False | NaN | [3.0, 15.0] | 5 | 90.0 | 1.0 | 0.1 | NaN | 7 | 8 |
| 7cell03 | OD70 | False | False | NaN | [3.0, 15.0] | 5 | 90.0 | 1.0 | 0.1 | NaN | 7 | 8 |
| 7cell04 | OD70 | False | False | NaN | [1.5, 13.5] | 5 | 90.0 | 1.0 | 0.1 | NaN | 7 | 8 |
| 7cell05 | OD70 | False | False | NaN | [1.5, 13.5] | 5 | 90.0 | 1.0 | 0.1 | NaN | 7 | 8 |
| 7cell06 | OD70 | False | False | NaN | [1.5, 13.5] | 5 | 90.0 | 1.0 | 0.1 | NaN | 7 | 8 |
| 7cell07 | OD70 | False | False | NaN | [1.5, 13.5] | 5 | 90.0 | 1.0 | 0.1 | NaN | 7 | 9 |
| hmr01 | RJ004 | False | True | [-27.8, 53.6] | [-7.9, 73.4] | 42 | 119.8 | 0.8 | 0.2 | cadherin | 4 | 47 |

Continued on next page

| replicate | strain | sim | gast | t E interval | t EMS interval | #t | res t | res z | res xy | protein | #cells start | #cells end |
|---|---|---|---|---|---|---|---|---|---|---|---|---|
| hmr02 | RJ004 | False | True | [-43.6, 57.5] | [-23.8, 77.4] | 52 | 119.8 | 0.8 | 0.2 | cadherin | 2 | 46 |
| hmr03 | RJ004 | False | True | [-65.4, 55.5] | [-41.6, 79.3] | 62 | 119.8 | 0.8 | 0.2 | cadherin | 1 | 44 |
| hmr04 | RJ004 | False | True | [-20.0, 98.0] | [-2.0, 116.0] | 57 | 120.2 | 0.8 | 0.2 | cadherin | 5 | 65 |
| hmr05 | RJ004 | False | True | [-44.0, 74.0] | [-26.0, 92.0] | 60 | 120.2 | 0.8 | 0.2 | cadherin | 2 | 50 |
| hmr07 | RJ004 | False | True | [-20.0, 72.0] | [0.0, 92.0] | 47 | 120.2 | 0.8 | 0.2 | cadherin | 6 | 61 |
| hmr08 | RJ004 | False | True | [-46.0, 46.0] | [-26.0, 66.0] | 47 | 120.2 | 0.8 | 0.2 | cadherin | 1 | 39 |
| hmr09 | RJ004 | False | False | [32.8, 74.6] | [62.6, 104.4] | 11 | 179.7 | 0.8 | 0.1 | cadherin | 36 | 72 |
| wt01 | RJ013 | False | True | [-30.0, 81.0] | [-12.0, 99.0] | 38 | 180.0 | 0.5 | 0.1 | NaN | 4 | 83 |
| wt02 | RJ013 | False | False | NaN | [-28.5, 15.0] | 30 | 90.0 | 0.3 | 0.1 | NaN | 2 | 8 |
| wt03 | RJ013 | False | False | NaN | [69.0, 156.0] | 30 | 180.0 | 0.5 | 0.1 | NaN | 12 | 79 |
| wt04 | RJ013 | False | True | [-34.5, 9.0] | [-9.0, 34.5] | 30 | 90.0 | 0.3 | 0.1 | NaN | 4 | 15 |
| wt05 | RJ013 | False | False | NaN | [1.5, 27.0] | 18 | 90.0 | 1.1 | 0.1 | myosin | 7 | 12 |
| wt07 | RJ013 | False | True | [-52.5, 21.0] | [-27.0, 46.5] | 50 | 90.0 | 0.8 | 0.1 | actin | 2 | 24 |
| wt08 | RJ013 | False | True | [-57.0, 31.5] | [-31.5, 57.0] | 60 | 90.0 | 0.5 | 0.1 | actin | 2 | 26 |
| wt09 | RJ013 | False | True | [-34.5, 54.0] | [-13.5, 75.0] | 60 | 90.0 | 0.5 | 0.1 | actin | 4 | 47 |
| wt10 | RJ013 | False | True | [-46.5, 42.0] | [-21.0, 67.5] | 60 | 90.0 | 0.8 | 0.1 | actin | 3 | 30 |
| wt11 | RJ013 | False | True | [-31.2, 41.5] | [-13.4, 59.3] | 50 | 89.9 | 0.8 | 0.1 | actin | 4 | 44 |
| wt12 | RJ013 | False | True | [-44.5, 40.0] | [-23.7, 60.8] | 58 | 89.9 | 0.8 | 0.1 | actin | 2 | 39 |
| wt13 | RJ013 | False | True | [-56.7, 119.3] | [-29.8, 146.2] | 60 | 179.8 | 0.8 | 0.1 | actin | 2 | 108 |
| wt14 | RJ013 | False | True | [-28.2, 59.3] | [-10.4, 77.1] | 60 | 89.8 | 0.8 | 0.1 | myosin | 4 | 52 |
| wt15 | RJ013 | False | True | [1.5, 89.0] | [16.3, 103.8] | 60 | 89.8 | 0.8 | 0.1 | myosin | 14 | 87 |
| wt16 | RJ013 | False | True | [-50.4, 37.1] | [-28.2, 59.3] | 60 | 89.9 | 0.8 | 0.2 | myosin | 2 | 28 |
| wt17 | RJ013 | False | True | [-35.6, 51.9] | [-14.8, 72.7] | 60 | 89.9 | 0.8 | 0.2 | myosin | 4 | 47 |
| wt18 | RJ013 | False | True | [-48.0, 40.5] | [-27.0, 61.5] | 60 | 90.1 | 0.8 | 0.2 | myosin | 2 | 28 |
| wt19 | RJ013 | False | True | [-46.5, 42.0] | [-24.0, 64.5] | 60 | 90.1 | 0.8 | 0.2 | myosin | 2 | 42 |

##### A.2.2 Segmentation

Segmentation was performed using the spheresDT-MpacsPiCS software (Thiels et al., 2021). Lineage trees from spheresDT were manually curated and named (appendix A.14).

#### A.3 Microscopy

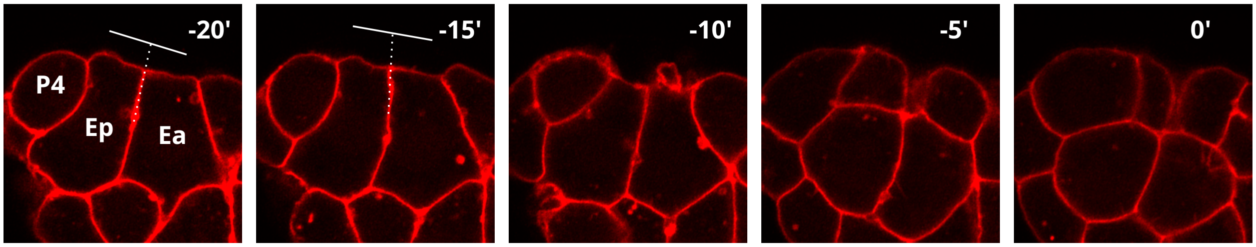


Figure A1: Zoomed in membrane image (strain RJ013, appendix A.1, membrane in red). Time indicates minutes until E(2) divisions. Apical surface angles approach 90 degrees as indicated by the white lines.

Table A2: Count of blebbing occurrences. 5 images were checked, all contained zoomed in images showing Ea and Ep and parts of the neighbors.

| Cell | Observed | Not Observed |
|---|---|---|
| Ea/Ep | 5 | 0 |
| P4 | 3 | 2 |
| MSxx | 1 | 4 |

#### A.4 Cell geometry

fig. A2

##### A.4.1 Volume

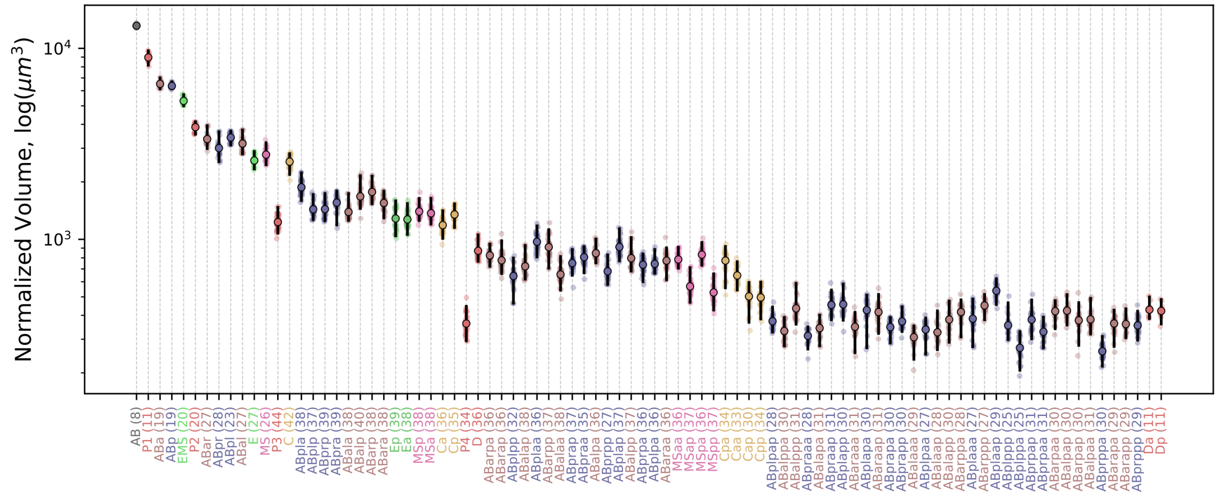


Figure A2: Each point represents the time-averaged volume for a single cell from one replicate, filtered for cells tracked for at least three timepoints. Number of data points indicated in brackets. Cell types are ordered by birth time along the x-axis. The central colored dot indicates the mean volume for each cell, while the vertical black line represents the 95% confidence interval. The volumes are plotted on a logarithmic scale in $\mu\mathrm{m}^3$.

##### A.4.2 Sphericity

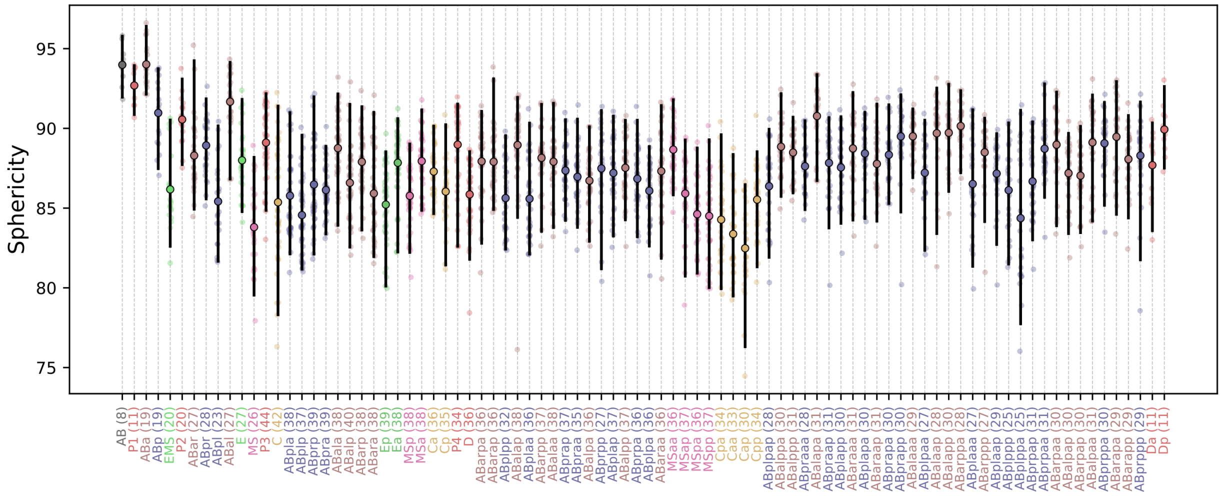


Figure A3: Each point represents the time-averaged sphericity for a single cell from one replicate, filtered for cells tracked for at least three timepoints. The number of data points for each cell is indicated in brackets. Cell types are ordered by birth time along the x-axis. The central colored dot indicates the mean sphericity, while the vertical black line represents the 95% confidence interval. Sphericity is a dimensionless ratio.

#### A.5 Cell lifetime

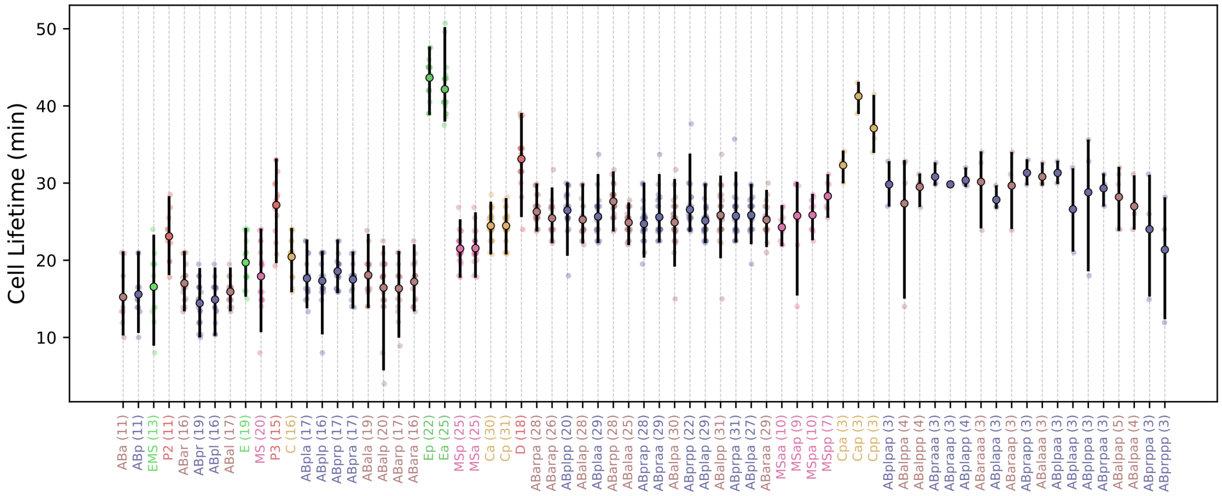


Figure A4: Cell lifetimes in the early *C. elegans* embryo. Each point represents the calculated lifetime for a single cell from one replicate. Outliers (lifetimes outside 3×IQR from the quartiles) were then identified and removed for each cell individually. The number of data points shown for each cell (after filtering) is indicated in brackets. Cell types are ordered by birth time along the x-axis. The central colored dot indicates the mean lifetime, and the vertical black line shows the 95% confidence interval. Lifetimes are measured in minutes (min).

#### A.6 Cell divisions

fig. A5

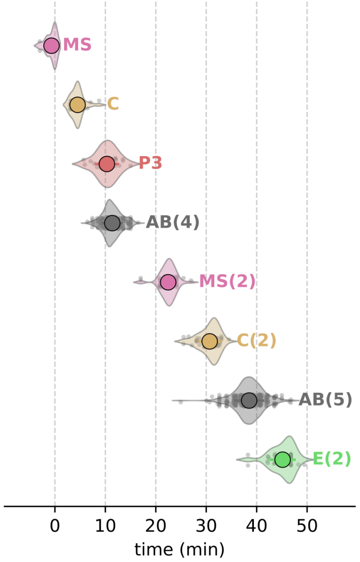


Figure A5: Timing of division rounds during Ea. X-axis is time from E division. Each transparant dot represent an individual cell division time. Word on notation: AB(4) is the fourth division round of AB, resulting in ABxxxx cells.

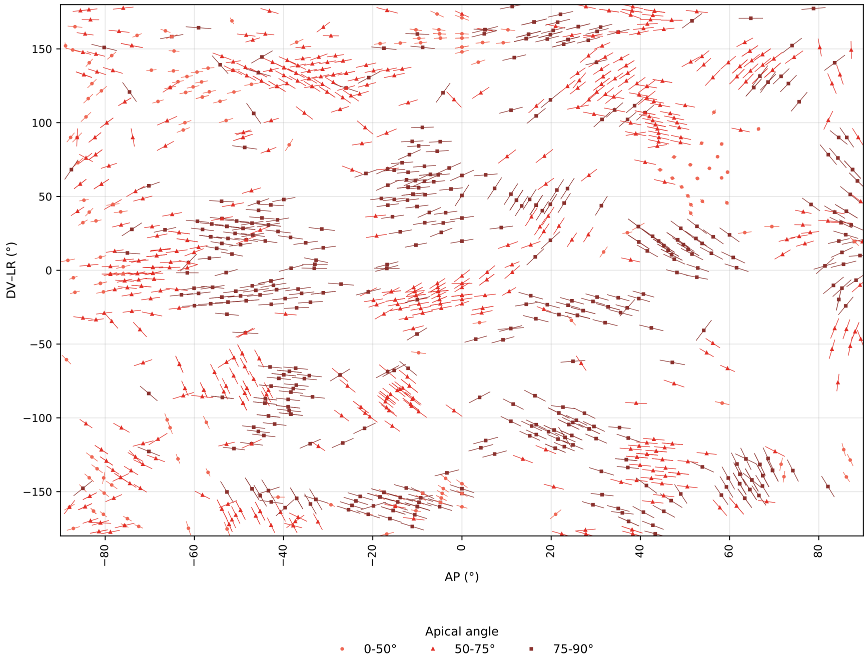


Figure A6: Division angles for all cell divisions occurring during E-ingression. This is the same figure as fig. 3B, but not averaged over replicates per cell, every datapoint is shown. The apical angle (degrees) represents the orientation relative to the eggshell normal (see appendix B.4.1), where 0° indicates a division orthogonal to the eggshell, towards the interior. Most divisions occur parallel to the eggshell and are oriented along the AP direction.

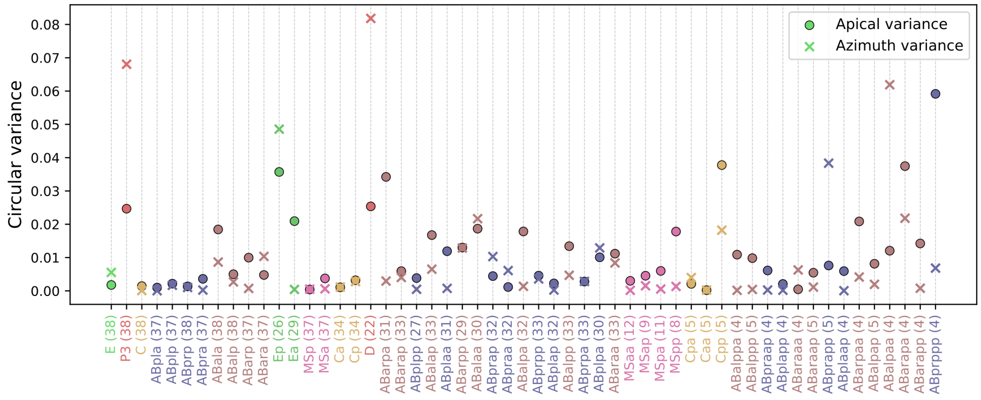


Figure A7: Circular variance of cell division orientations. The plot displays the variability for different cells, sorted by their time of birth. The number of data points for each cell is indicated in brackets. Apical variance (circles) quantifies the division's tilt relative to the embryo's surface, while azimuth variance (crosses) measures its 'compass direction' on the surface. Circular variance ($V$) is a measure of angular dispersion ranging from 0 to 1, where $V = 0$ indicates perfect consistency (all divisions have the identical orientation) and $V = 1$ indicates complete randomness. It is defined as $V = 1 - R$, where $R$ is the length of the mean division vector (unit vector) averaged across all replicates. To account for the undirected nature of the division axis, the 360° azimuth angles are folded into a 0–90° range before calculating the variance. This step prevents 180° ambiguities from artificially inflating the variance. The resulting low variance values for most cells, typically $V < 0.05$, demonstrate that division orientations are highly stereotypical and exhibit little variation.

#### A.7 Ingression characterization

##### A.7.1 Volume flux

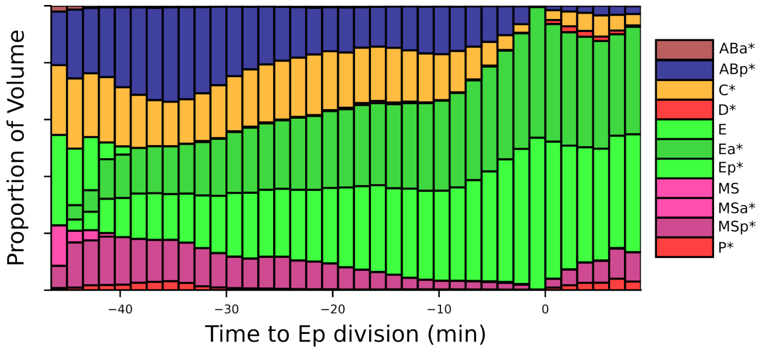


Figure A8: The volumetric composition of the space that will eventually be occupied by Ea and Ep over time. Time is aligned relative to the division of the Ep cell, which is set to $t = 0$. Incorporates data from crm-replicates (n=17).

##### A.7.2 Curvatures

fig. A9

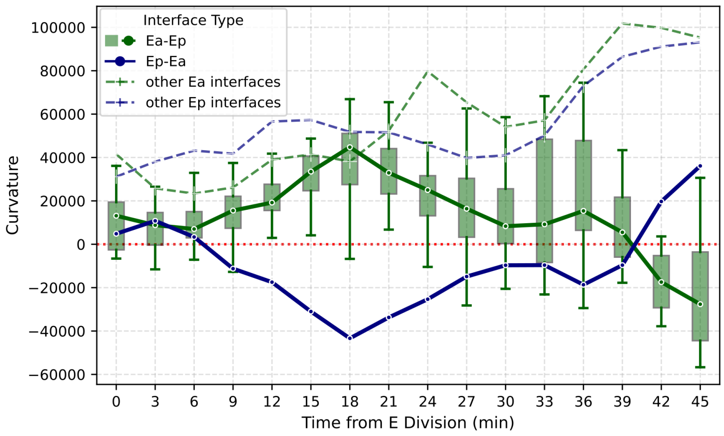


Figure A9: Curvature of the interfaces of Ea and Ep over time. The green boxplot represent the curvature of the Ea-Ep interface, boxed per 3 minute intervals, showing a slightly convex surface suggesting Ea to be pushing into Ep. For completeness the curvature of the Ep-Ea interface (in blue, full line) is also shown. The average curvature of the other interfaces of Ea and Ep are shown as a dotted line, showing both Ea and Ep to be rounding up over the course of ingression.

##### A.7.3 Cell trajectories

fig. A10

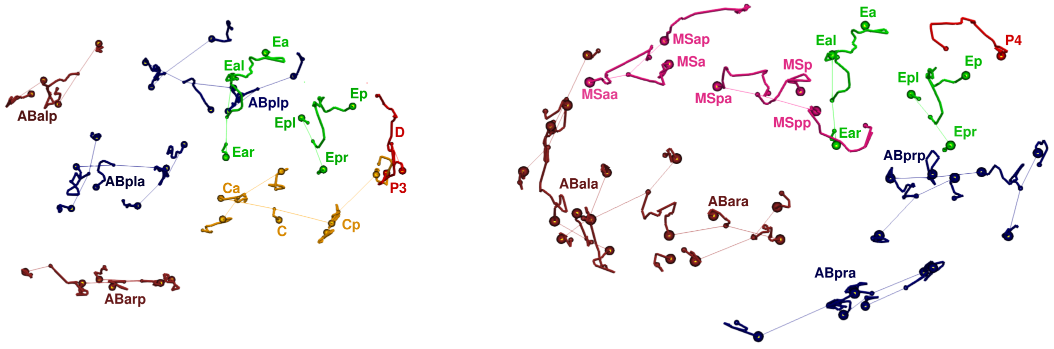


(a) Left side

(b) Right side

Figure A10: Replicate *crm04* showing the cell trajectories of all cells during Ea-Ep gastrulation viewed in the AP-DV plane. For visual clarity, only half of the cells are shown per panel. Cell trajectories are indicated with full lines. Mother-daughter connections are shown in thin lines. Cell trajectory starting points are shown with big spheres, end points are shown with small spheres.

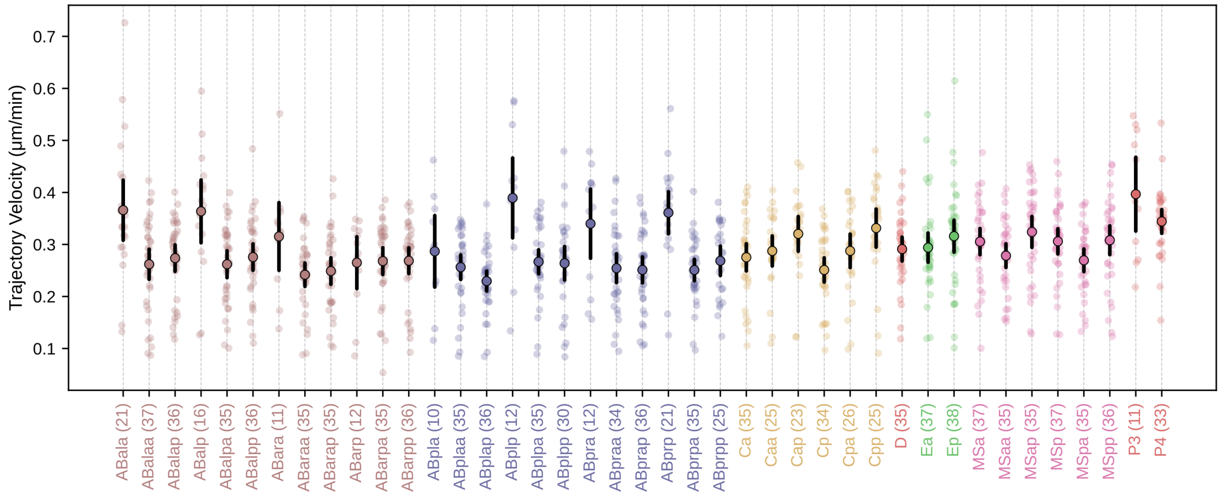


Figure A11: The trajectory velocity ($\mu$m/min) for cells calculated during the Ea/Ep gastrulation phase. Each colored dot represents the average velocity for a specific cell from a single embryo replicate. Number of data points indicated in brackets. The circle with the black outline indicates the mean velocity across all replicates, while the black vertical line represents the 95% confidence interval of the mean.

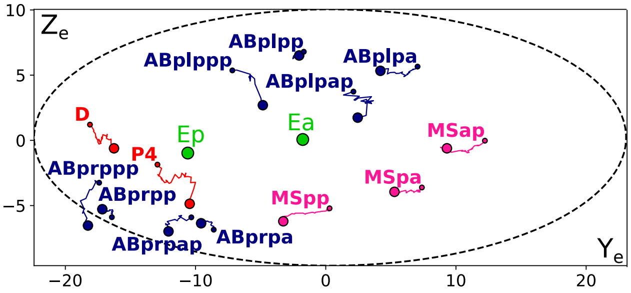


Figure A12: Trajectories of covering cell centroids. projected onto the $Y_E Z_E$ plane (the plane orthogonal to the ingression direction, appendix B.2). The dashed ellipse approximates the embryo boundary. These trajectories illustrate the collective convergence of neighboring cells towards the central ingression site.

##### A.7.4 Ingression metrics

fig. A13

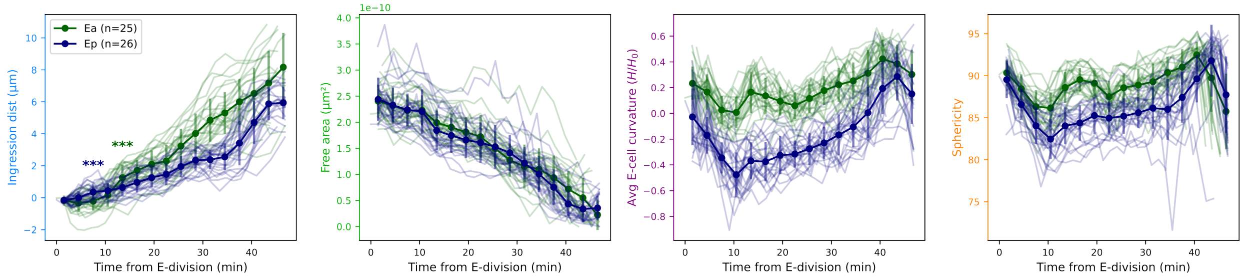


Figure A13: Ingression metrics: Ingression distance ($\mu$m), free apical area ($\mu m^2$), average E-cell curvature ($H/H_0$, $H_0 = 10^5$), and sphericity. Data is binned into 3-minute intervals, with binned means shown as markers connected by lines and vertical error lines representing 1SD. Only replicates spanning the full lifetimes of Ea or Ep are considered. A normalized time is used which scales Ep lifetime to the overall mean (46.8 minutes)

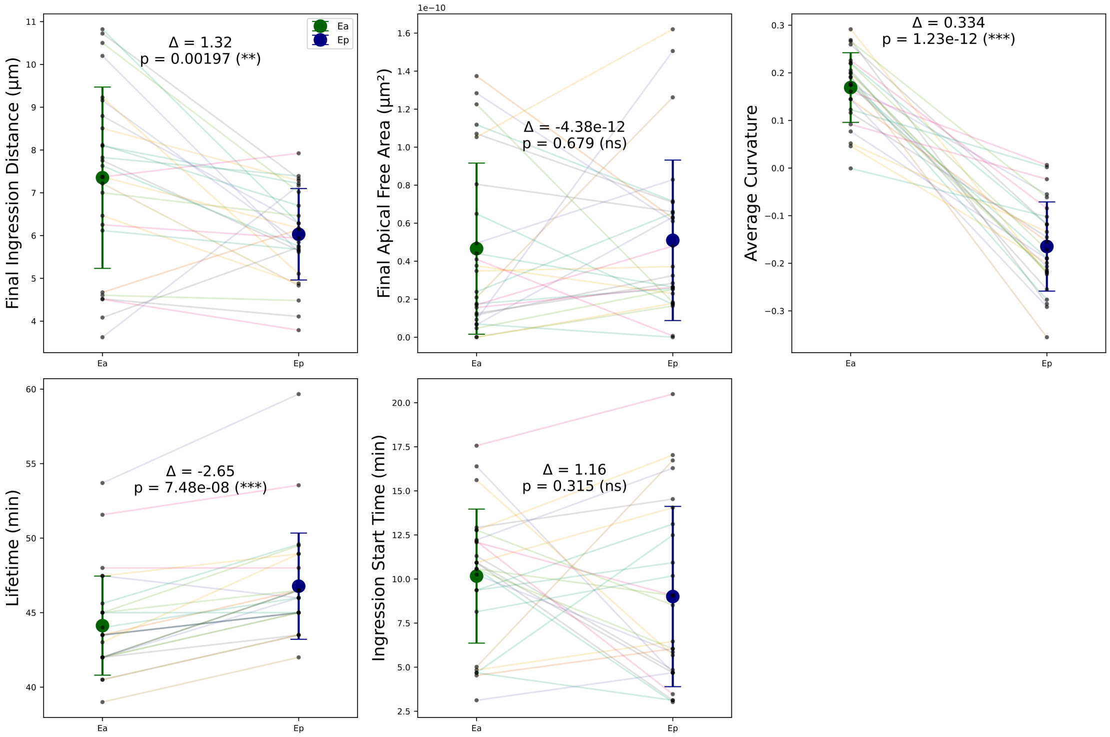


Figure A14: Comparison of Ea and Ep cell metrics. (from left to right) Final ingression distance, Final apical free area, and Average E cell curvature ($H/H_0, H_0 = 10^5$), Lifetime, and ingression start time. The ingression start time is defined as the beginning of the first 6-minute interval where the ingression distance is continuously positive and increases by at least 0.5 µm. Error bars indicate ±SD and individual replicate values are overlaid (n=26). Each panel is annotated with the paired t-test result, showing the mean difference (Δ) and the corresponding $p$-value along with significance levels [$(***)$ for $p < 0.001$, $(**)$ for $p < 0.01$, $(*)$ for $p < 0.05$, and (ns) for non-significance].

#### A.8 Triple junction angles

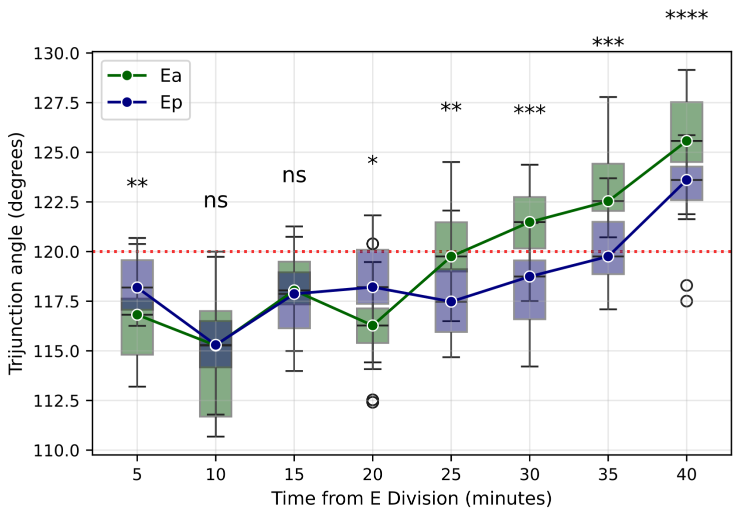


Figure A15: The evolution of internal trijunction angles was measured for Ea (green) and Ep (blue). In a system with uniform tension, these angles would be 120° (red dotted line). If cells build up tension their corresponding trijunction angle widen to preserve force balance. The analysis shows that both Ea and Ep experience a notable drop in tension around 10 minutes post-division. Following this, both cells undergo a gradual build-up of tension, evidenced by the increasing trijunction angles. This tension increase starts earlier (at t=20 min) for Ea than for Ep (t=30 min), which aligns with observations of their ingression speed (fig. A13). Significance markers represent the result of a paired t-test comparing the two cells at each time point: ns = not significant; * = $p < 0.05$; ** = $p < 0.01$; *** = $p < 0.001$.

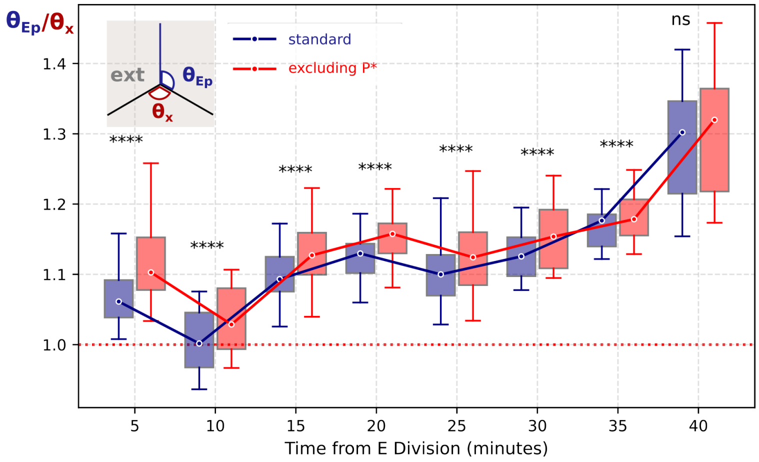


Figure A16: Apical trijunction angle ratio between Ep and its neighbors ($\theta_{Ep}/\theta_x$). This figures accompanies the plot in fig. 2C. This plot compares the standard condition where all neighboring cells are included (blue) with a condition where neighbors from the P-lineage (P3, P4) are excluded from the analysis (red). Significance markers denote the p-value from a paired t-test,: ns = not significant; **** = $p < 0.0001$. Pairing is done per combination of replicate and time-bin. Although the difference between the two conditions is highly statistically significant at most time points, the magnitude of this difference is very small.

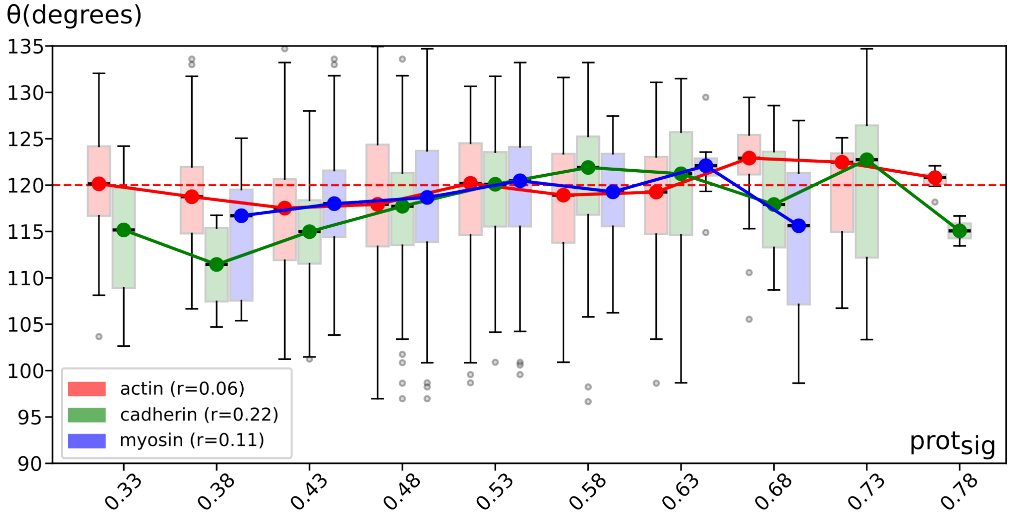


Figure A17: Relationship between cortical protein content at a cell-cell interface and the angle opposing this interface at the triple junction ($\theta$). The protein content for actin (red), E-cadherin (green), and myosin (blue) is binned on the x-axis, with the corresponding distribution of opposing triple junction angles shown as boxplots. Lines connect the mean angle for each protein across the bins. In principle, higher interfacial tension is balanced by a sharper opposing angle ($< 120°$). The dashed reference line at 120 degrees marks isotropic interfacial tensions at the triple junction. The data shows a significant mild positive correlation between the E-cadherin signal and the triple junction angle (r=0.22, p=$2.2 \times 10^{-9}$), consistent with the principle that higher cadherin-based adhesion lowers interfacial tension, resulting in a wider opposing angle. A very weak, but still significant positive correlation is also observed for myosin (r=0.11, p=0.0026), while actin shows no significant linear correlation (r=0.06, p=0.11).

#### A.9 Protein analysis

##### A.9.1 Protein quantification

First, the raw protein signal from a microscopy time-lapse is smoothed using a Gaussian kernel with a sigma of 0.2 microns. The smoothed signal is then projected onto the cell meshes, where each mesh node takes on the value of the nearest pixel. To account for bleaching over time and signal weakening with depth, a linear generalized additive model (GAM) is applied. Time (t) and the logarithm of the depth (log(z)) are fitted to piecewise second-order splines using the following pyGAM syntax:

```
gam = LinearGAM(s(t, n_splines=10, spline_order=2) +
                s(log(z), n_splines=20, spline_order=2))
```

The signal is subsequently corrected by taking the residual, which is the difference between the measured signal and the GAM prediction (representing the depth/time trend). These residuals are then normalized to produce the final score: values are first clipped to fall within ±3 SD of the residual mean, and then these clipped values are linearly scaled to the range [0, 1] using the clipping boundaries as the minimum and maximum for scaling. For calculating the protein value at a cell-cell interface or for an entire cell, a simple average of the corresponding mesh nodes is used.

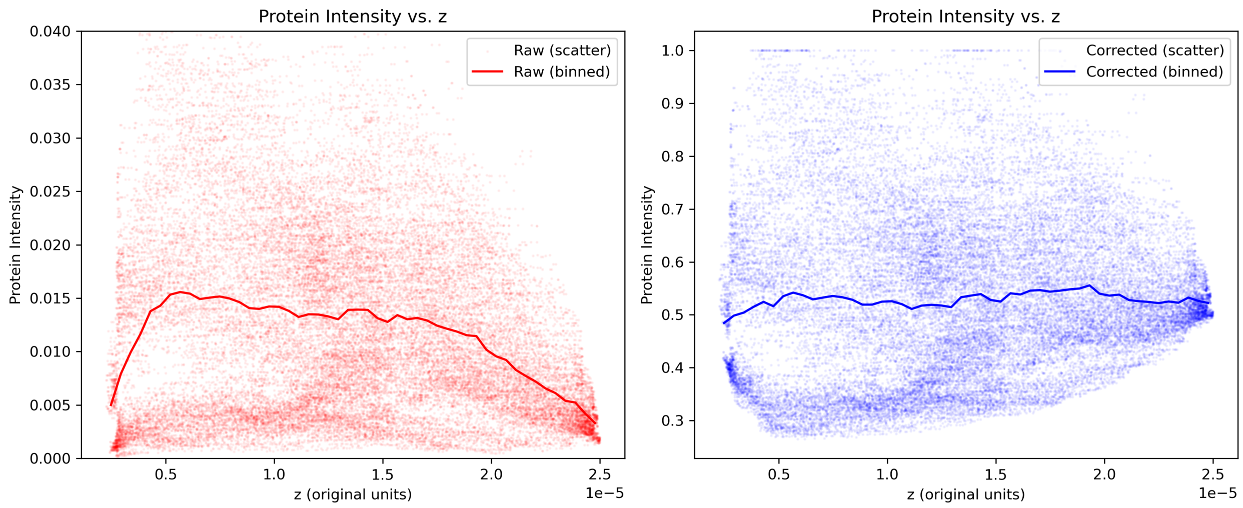


Figure A18: Example of actin signal correction (replicate wt08)

##### A.9.2 Statistical analysis of protein metrics

To quantify the factors influencing the protein distribution metrics defined in appendix B.5, a statistical analysis was performed for each protein (E-cadherin, myosin, and actin) and for each metric ($prot_{sig}$, $prot_{com}$, and $prot_{moi}$). The analysis focuses on three key metrics derived from protein intensity measurements on the cell surface. The base metric, $prot_{sig}$, represents the normalized protein signal intensity. From this, two distributional metrics are calculated: $prot_{com}$, which measures the asymmetry of the protein distribution by quantifying the distance between a surface's geometric centroid and its intensity-weighted center of mass, and $prot_{moi}$, which describes whether the protein signal is concentrated towards the center ($< 1$) or dispersed towards the periphery ($> 1$) relative to a uniform distribution. The statistical analysis was conducted at two distinct levels: for whole cells ('cell' analysis) and for the interfaces between cells ('surf' analysis).

First, the data is aggregated to create a single data point per replicate for each feature. For the 'surf' analysis, measurements for opposing interfaces (e.g., A-B and B-A) are averaged per replicate. For the 'cell' analysis, all measurements for a given cell are averaged per replicate. The metric values themselves were not transformed or normalized prior to fitting.

An Ordinary Least Squares (OLS) linear model was then fitted to the aggregated data. For the 'surf' analysis, the model predicts the metric value ($\hat{y}$) for an interface between cell $j$ and cell $k$ using the following general form:

$$\hat{y}_{jk} = \beta_0 + \beta_j + \beta_k + \delta_{jk} \qquad \text{(for a cell-cell interface)} \qquad (1)$$

$$\hat{y}_{jj} = \beta_0 + \beta_j + \gamma_{\text{free}} \qquad \text{(for an apical surface)} \qquad (2)$$

Here, $\beta_0$ is the global intercept, $\beta_j$ and $\beta_k$ are the coefficients for the additive contributions of cells $j$ and $k$, $\gamma_{\text{free}}$ is a specific coefficient for apical surfaces, and $\delta_{jk}$ represents the non-additive synergy term unique to the interface between cells $j$ and $k$. The model for the 'cell' analysis is a simplified version, predicting the metric for cell $j$ as $\hat{y}_j = \beta_0 + \beta_j$.

For the analysis of the $prot_{sig}$ metric at cell interfaces, a LASSO (L1) regularization was applied exclusively to the synergy coefficients ($\delta_{jk}$). This approach selectively shrinks the coefficients of non-essential synergy terms towards zero, functioning as a feature selection method to identify the most significant interface-specific effects. For the $prot_{com}$ and $prot_{moi}$ metrics, as well as for all cell-level analyses, a standard OLS model was used without regularization.

##### A.9.3 Cell level: mean concentrations

fig. A19

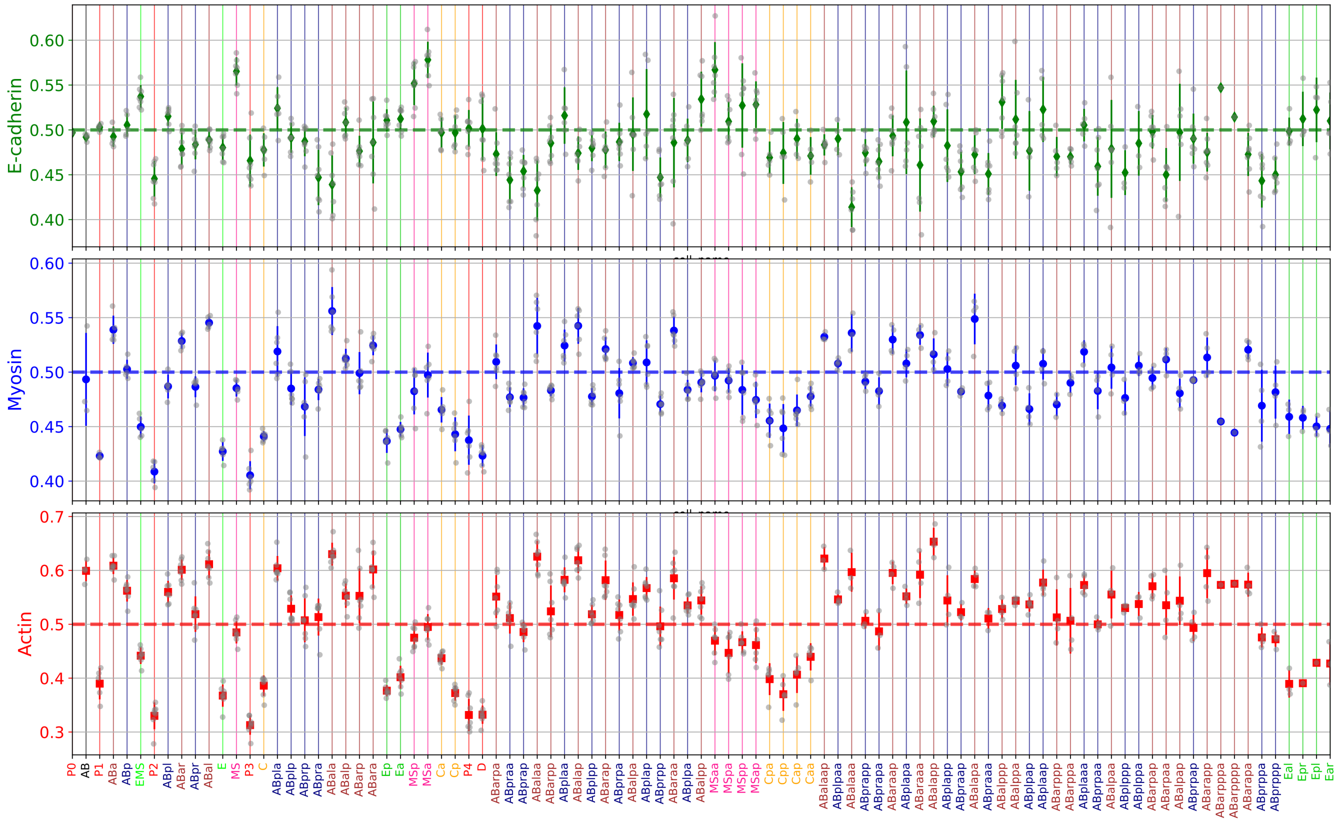


Figure A19: Mean cortical protein concentrations per cell. The cells along the x-axis are sorted by their birth time and include the cells from the zygote up to the Ea and Ep daughters. Each gray dot represent the time-averaged signal for a single replicate. The main colored markers indicate the mean signal averaged across all replicates, with the corresponding error bars showing the standard deviation across those replicates. The horizontal dashed line in each panel is set at a reference signal level of 0.5.

##### A.9.4 Cell level: daughter cell ratios

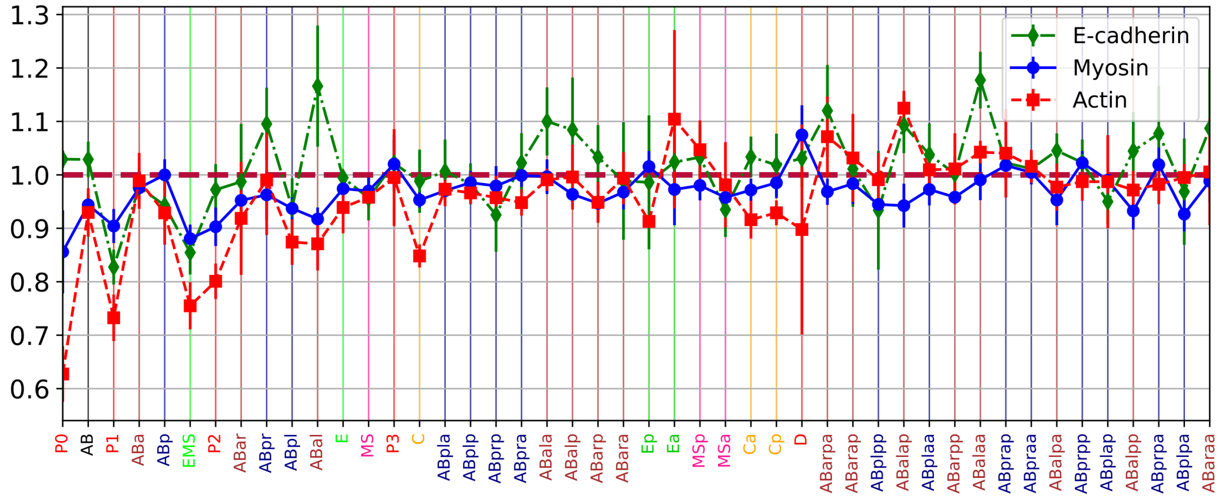


Figure A20: Cortical protein ratios between daughter cells. The x-axis shows the mother cells, sorted by their time of birth. Each data point represents the mean ratio for a given mother cell, with error bars indicating the standard deviation across replicates. The ratio is calculated by dividing the cortical protein signal of the anterior (or left) daughter cell by that of its posterior (or right) sister cell. For example, for the mother cell AB, the ratio is the signal in ABa divided by the signal in ABp.

##### A.9.5 Cell level: distributions over time per lineage

fig. A21

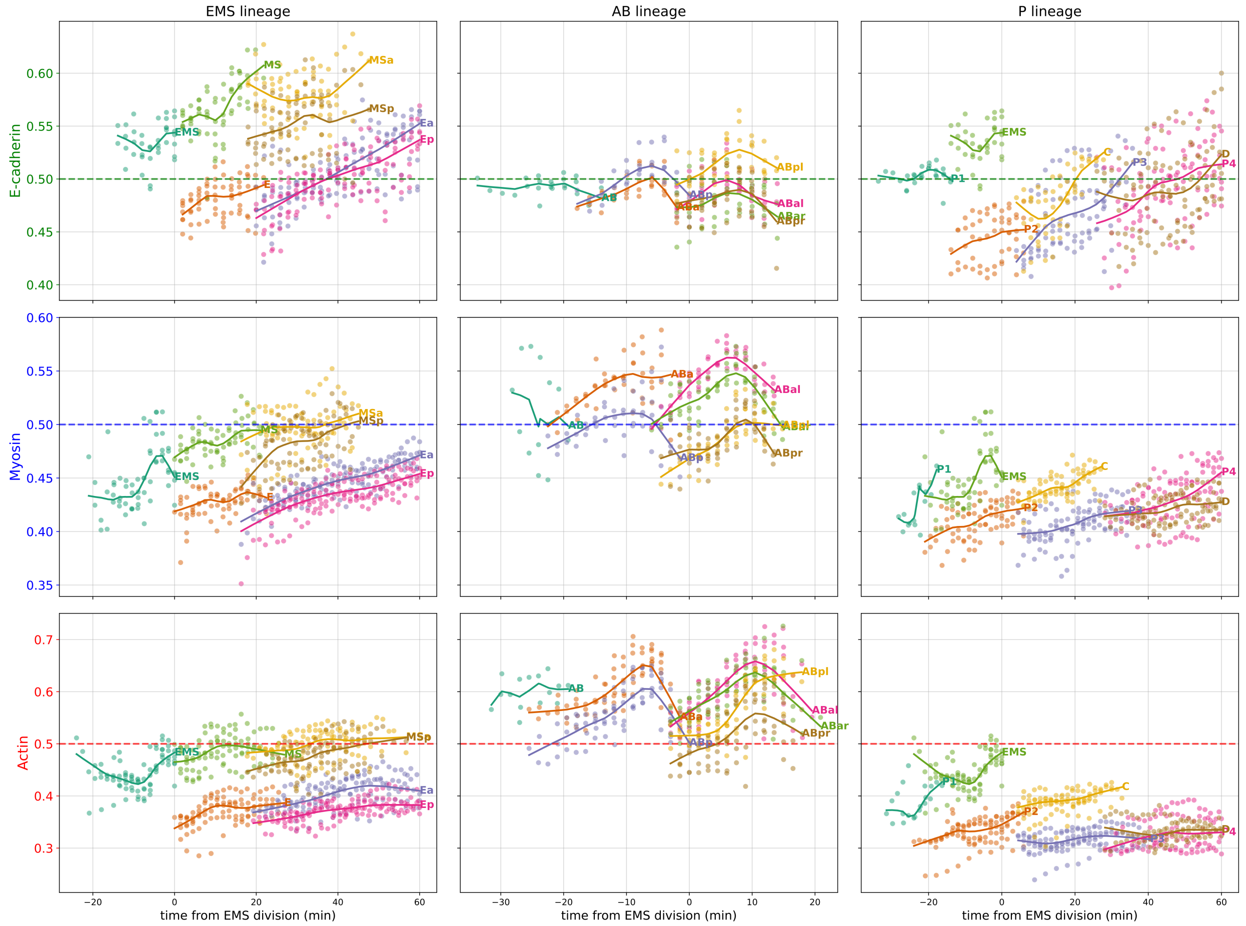


Figure A21: Cortical proteins (E-cadherin, myosin, and actin) within three lineages over time. Each dot represents a single measurement of protein signal at a specific time point from a given cell. For each cell, data from all replicates are aggregated, and a smoothed trendline is generated using a locally weighted scatterplot smoothing (LOWESS) filter. The x-axis indicates time relative to the division of the EMS cell, and the horizontal dashed line indicates a reference signal level of 0.5.

##### A.9.6 Cell interface level: mean concentrations and polarity

fig. A22

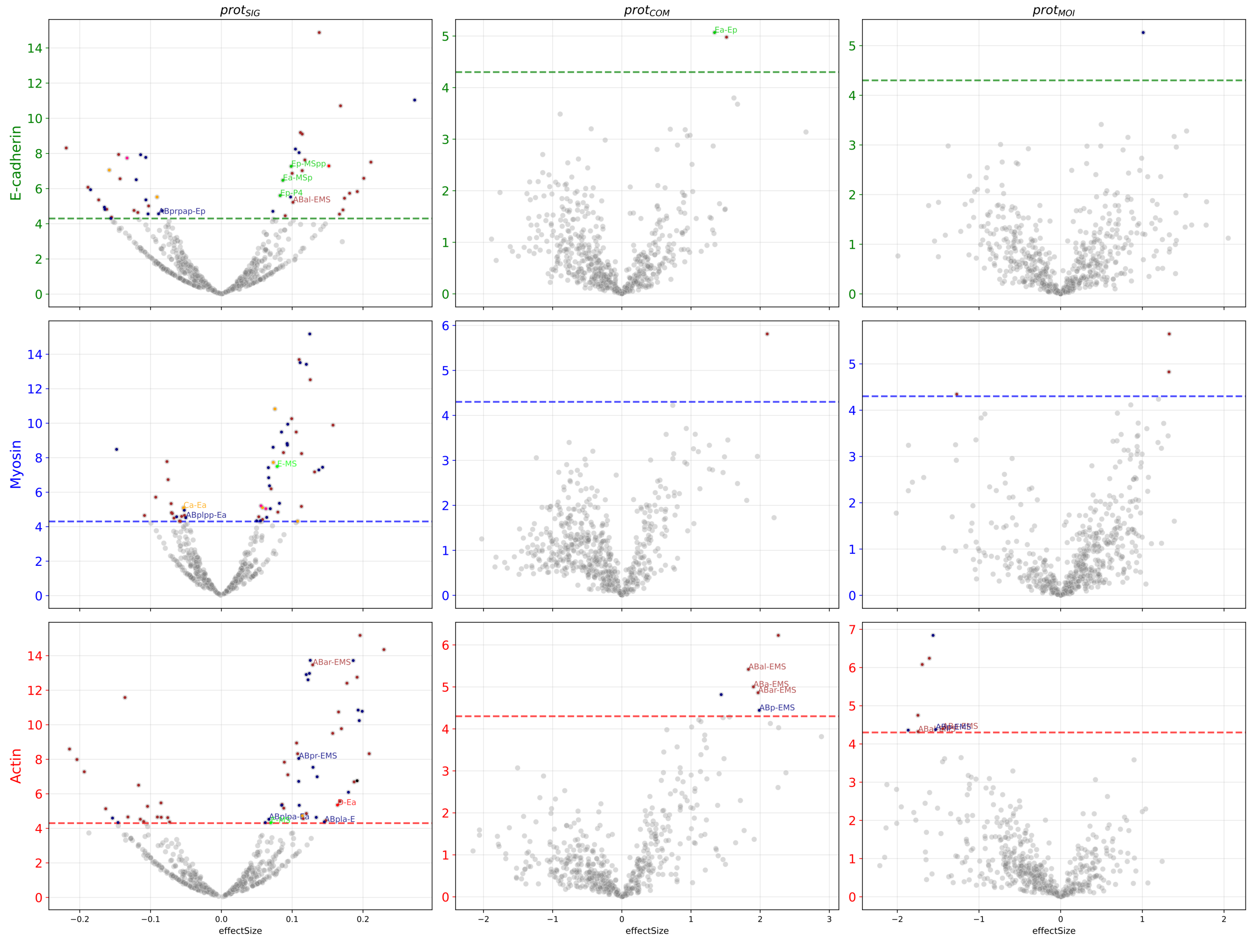


Figure A22: Volcano plots for protein metrics (appendix B.5) at individual cell-cell interfaces for actin, myosin, and E-cadherin. Metrics are: the mean signal intensity ($prot_{sig}$), the center of mass ($prot_{com}$), and the moment of inertia ($prot_{moi}$). $prot_{com}$ quantifies spatial asymmetry of the protein distribution, while $prot_{moi}$ indicates whether protein is concentrated towards the center or spread to the periphery of the interface. Interfaces with a p-value $< 0.0005$ are highlighted in color.

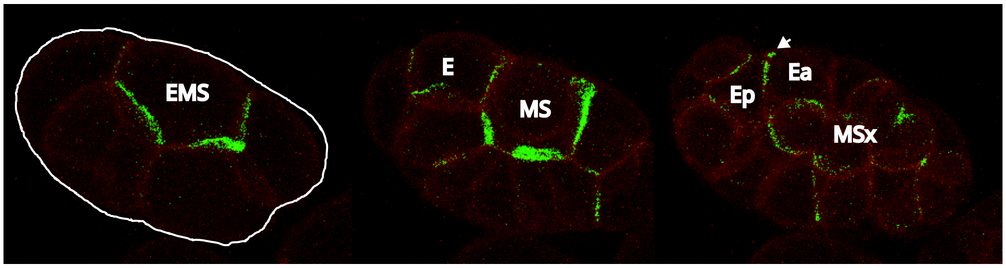


Figure A23: Distribution of E-cadherin over time in EMS lineage cells, emphasizing the highly asymmetric cadherin enrichment initially in MS versus E, with progressive cadherin accumulation observed in Ea/p. The white arrow marks the Ea-Ep anchor.

##### A.9.7 Cell interface level: a look at the E-cells and its neighbors

fig. A24

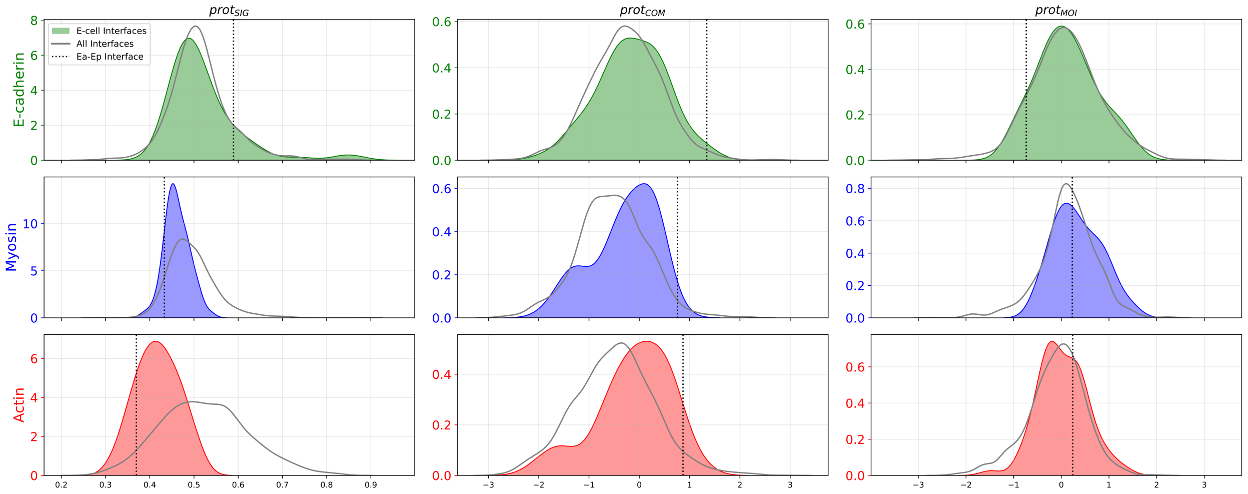


Figure A24: Distribution plots for protein metrics (appendix B.5) . Colored regions indicate all interfaces, gray regions highlight interfaces involving the E cells (Ea or Ep), and dotted lines mark the Ea-Ep interface. Interfaces of the E cells with their neighbors have less myosin and actin as compared to the average ($\texttt{prot}_{\texttt{sig}}$), which is to be expected as E is a descendant of P1. Within the E interfaces, Ea-Ep shows even less myosin and actin, and more more asymmetrically distributed. E-cadherin is no different in the E-interfaces as compared to the average, but here Ea-Ep is a clear outlier, showing more E-cadherin signal, with a strong asymmetric distribution ($\texttt{prot}_{\texttt{com}}$). The distribution around the center ($\texttt{prot}_{\texttt{moi}}$), shows no clear differences for any of the proteins.

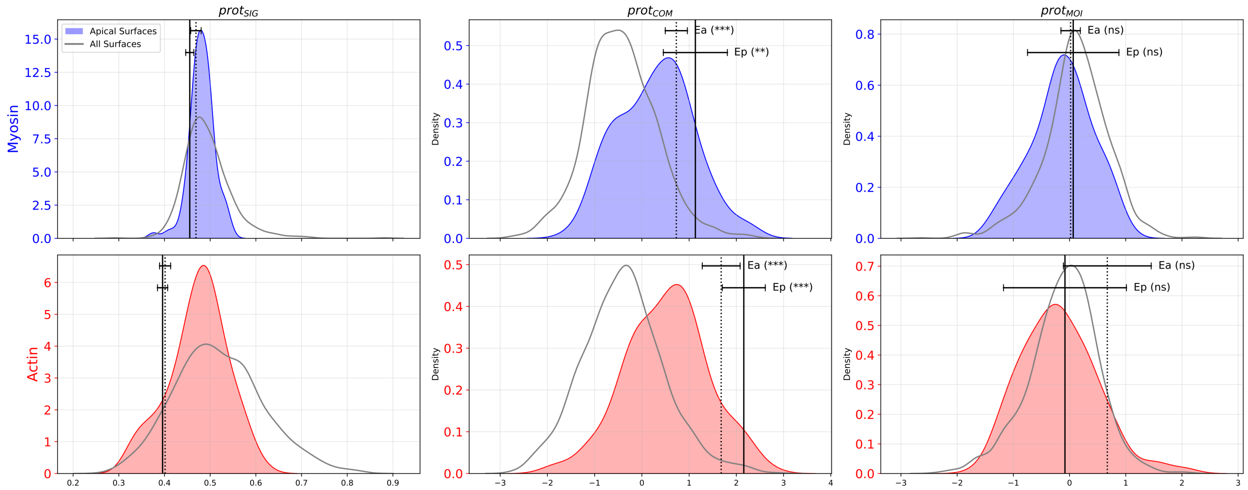


Figure A25: Kernel density estimates of protein metrics (Appendix appendix B.5) for myosin and actin on the apical surface. The grey outline represents the distribution across all cell surfaces, while the colored region shows the distribution for apical surfaces only. Vertical lines indicate the mean values for the apical surfaces of the Ea (dotted) and Ep (solid) cells, with horizontal bars showing the 95% confidence interval. Significance annotations report the results of one-sample t-tests, where, for $prot_{com}$ and $prot_{moi}$, it tests if their means are significantly different from zero. Stars denote significance levels: ns (not significant), * ($p < 0.05$), ** ($p < 0.01$), *** ($p < 0.001$), and **** ($p < 0.0001$). The observed distributions are inconsistent with a sarcomere-like network, which would feature a centrally enriched myosin population and a resulting $prot_{moi} < 0$ (Zhang et al., 2023), fig. B7. Here, for Ea and Ep, myosin appears relatively uniform, while actin exhibits both a significant spatial asymmetry (high $prot_{com}$) and peripheral enrichment (high $prot_{moi}$). This asymmetric peripheral actin enrichment is inconsistent with a simple purse-string model (which would have a lower $prot_{com}$) and instead supports the diffuse medioapical network architecture for *C. elegans* proposed by Zhang et al. (2023).

##### A.9.8 Statistical analysis of protein metrics at E-cell interfaces

To assess whether protein distribution metrics were significantly elevated at cell-$\mathrm{E}_{\mathrm{a/p}}$ interfaces, we performed a statistical analysis for each protein (E-cadherin, myosin, and actin) and for each metric ($prot_{sig}$, $prot_{com}$, and $prot_{moi}$) (fig. A26).

For each cell at a given time point, we calculated a delta metric ($\Delta$). This was defined as the difference between the metric's value on a specific cell-$\mathrm{E}_{\mathrm{a/p}}$ interface and the mean value of the metric across all other non-apical and non-basal interfaces of that same cell. This approach normalizes the metric against the cell's own baseline protein distribution.

To account for repeated measurements over time for each cell-cell interface, we fitted a Linear Mixed-Effects Model (LMM):

$$\Delta \sim 1 + t_{\text{centered}} + (1 + t_{\text{centered}} | \text{interface}) \tag{3}$$

Here, the response variable is the calculated $\Delta$. The time variable ($t_E$) was centered for each interface by subtracting the mean observation time for that interface ($t_{\text{centered}}$). The model includes $t_{\text{centered}}$ as a fixed effect to account for a common temporal trend, and ‘interface‘ and slope as a random effect, allowing each specific cell-$\mathrm{E}_{\mathrm{a/p}}$ interface to have its own baseline and a rate of change over time . This intercept represents the average $\Delta$ for that interface across its lifespan.

From the fitted model, we extracted the estimated random intercept for each interface. A one-tailed p-value was derived from the Z-score of this intercept to test the hypothesis that the average $\Delta$ is significantly greater than zero.

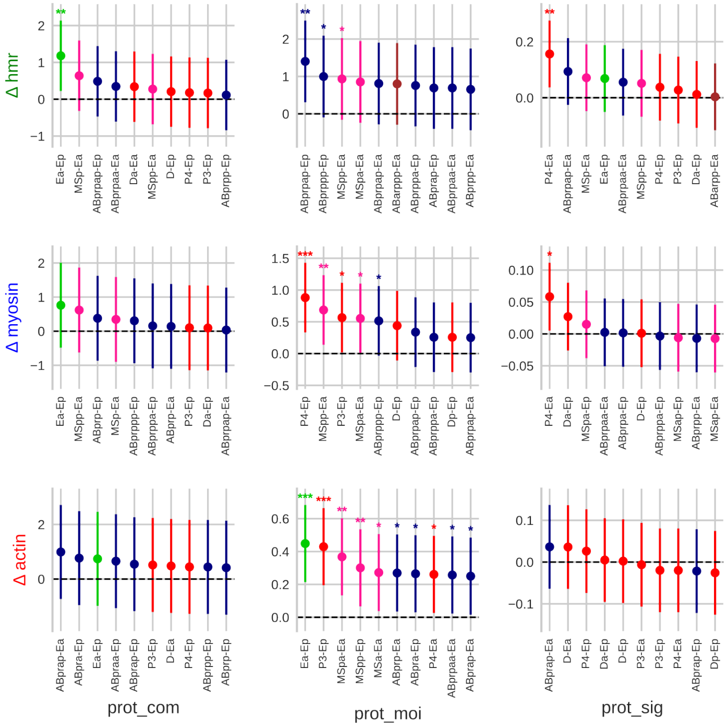


Figure A26: Statistical analysis of protein distribution metrics at interfaces with Ea and Ep (appendix A.9.8). Each panel displays the results for a specific protein (rows) and metric (columns). For each combination, the top ten most significant interfaces (lowest p-value) are shown. Each point represents the time-averaged delta (Δ) for a specific cell-cell interface, calculated as the deviation from the cell's own mean. The color indicates the name of the interfacing cell. Vertical lines show the 95% confidence interval for the mean delta. Significance levels for the one-tailed test ($\Delta > 0$) are indicated by asterisks: ($***$) for $p < 0.001$, ($**$) for $p < 0.01$, and ($*$) for $p < 0.05$.

##### A.9.9 Cell Interface level: selected interfaces over time

**Kymograph** fig. A27 fig. A28 fig. A29 fig. A30

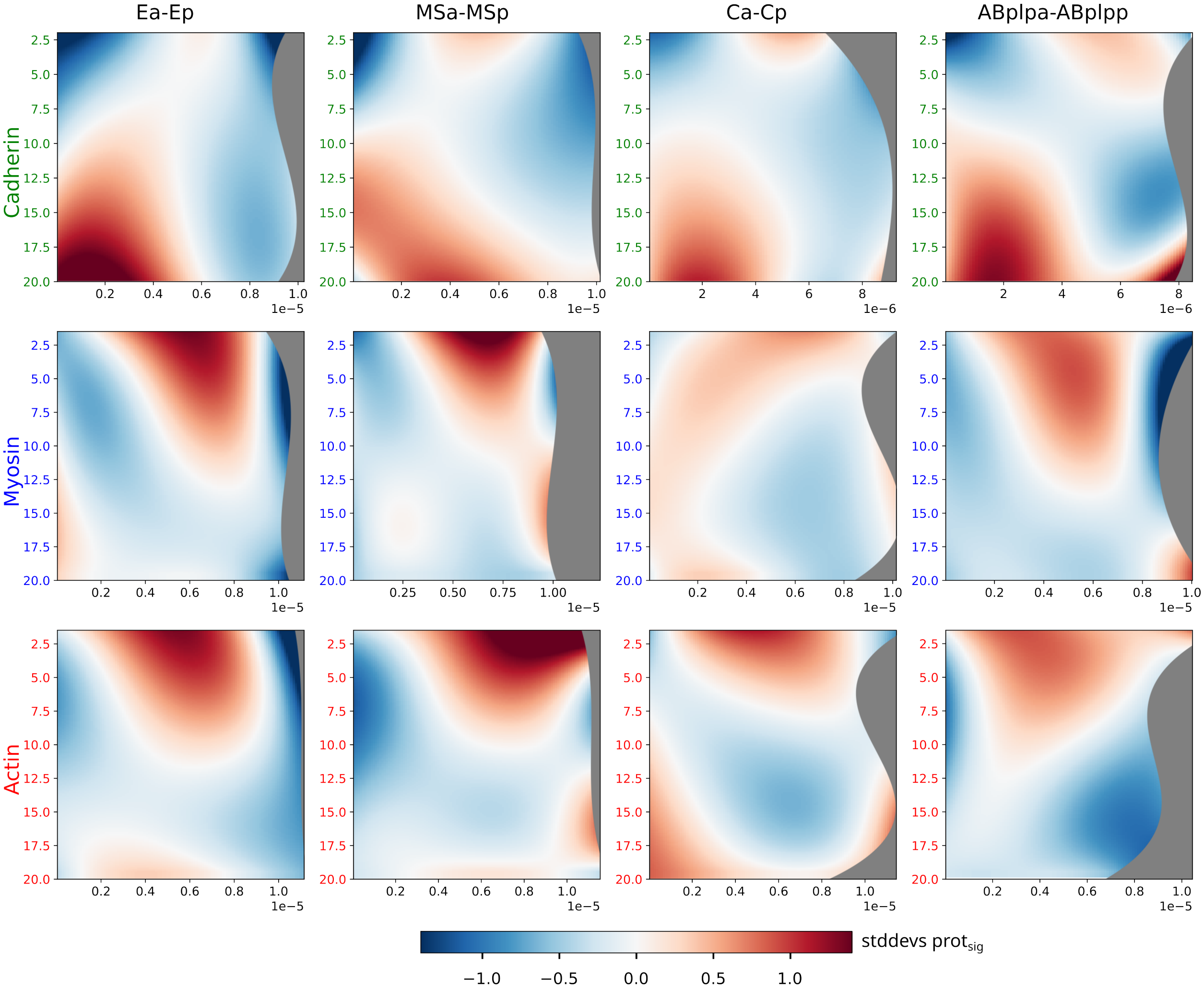


Figure A27: Kymographs (appendix B.5.4) showing the spatio-temporal dynamics of cortical proteins at the shared interface of four daughter cell pairs: Ea-Ep, MSa-MSp, Ca-Cp, and ABplpa-ABplpp. The vertical axis represents time after division up to 20 minutes, while the horizontal axis indicates the distance from the most apical point of the interface. Each panel's colormap is centered (white) on the mean protein concentration for that specific interface, with red and blue indicating enrichment and depletion relative to this mean. This per-interface normalization means that concentrations cannot be quantitatively compared between different columns. Greyed-out zones represent areas where no data was available. At the Ea-Ep interface, E-cadherin shows a distinct build-up at the apical side, a trend also observed, albeit less prominently, in the other cell pairs. In contrast, myosin and actin exhibit an initial enrichment across the interface, a remnant of the cytokinetic ring, which then dissipates over time in all pairs.

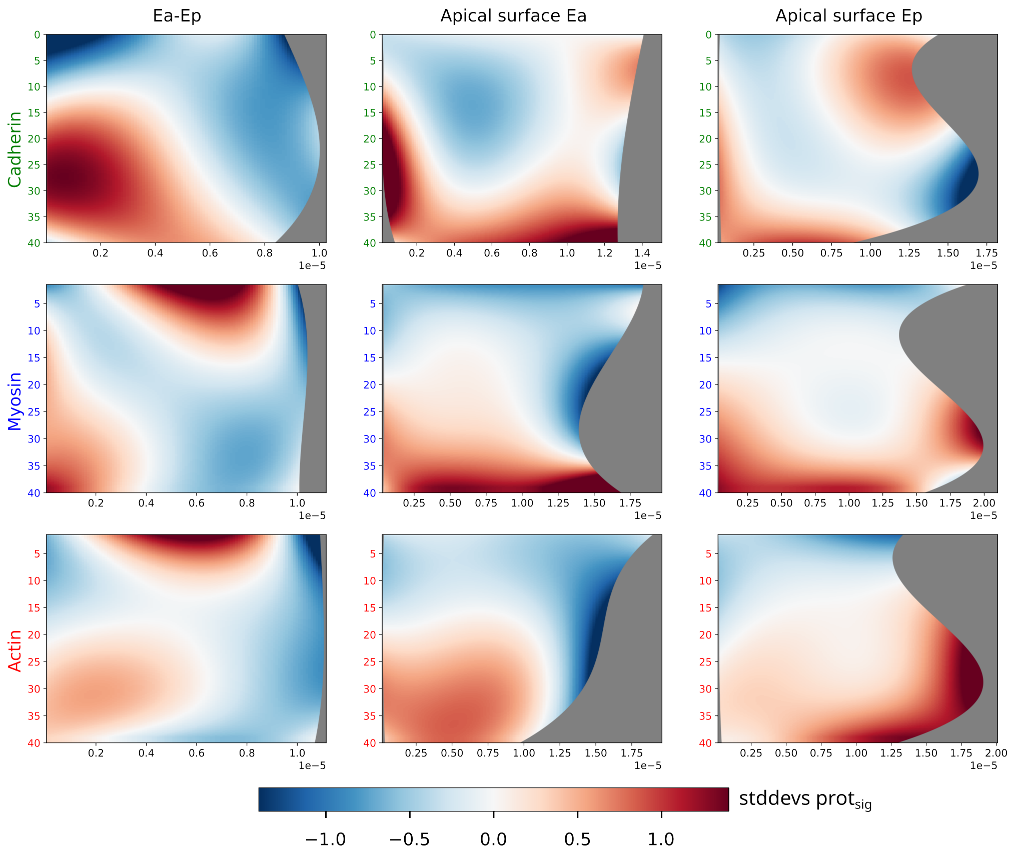


Figure A28: Kymographs (appendix B.5.4) of cortical protein dynamics, comparing the shared Ea-Ep interface (left column) with the adjacent apical surfaces of Ea (middle column) and Ep (right column). The vertical axis represents time after E-cell division up to 40 minutes. For the Ea-Ep interface, the horizontal axis is the distance from the most apical point; for the apical surfaces, it is the distance along the surface starting from the junction with the Ea-Ep interface. E-cadherin accumulates at the apical side of the Ea-Ep contact, stabilizing around 20 minutes, while remaining at low levels on the adjacent apical surfaces. Myosin enrichment begins around 10 minutes at the Ea-Ep interface and is also observed on both the Ea and Ep apical surfaces. Actin enrichment at the interface appears later, around 20 minutes; its distribution is uniform on the apical surface of Ea, but on Ep it is more pronounced at distances further away from the cell-cell contact.

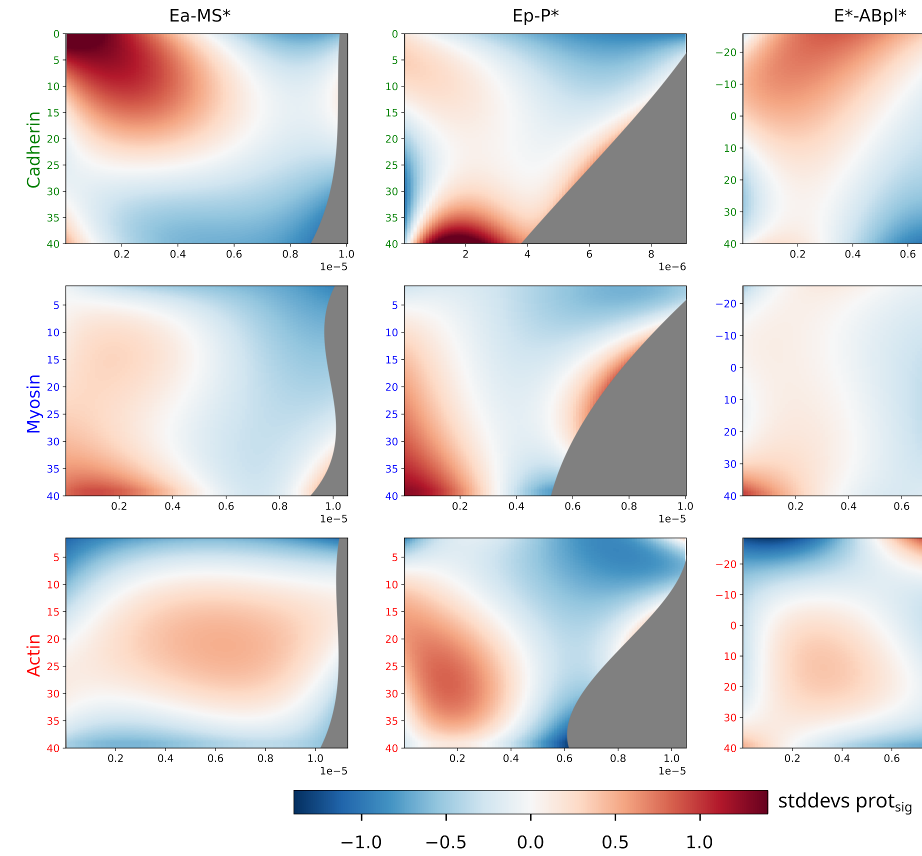


Figure A29: Kymographs (appendix B.5.4) of cortical protein dynamics at the interfaces between the endodermal precursors (Ea, Ep) and their neighbors: MS-lineage cells (left), P-lineage cells (middle), and ABpl-lineage cells (right). The horizontal axis represents the distance from the most apical point of the respective interface. The vertical axis is time relative to E-cell division, with the time window for each panel reflecting the existence of that specific cell-cell contact. No signs of significant E-cadherin build up on the apical side (cfr clutch) with the cells of the MS or ABpl lineage . Enrichment does occur at the Ep-P* interface, but this accumulation is broad rather than distinctly apical. A slight apical enrichment of myosin is visible at both the Ep-P* and E*-ABpl* interfaces. Actin does not show this apical polarization but is strongly enriched across the entire E*-ABpl* interface. In summary, there is no clear indication of an apical junction-like build-up of E-cadherin at the interfaces with the neighboring cells.

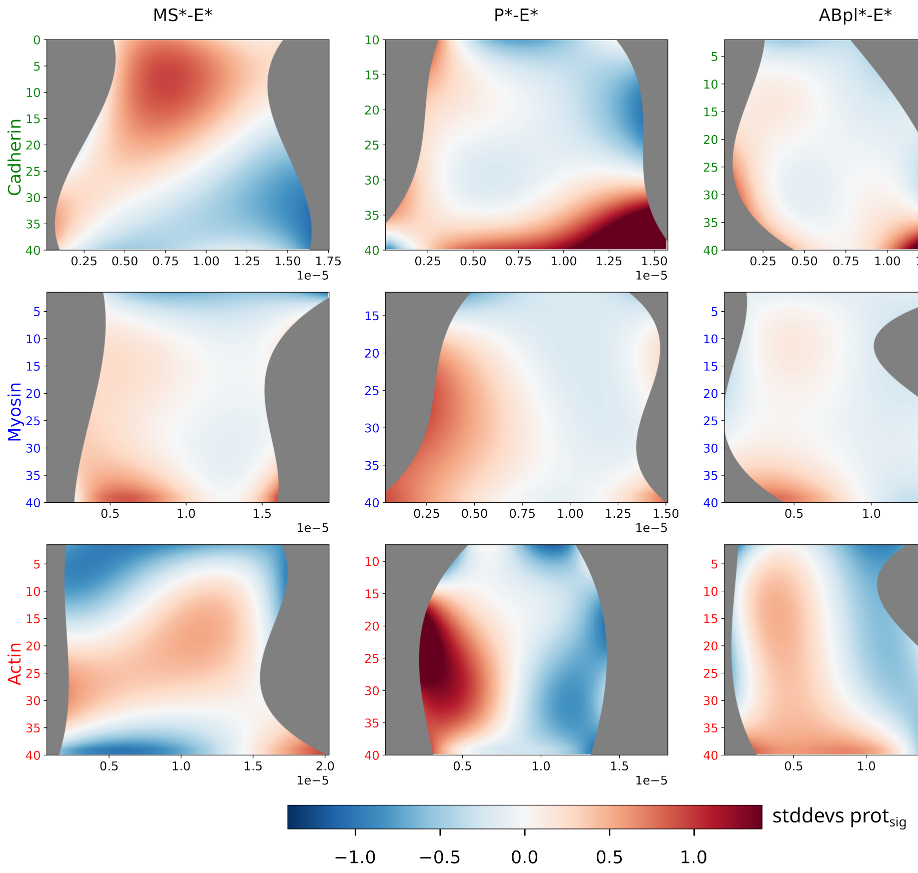


Figure A30: Kymographs (appendix B.5.4) of neighboring cells with the E cells . The horizontal axis is the distance along the neighbor's apical surface, starting from its contact with an E cell, used to assess for protein polarization towards the gastrulation cleft. For each protein (row), the colormap is normalized using data from the MS* apical surfaces (left column) and then applied across all three panels to allow for comparison. E-cadherin is strongly enriched on the apical surface of P* neighbors, building up over time but without clear polarization; this enrichment is weaker on MS* neighbors. In summary, these results do not show a consistent pattern of protein polarization towards the tips of the covering cells.

#### A.10 Simulations - models

See appendix C

#### A.11 Simulations - results

##### A.11.1 Simulation scenarios

##### A.11.2 Apical tension scenarios

fig. A31 fig. A32 fig. A33 fig. A34

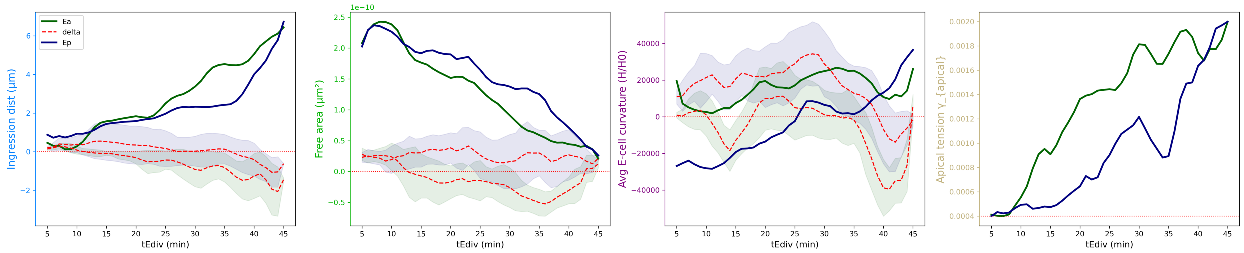


Figure A31: Ingression metrics for Ea (dark blue) and Ep (dark green) in the $A_{\text{dist}}D$ simulation, showing the mean of 17 replicates (shaded region: SD, 1-min time-bin). Dashed red lines indicate deviation from target data. The simulation captures Ep's ingression acceleration but develops a final lag of 1 $\mu$m. Apical area is reduced, with Ep retaining excess area of around 50 $\mu m^2$ while Ea is a closer fit. The simulated curvature of Ep is excessively convex compared to the target. In contrast, Ea's curvature, after an initial increase, incorrectly decreases during the final stage of ingression.Similarly, Ep's mean curvature is excessively convex, and Ea's incorrectly decreases in the final phase. The cells exhibit distinct apical tension profiles: Ea displays a sharp increase 10 min post-E-division, whereas Ep's tension rises gradually, accelerating only in the final 10 minutes.

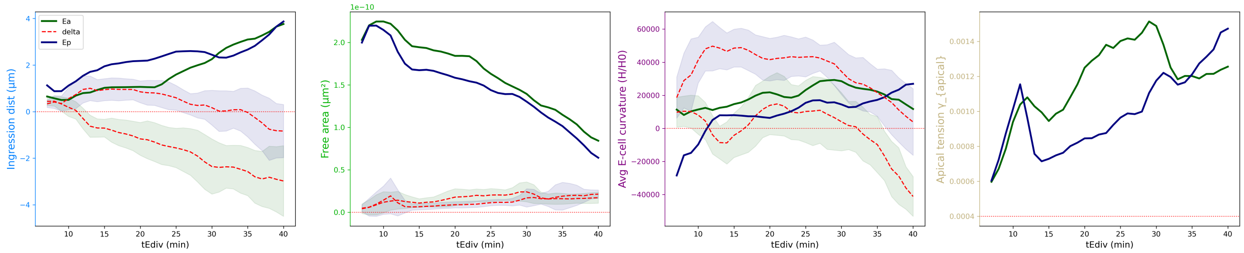


Figure A32: Ingression metrics for Ea (dark blue) and Ep (dark green) in the $A_{\text{area}}D$ simulation. Plots show the mean of 17 replicates (shaded region: SD, 1-min time-bin); dashed red lines indicate deviation from target data. The dynamics are highly similar to the $A_{\text{dist}}D$ scenario, with comparable final lags in ingression distance and analogous deviations in cell curvature and tension. The key improvement in this scenario is the accurate tracking of the apical free area. (cf. Table fig. A35).

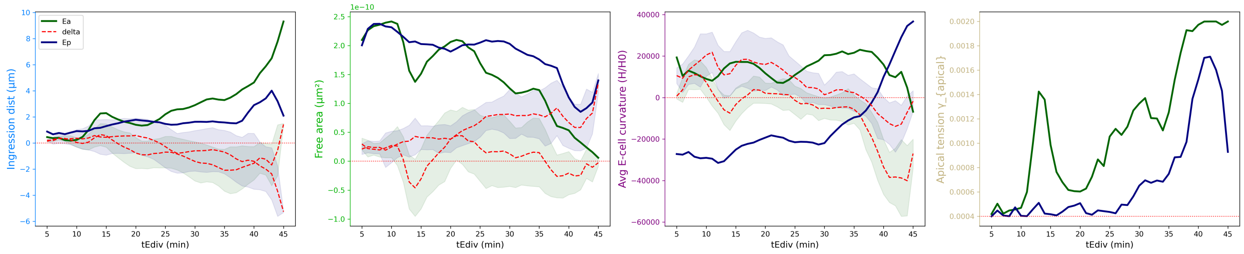


Figure A33: Ingression metrics for Ea (dark blue) and Ep (dark green) in the $A_{\text{curv}}D$ simulation. Plots show the mean of 17 replicates (shaded region: SD, 1-min time-bin); dashed red lines indicate deviation from target data. While this scenario more accurately tracks the target cell curvature, it fails to match ingression distance and apical area.

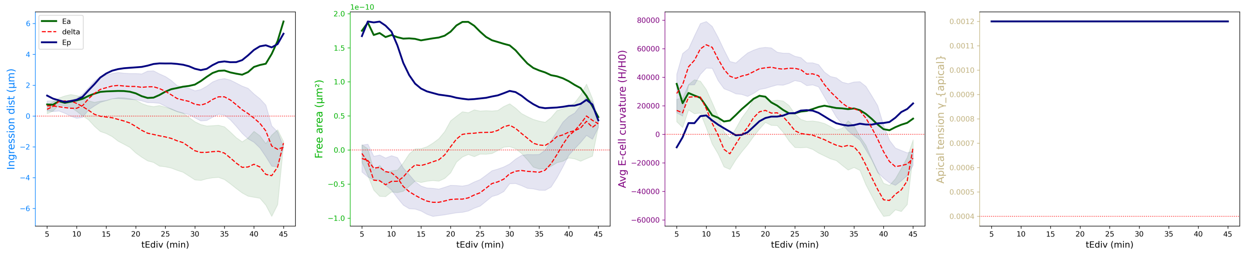


Figure A34: Ingression metrics for Ea (green) and Ep (blue) in the $A_{\text{abs}}D$ scenario, which applies a constant apical tension. Data represents the mean of 17 replicates (shaded region: SD, 1-min bin), with red dashed lines showing deviation from target.

##### A.11.3 Apical tension

fig. A35 fig. A36

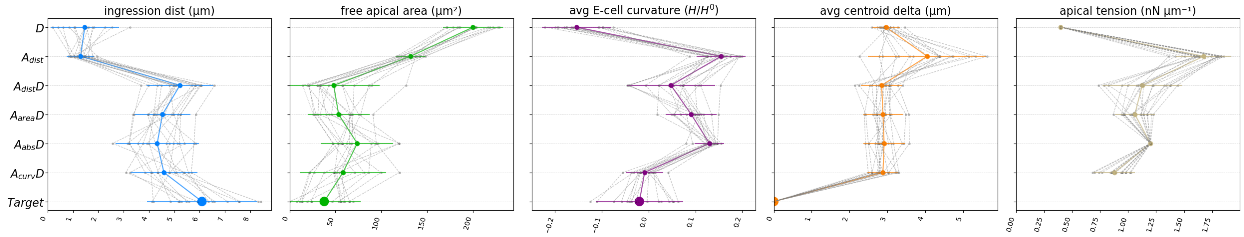


Figure A35: Evaluation of ingression metrics for simulation scenarios: **D**, which implements only cell divisions without any apical tension; $A_{\mathbf{dist}}$, where apical constriction tracks ingression distance, without cell divisions; $A_{\mathbf{dist}}D$, which tracks ingression distance; $A_{\mathbf{area}}D$, where tension tracks the free apical area; and $A_{\mathbf{curv}}D$, where apical constriction tracks E-cell curvature. The results show that increasing apical tension leads to greater ingression distance, higher E-cell curvature, and reduced free apical area, with a minimal effect on overall cell positioning. Notably, matching the target ingression distance and area, as in $A_{\text{area}}D$, results in excessive E-cell curvature.

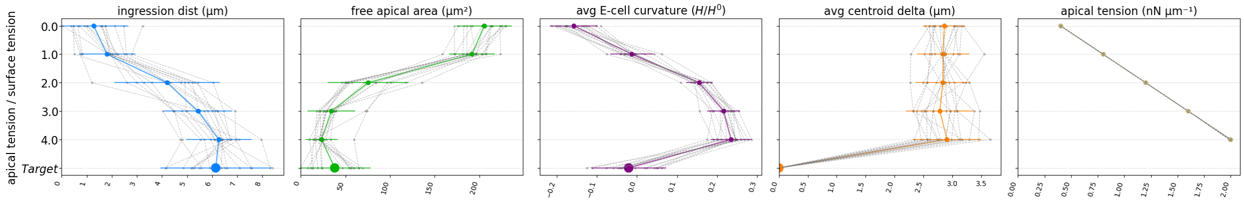


Figure A36: Effect of increasing apical tension on simulated ingression metrics ($A_{\text{abs}}D - sweep$). An added apical tension of three times the surface tension result in an ingression distance and free apical area close to the target, but leads to an excess curvature of the E-cells. This graph also serves to show that curvatures can be used as a proxy for apical tension, following the reasoning that increasing apical tension, increases cell pressure, which leads to more convex E-cell interfaces.

##### A.11.4 Cell division

**Cell division variations** fig. A37 fig. A39 fig. A41 fig. A40

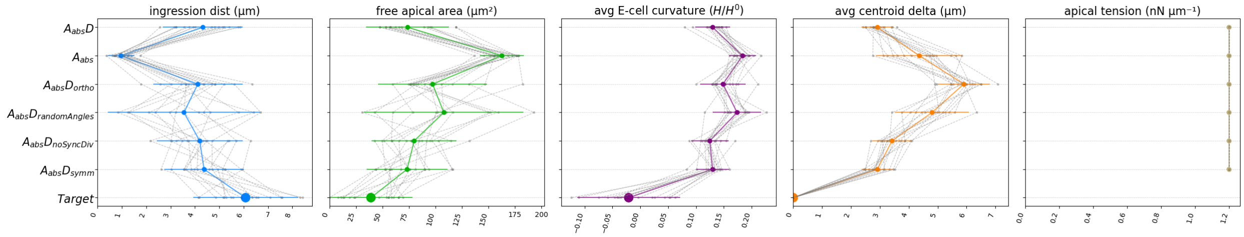


Figure A37: Effect of division variations on simulated ingression metrics. All scenarios were run with constant apical tension (3X surface tension). The plot compares the reference $A_{\mathbf{abs}}D$ scenario to variations including orthogonal ($A_{\mathbf{abs}}D_{\mathbf{ortho}}$) or randomized ($A_{\mathbf{abs}}D_{\mathbf{randomAngles}}$) division angles, unsynchronized divisions ($A_{\mathbf{abs}}D_{\mathbf{noSyncDiv}}$) ((fig. A41), and symmetrical divisions ($A_{\mathbf{abs}}D_{\mathbf{symm}}$). Altering division angles disrupts cell positioning (increased centroid delta) and hinders ingression. In contrast, modifying division timing or symmetry has a negligible effect on the final ingression metrics compared to the base scenario.

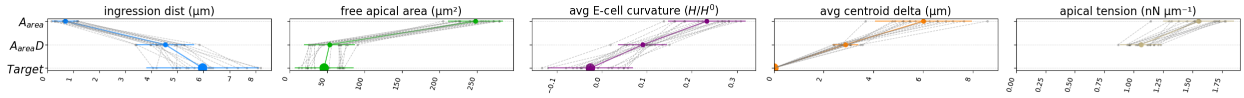


Figure A38: Effect of eliminating cell divisions on ingression metrics. Contrasting $A_{area}$ with $A_{area}D$ shows that eliminating cell divisions leads to failed ingression, even with higher apical tension.

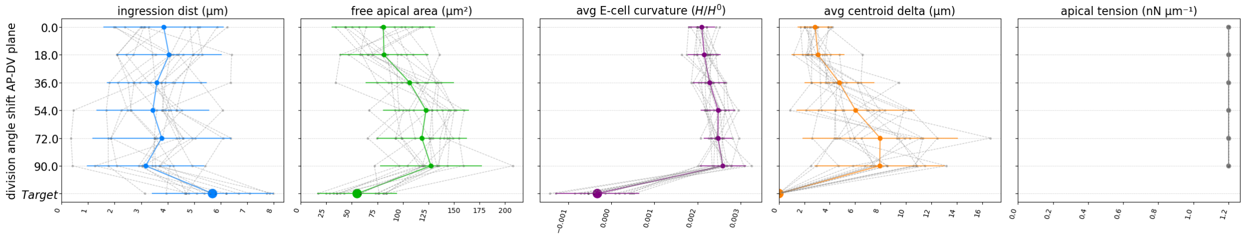


Figure A39: Effect of rotating division angles in the AP-DV plane on simulated ingression metrics ($A_{\mathrm{abs}}D - sweepDivAngle$). All simulations run with constant apical tension. Shifting the division angle away from the in vivo orientation (0 degrees) has a negligible impact on ingression distance but significantly hinders apical area reduction and leads to less precise cell positioning.

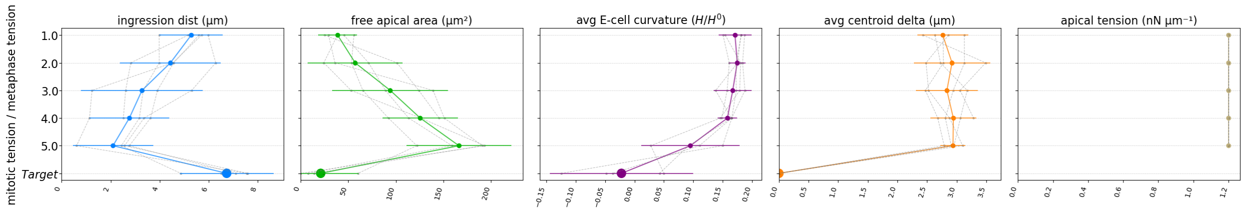


Figure A40: Effect of modulation of mitotic tension on simulated ingression metrics ($A_{\mathrm{abs}}D - sweepDivTension$). All simulations are run with constant apical tension. The results show that increasing mitotic tension has a strong negative impact on ingression. Higher levels of mitotic tension lead to a significant reduction in ingression distance, an increase in the final apical area. A mitotic tension of three times surface tension is the default setting. This suggests that pressure generated by mitotic rounding works against the forces driving apical constriction.

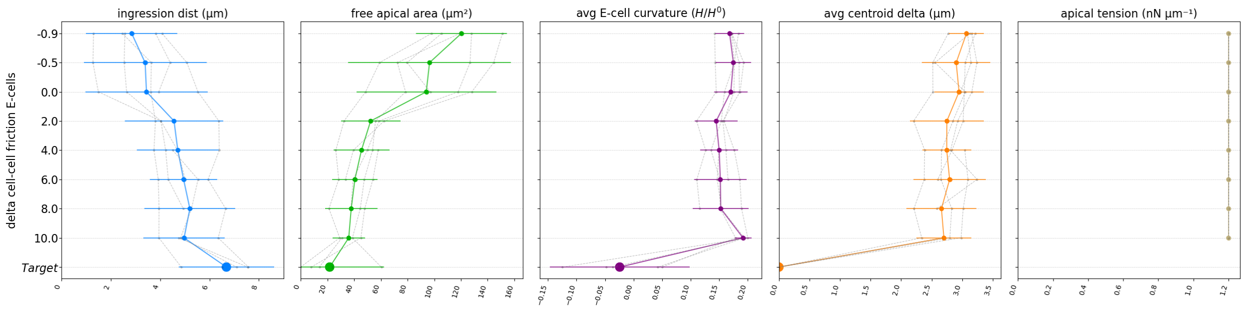


Figure A43: Effect of modulating cell-cell friction of Ea and Ep on simulated ingression metrics. The Y axis represents the added cell-cell friction expressed relative to the default cell-cell friction. Five replicates were used (crm04-08)

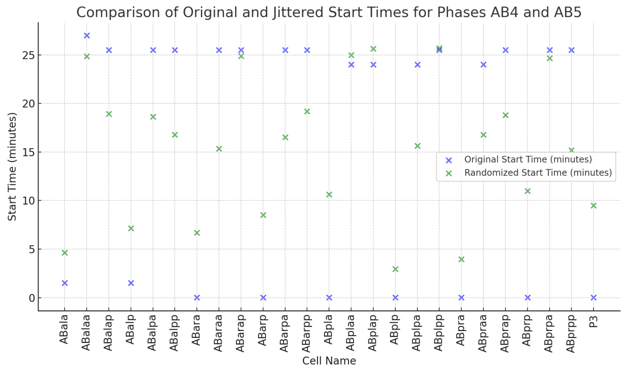


Figure A41: Plot detailing how $A_{abs}D_{noSyncDiv}$ breaks up synchronized AB division times by randomizing and spreading division out over the entire Ea/p gastrulation interval. Replicate crm04 is taken as an example.

##### A.11.5 Viscosities and frictions

fig. A42 fig. A44 fig. A46

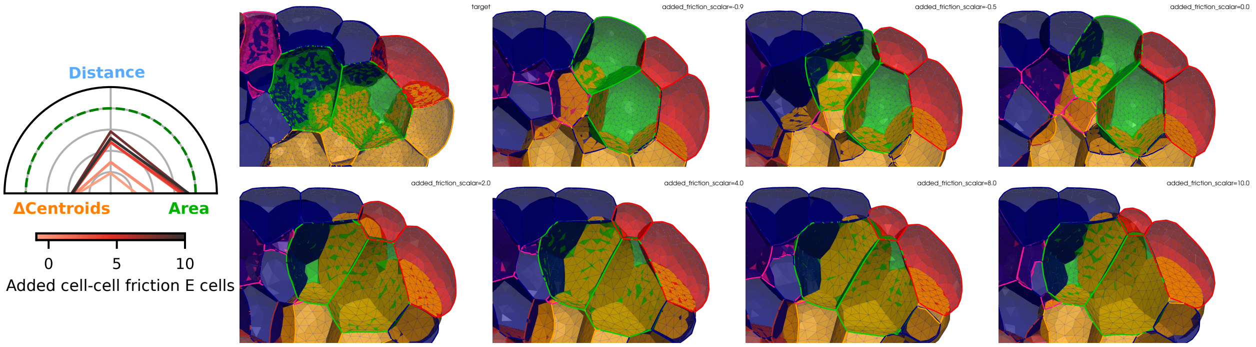


Figure A42: Effect of modulating cell-cell friction of Ea and Ep on simulated ingression metrics. A constant apical tension of 3X surface tension was applied. Five replicates were used (crm04-08). The added friction is expressed relative to the default cell-cell friction of 0.12 kPa*s/um, and following values were used: -0.9, -0.5, 0, 2, 4, 6, 8, 10. For example, an added friction equal to 2 represent a total friction of 0.36 kPa*s/um The banner uses replicate crm04 to illustrate.

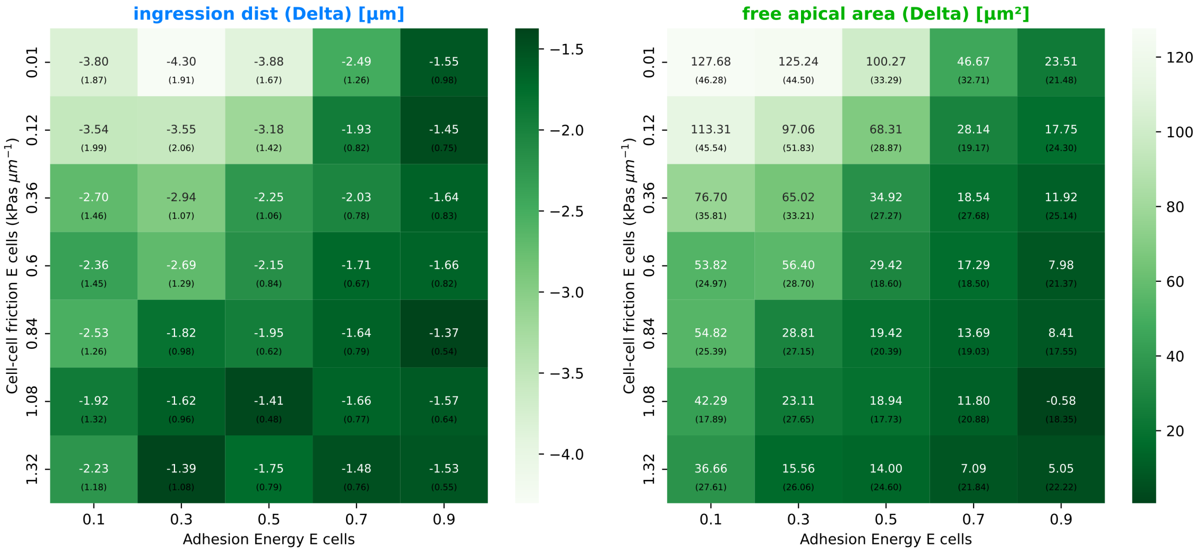


Figure A44: The effect of modulating adhesion density and cell-cell friction of the E-cells on simulated ingression metrics ($A_{abs}D - sweepDiff$). The x-axis represents the differential adhesion density for Ea and Ep and is expressed relative to surface tension ($\gamma_I$); the default value is 0.5, which corresponds to 0.2 nN $\mu$m$^{-1}$. The y-axis indicates the cell-cell friction, with a default value of 120 Pas $\mu$m$^{-1}$. A constant apical tension of 3X surface tension was applied. Values in each cell of the heatmap represent the mean across replicate simulations, with the standard deviation shown in parentheses. Five replicates (crm04-08) were used.

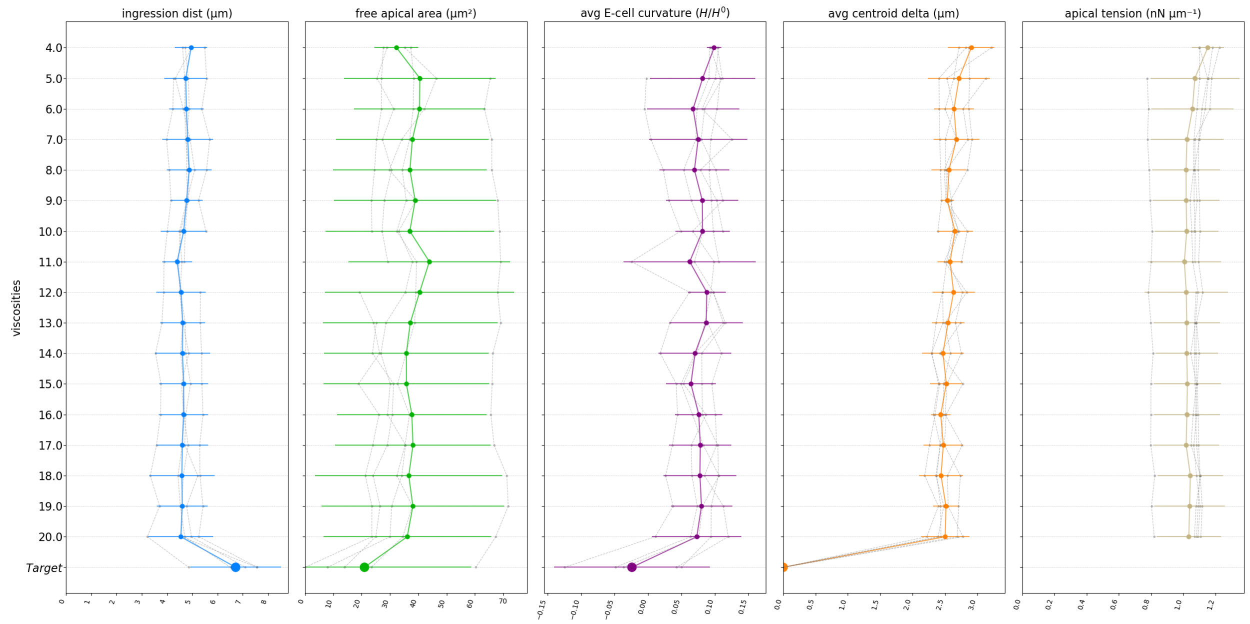


Figure A45: The effect of modulating frictions on simulated ingression metrics ($A_{area}D - sweepViscoBal$). Cortex viscosity, liquid viscosity and cell-cell friction are increased in tandem, always keeping roughly a 120:1:1 balance. The base setting used in other simulations is 11, which amounts to a medium viscosity of 0.09 kPa·s, a cortex viscosity of 11 kPa·s, and a cell-cell friction of 0.12 kPa·s/$\mu$m. Five replicates (crm04-08) were used. The results show that when the balance of friction parameters is preserved, the absolute viscosity values have a minimal impact on the simulation outcome, suggesting the system is in a regime that allows for sufficient mechanical relaxation.

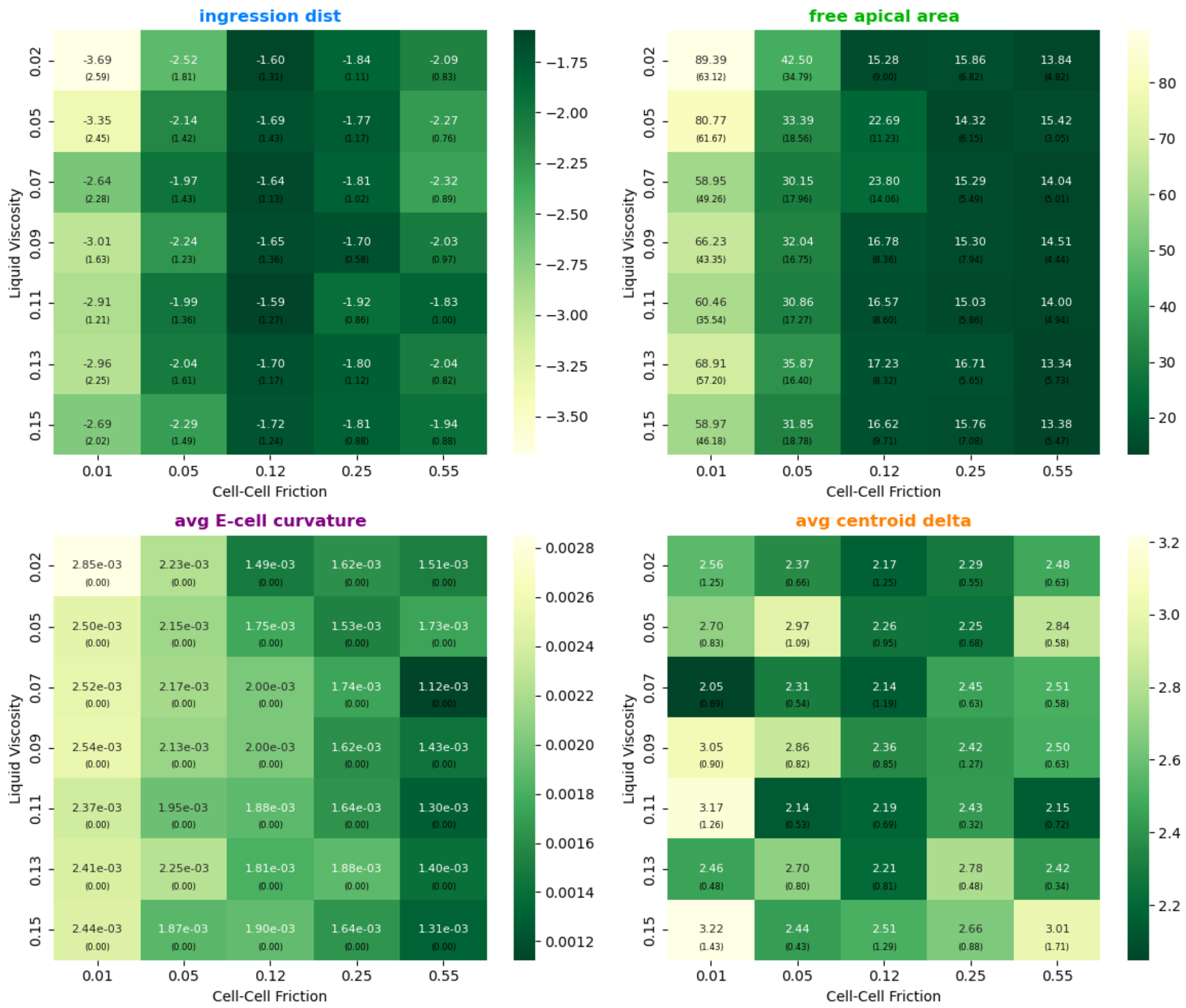


Figure A46: The effect of cell-cell friction and medium viscosity on ingression metrics ($A_{area}D - sweepVisco$) using $A_{area}D$ as the base case. Delta values are shown, and darker green represent metrics closer to target. The base friction settings are in the center of each heatmap (medium viscosity of 0.09 kPa·s, cortex viscosity of 11 kPa·s, and cell-cell friction of 0.12 kPa·s/$\mu$m). Five replicates (crm04-08) were used.

##### A.11.6 Tension and adhesion

fig. A49

fig. A48 fig. A47

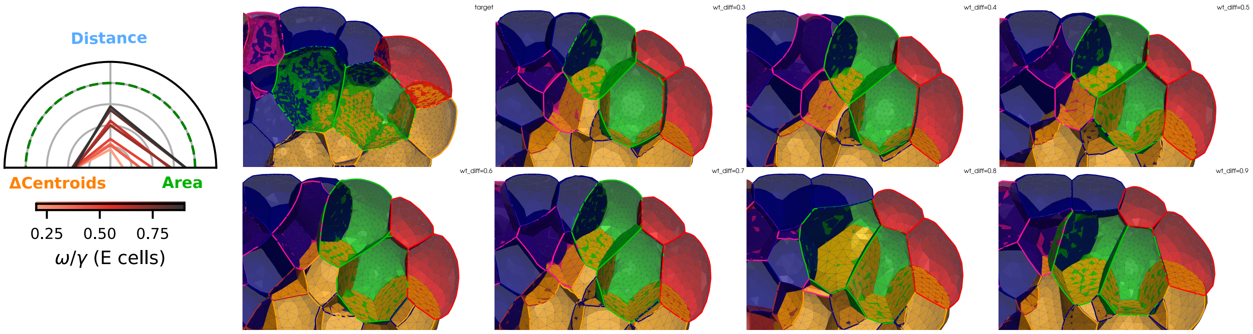


Figure A47: Effect of modulating adhesion energy of Ea and Ep on simulated ingression metrics. A constant apical tension of 3X surface tension was applied. Five replicates were used (crm04-08). The banner uses replicate crm04 to illustrate.

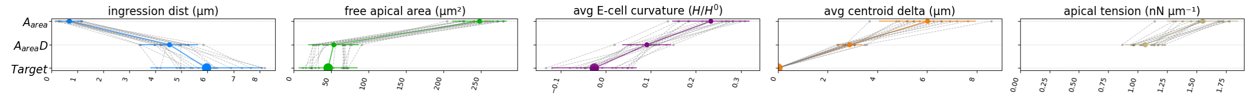


Figure A48: Effect of overall adhesion on simulated ingression metrics ($A_{abs}D - sweepAdh$). The y-axis represent the ratio between adhesive tension ($\omega$) and surface tension ($\gamma$), with a ratio of 0.5 being the base setting. A constant apical tension of 3X surface tension was applied.

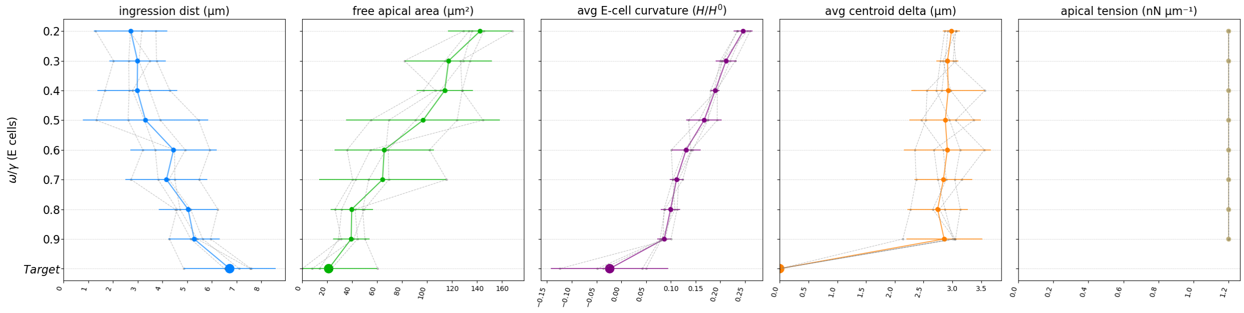


Figure A49: The effect of differential adhesion of the E-cells on ingression metrics ($A_{abs}D - sweepAdhE$). A constant apical tension of 3X surface tension was applied. The default ratio of adhesive tension ($\omega$) vs. surface tension ($\gamma$)is 0.5.

##### A.11.7 Anchoring of E-cells

fig. A50

These scenarios are designed to see what the effect is of the E-cells anchoring, either to one another (via the Ea-Ep anchor), or with the neighboring cells (clutch)

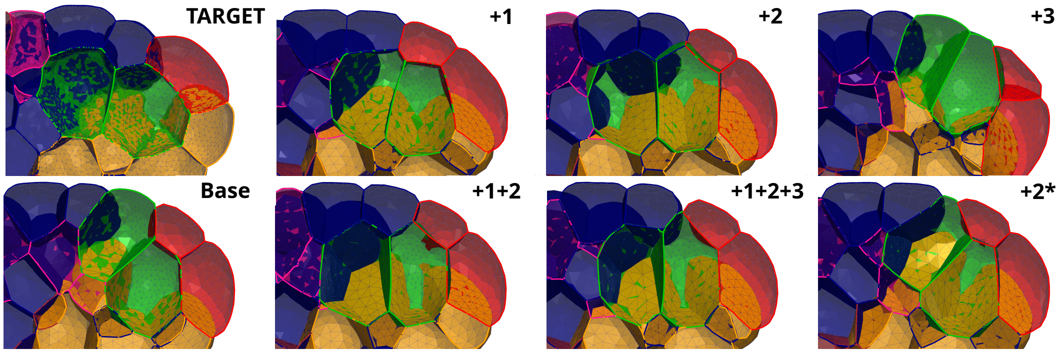


Figure A51

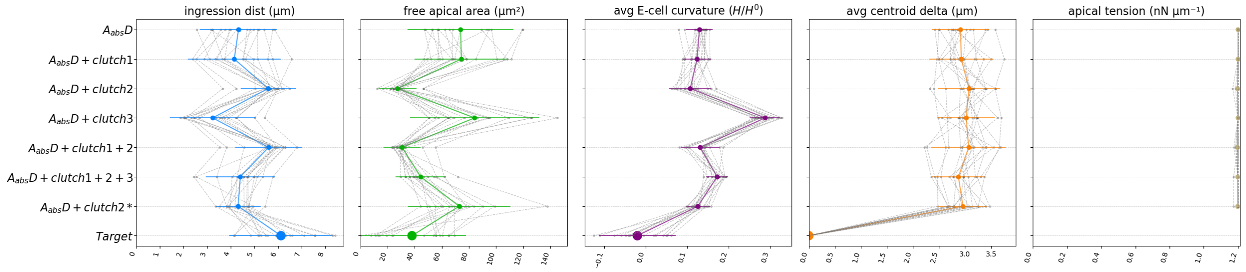


Figure A50: Effect of localized frictional clutches on ingression metrics. Simulations were performed with a constant absolute apical tension of $3\gamma_I$. The clutches are modeled as localized increases in cell-cell friction at specific interfaces using two approaches: a gradient friction is applied to the apical contact between the two E-cells (+1), scaling linearly from a maximum at the apical surface to zero at the cell centroid, whereas a uniform friction is applied to the apical ring contacting neighbors (+2) and the basolateral region (+3). A late-stage clutch (+2*) represents the uniform neighbor friction (+2) activated only during the final 10 minutes of the simulation.

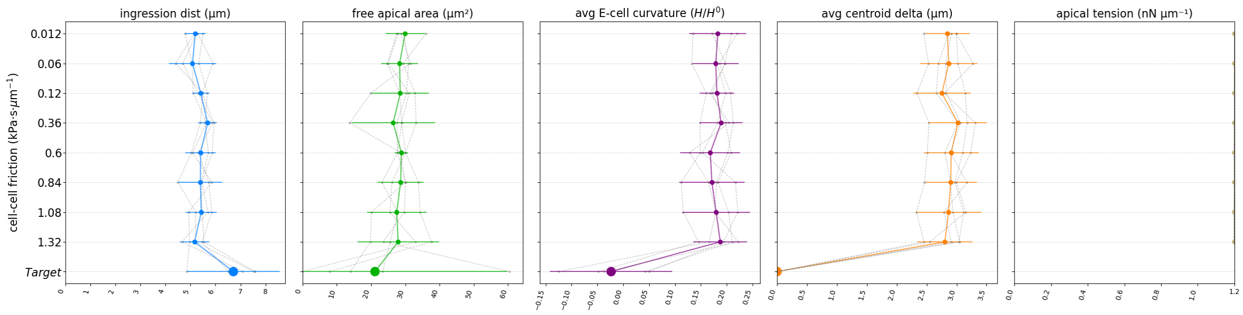


Figure A53: The effect of increasing global cell-cell friction on ingression metrics. An increased friction (1000 fold) is applied to the apical ring contacting neighbors ($A_{dist}D + clutch2$)

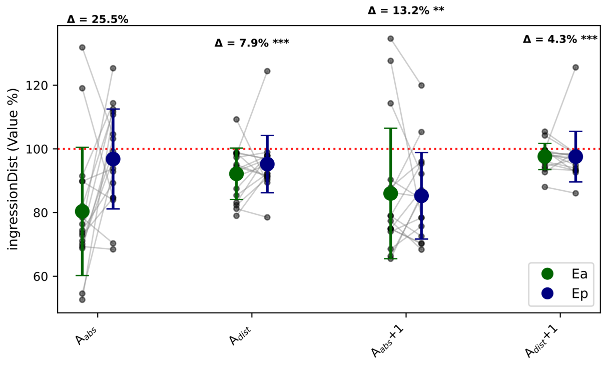


Figure A52: Comparison of ingression distance between Ea and Ep across different simulated scenarios. Each scenario on the x-axis displays the final ingression distance, represented as a percentage of the target value. Large green (Ea) and blue (Ep) dots indicate the mean performance across all replicates, with error bars showing the standard deviation. Small gray dots represent individual replicate values, connected by light gray lines to indicate the replicate cell pairs. The annotation $\Delta$ above each scenario indicates the mean of the absolute difference in distance between Ea and Ep cell for that scenario. Significance markers (* for $p < 0.05$, ** for $p < 0.01$) denote the result of a one-sided, paired t-test comparing the $\Delta$ of that scenario to the 'base' scenario (the first scenario on the left), indicating if the delta is significantly smaller. A smaller delta points to a more balanced ingression between the two E cells.

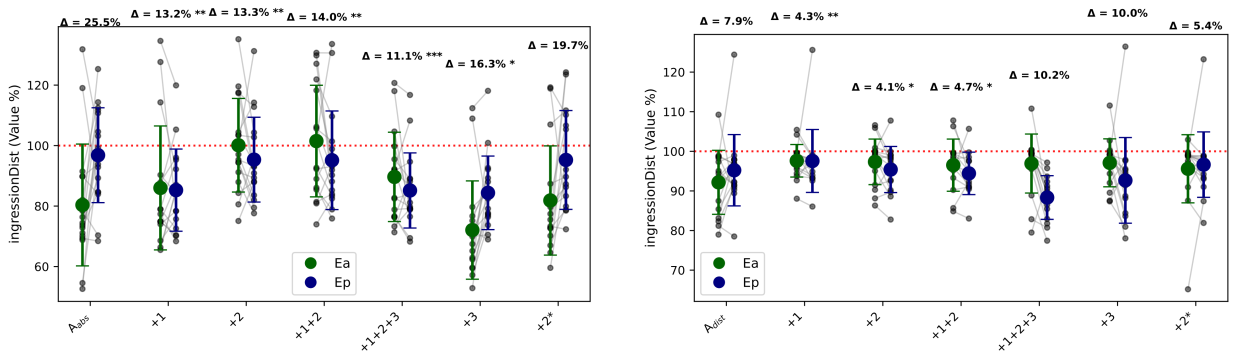


(a) Reference scenario: $A_{abs}$

(b) Reference scenario: $A_{dist}$

Figure A54: Comparison of ingression distance between Ea and Ep across different simulated clutch scenarios with either (a) a constant apical tension (base scenario = $A_{abs}D$) or (b) an apical tension tracking ingression distance (base scenario = $A_{dist}D$). In both plots, each scenario on the x-axis displays the final ingression distance, represented as a percentage of the target value. Large green (Ea) and blue (Ep) dots indicate the mean performance across all replicates, with error bars showing the standard deviation. Small gray dots represent individual replicate values, connected by light gray lines to indicate the replicate cell pairs. The annotation Δ above each scenario indicates the mean of the absolute difference in distance between Ea and Ep cell for that scenario. Significance markers (* for $p < 0.05$, ** for $p < 0.01$) denote the result of a one-sided, paired t-test comparing the Δ of that scenario to the respective 'base' scenario (the first scenario on the left), indicating if the difference between cells is significantly smaller. A smaller delta points to a more balanced ingression between the two E cells.

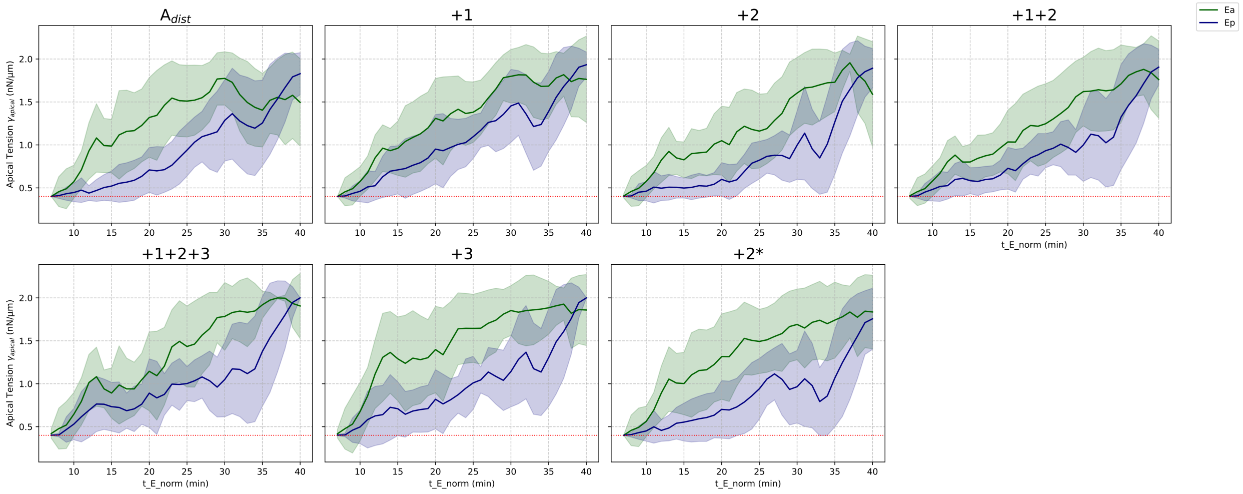


Figure A55: Comparison of simulation tension profiles of Ea and Ep across different clutch mechanism scenarios, all run with a apical tension tracking ingression distance (base scenario = $A_{\text{dist}}$). The shaded error bands represents ± 1SD from the mean across all simulation replicates. The red dotted line indicates the base tension of 0.4 nN $\mu$m$^{-1}$.

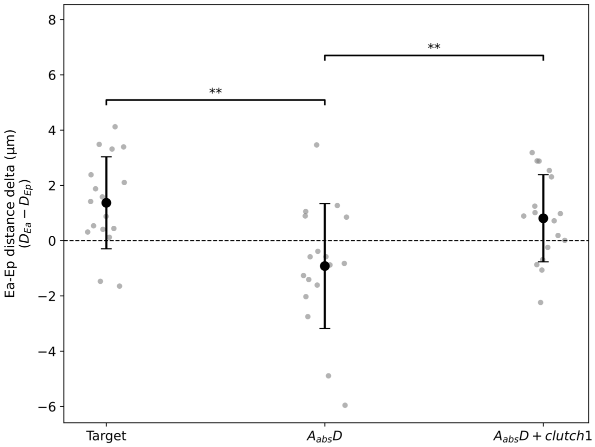


Figure A56: Comparison of the final ingression distance delta ($D_{Ea} - D_{Ep}$) between Ea and Ep across experimental data (Target) and simulation scenarios. Each gray dot represents a single replicate, while the black markers indicate the mean ± standard deviation for each scenario. Significance markers denote the result of a paired t-test between scenarios (*$p < 0.05$, **$p < 0.01$, ***$p < 0.001$). The horizontal dashed line at zero represents a perfect balance, where both cells ingress an equal distance.

#### A.12 Stokes flow simulation for visualizing collective cell movements

To assess and visualize the overall, collective movements of cells during E-cell ingression, we utilized Stokes flow simulations (fig. A59). This computational fluid dynamics (CFD) approach models the cellular environment as a highly viscous fluid, where inertial forces are negligible compared to viscous forces. The resulting 3D velocity field provides an approximation of the coordinated, flow-like behavior of the cell collective.

##### A.12.1 Input data: averaged cell displacements

The input for the Stokes flow simulation was derived from the observed cell movements, gathered from multiple replicates ($n = 39$). Normalization was performed with respect to the principal embryonic axes (appendix B.2), and the centroid positions were scaled to a common embryonic length scale, determined by averaging the extent of each principal axis across all replicates. Averaged 3D displacement vectors for each cell were then calculated over 1-minute intervals using linear interpolation, which served as the input forces within an OpenFOAM simulation.

##### A.12.2 Computational fluid dynamics (CFD) simulation setup

The collective cell behavior within the embryo was modeled as a slow, highly viscous, incompressible fluid flow, characteristic of the Stokes (or creeping) flow regime. This physical approximation is suitable for biological systems at the cellular scale, where viscous forces are dominant over inertial forces. The simulation aimed to determine the steady-state velocity field within the embryonic environment that results from the averaged cellular displacements.

A convex hull mesh encompassing all centroid positions was used as a proxy for the eggshell. On the eggshell boundary surface, a free-slip boundary condition was imposed. This condition signifies that the simulated cellular medium can move tangentially along the eggshell surface without frictional resistance, while preventing any flow across this boundary.

The mean displacement vectors of individual cells, determined from experimental observations as previously described, were translated into localized driving forces within the CFD simulation. These forces were applied to distinct regions within the mesh, corresponding to the initial average positions and volumes of the cells. The magnitude and direction of the applied force for each such region were derived from the respective cell's averaged displacement vector, scaled by its cell volume.

The governing equations for incompressible, laminar viscous flow were then solved numerically using OpenFOAM (version v2412) to obtain a steady-state 3D velocity field throughout the domain (fig. A59). This velocity field represents the emergent, collective motion of the cellular medium under the influence of the applied localized forces.

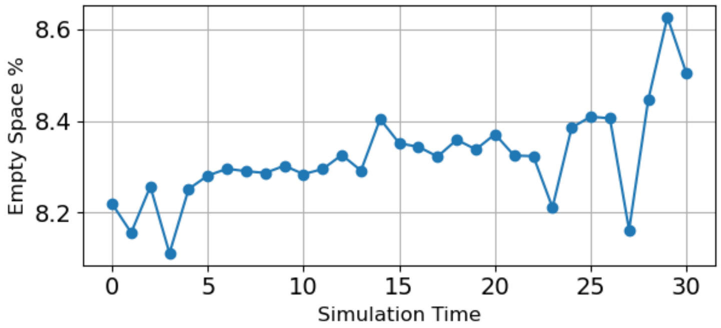


Figure A57: The evolution of the total empty space over the simulation timesteps using replicate crm04 as an example. Expressed as a percentage of the hull volume. This percentage stays within a narrow range of 0.5% [8.11, 8.62] for this replicate, which indicate a good approximation of incompressibility.

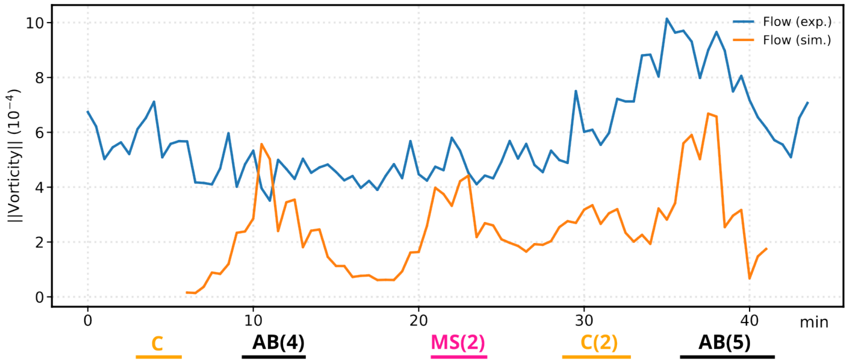


Figure A58: Mean curl magnitude of the velocity field (vorticity). The velocity flow field derived from in vivo centroid movements ($Flow_{exp}$, see appendix A.12) shows a distinct peak in vorticity (appendix B.5.5)during the late phase of ingression. In contrast, in the velocity flow field derived from centroid movements of the simulations ($Flow_{sim}$, orange), this peak is less pronounced. Simulation scenario $A_{dist}D + clutch1 + 2$ was used.

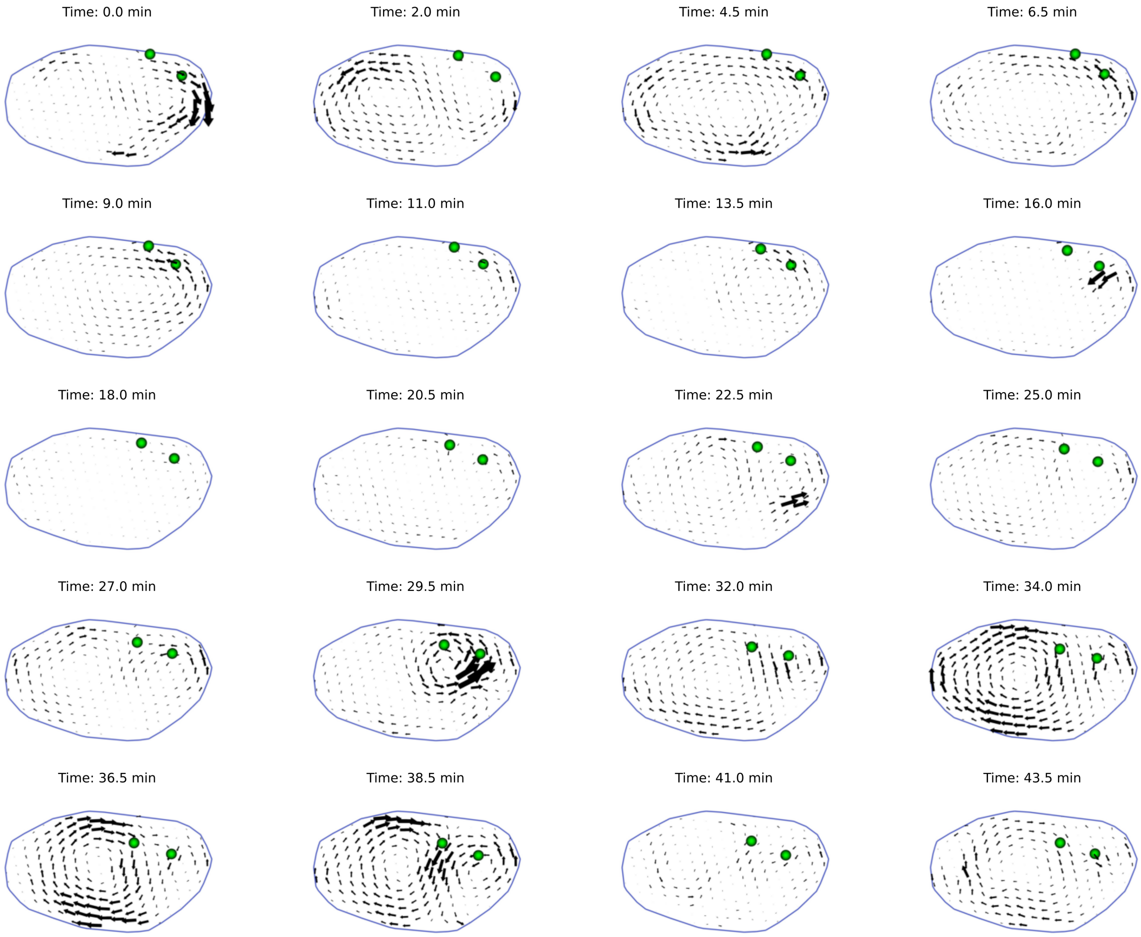


Figure A59: Visualization of the collective movements of cells during E-cell ingression by solving Stokes flow on the cell centroid displacements.

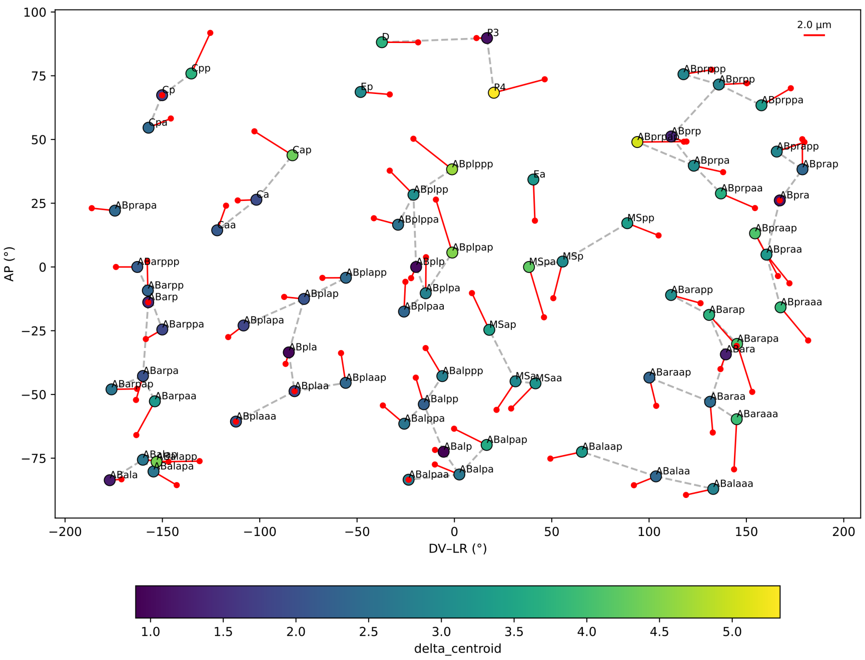


Figure A60: Visualization of the cell positioning error of the simulations. The average final 3D centroid of each cell from the simulation is projected onto a reference ellipsoid to generate 2D coordinates, where latitude corresponds to the AP axis and longitude to the DV-LR plane. The 2D map is an equirectangular projection onto an ellipsoid representing the eggshell. Each dot represents the mean final position of a cell, and its color indicates the magnitude of the displacement ($\Delta_{centroid}$ in µm) from its target position. The red lines illustrate this displacement vector; they originate from the cell's final position and point in the direction of its target position. The length of each line is scaled proportionally to the displacement magnitude, with a reference scale of 2.0 µm.

#### A.13 Supplementary videos

#### A.14 Data availability

All input data is available at https://zenodo.org/uploads/XXXXX.
