## Appendix B for "Integrated quantitative imaging and biomechanical modeling of early gastrulation in *C. elegans*"

### B Appendix-definitions/calculations

#### B.1 Mesh

A 3D mesh of an embryo is loaded from its corresponding VTP file. The mesh consists of:

$$\mathcal{V}_{\text{emb}} = \{\mathbf{v}_i \in \mathbb{R}^3 \,|\, i = 1, 2, \ldots, N_v\}, \quad \mathcal{T}_{\text{emb}} = \{T_j = (\mathbf{v}_{j1}, \mathbf{v}_{j2}, \mathbf{v}_{j3}) \,|\, j = 1, 2, \ldots, N_T\},$$

where $\mathcal{V}_{\text{emb}}$ represents the set of all vertices (nodes) of the embryo, and $\mathcal{T}_{\text{emb}}$ represents the set of all triangular faces in the mesh. A cell of interest is identified in the mesh using its parent index (`pid`):

$$\mathcal{V}_{\text{cell}} = \{\mathbf{v}_i \,|\, \texttt{pid}(\mathbf{v}_i) = \texttt{pid}_{cell}\}.$$

#### B.2 Coordinate systems for embryo and E-cells

For each replicate, two distinct coordinate systems are established: the embryonic axes (AP, DV, LR) and an axis system specific to the E-cell ingression process (Xe, Ye, Ze).

1. **Embryonic axes (AP, DV, LR)**: The embryonic axes define a global coordinate system for the entire embryo. In some cases minor manual adjustments were made using the E-division timepoint as a reference.

   - AP axis (Anterior-Posterior): The AP axis is defined as the line between the two points that are furthest apart in the XY-plane, projected through the embryo centroid. It points to the posterior side. The points are shifted such that the AP axis passes through the centroid:

   $$\text{AP} = \frac{\mathbf{p}_1 - \mathbf{p}_2}{\|\mathbf{p}_1 - \mathbf{p}_2\|}, \quad \text{with } \mathbf{p}_1, \mathbf{p}_2 = \arg\max\,(\text{distance in XY-plane}).$$

   - LR axis (Left-Right): The LR axis is aligned with the Z-direction ($\mathbf{e}_z$). It points to the right.
   - DV axis (Dorsal-Ventral): The DV axis is orthogonal to both the AP and LR axes, computed using the cross product:

   $$\text{DV} = \text{AP} \times \text{LR}.$$

   It points to the ventral side.

2. **Ingression axes (Xe, Ye, Ze)**: The ingression axes define a local coordinate system tailored to the ingression of the E-cells (fig. B1a). The axes are constructed using the centroids of specific cells:

   - $\mathbf{c}_{\text{Ea}}$, $\mathbf{c}_{\text{Ep}}$: Centroids of the cells Ea and Ep.
   - $\mathbf{c}_{\text{E4}}$: Mean centroid of the daughter cells (Eal, Ear, Epl, Epr) after division:

   $$\mathbf{c}_{\text{E4}} = \frac{\mathbf{c}_{\text{Eal}} + \mathbf{c}_{\text{Ear}} + \mathbf{c}_{\text{Epl}} + \mathbf{c}_{\text{Epr}}}{4}.$$

   - $\mathbf{c}_{\text{E0}}$: Projection of $\mathbf{c}_{\text{E4}}$ onto the Ea-Ep axis:

   $$\mathbf{c}_{\text{E0}} = \mathbf{c}_{\text{Ep}} + \frac{(\mathbf{c}_{\text{E4}} - \mathbf{c}_{\text{Ep}}) \cdot (\mathbf{c}_{\text{Ea}} - \mathbf{c}_{\text{Ep}})}{\|\mathbf{c}_{\text{Ea}} - \mathbf{c}_{\text{Ep}}\|^2}(\mathbf{c}_{\text{Ea}} - \mathbf{c}_{\text{Ep}}).$$

   Using these reference points, the axes are defined as:

   - Xe: Direction of ingression, from $\mathbf{c}_{\text{E0}}$ to $\mathbf{c}_{\text{E4}}$:

   $$\text{Xe} = \frac{\mathbf{c}_{\text{E4}} - \mathbf{c}_{\text{E0}}}{\|\mathbf{c}_{\text{E4}} - \mathbf{c}_{\text{E0}}\|}.$$

   - Ye: Direction from Ep to Ea:

   $$\text{Ye} = \frac{\mathbf{c}_{\text{Ea}} - \mathbf{c}_{\text{Ep}}}{\|\mathbf{c}_{\text{Ea}} - \mathbf{c}_{\text{Ep}}\|}.$$

   - Ze: Orthogonal to both Xe and Ye, ensuring a right-handed coordinate system:

   $$\text{Ze} = \text{Xe} \times \text{Ye}.$$

   To ensure Ze points leftward (based on ABplp orientation), it is flipped if necessary.

#### B.3 Ingression metrics

##### B.3.1 Ingression distance

The ingression distance quantifies the position of an E cell along the ingression axis ($X_e$):

1. The ingression distance is calculated by projecting the cell's centroid vector ($\mathbf{c}$) onto the ingression axis ($X_e$) after shifting the coordinate system origin to $\mathbf{E_0}$:

$$\text{IngressionDist} = (\mathbf{c} - \mathbf{E_0}) \cdot \mathbf{X_e}$$

2. A normalized ingression distance is also calculated by dividing the ingression distance by its value at the final time point:

$$\text{IngressionDistNorm} = \frac{\text{IngressionDist}}{\text{IngressionDist}_{\text{final}}}$$

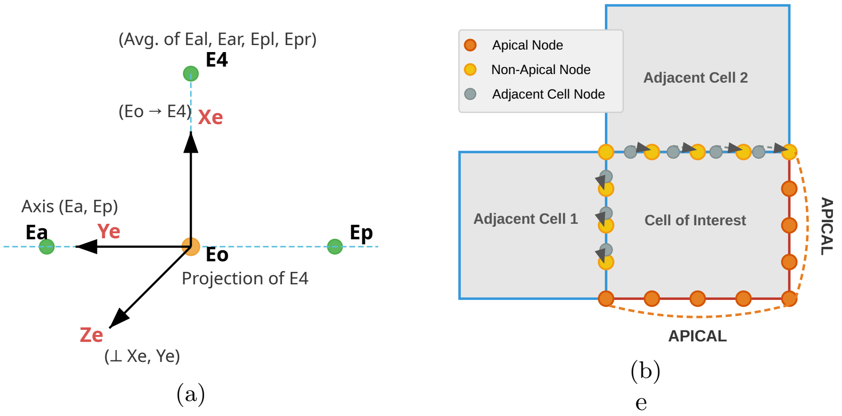


Figure B1: Schematic illustrations of ingression metric calculations. (a) Construction of the ingression-axes. (b) Determination of apical nodes for apical area calculation.

##### B.3.2 Apical area

The apical area represents the exposed surface of a cell, not in contact with neighboring cells (fig. B1b).

1. Apical nodes are determined as follows:

   (a) For each node in the embryo ($\mathbf{v}_k \in \mathcal{V}_{\text{emb}}$), find its nearest neighbor among the cell's nodes ($\mathcal{V}_{\text{cell}}$):

$$\text{NN}(\mathbf{v}_k) = \arg\min_{\mathbf{v}_j} \|\mathbf{v}_k - \mathbf{v}_j\|, \quad \forall\, \mathbf{v}_j \in \mathcal{V}_{\text{cell}}.$$

   (b) Exclude any nodes in $\mathcal{V}_{\text{cell}}$ that serve as the nearest neighbor for nodes outside the cell:

$$\mathcal{V}_{\text{apical}} = \mathcal{V}_{\text{cell}} \setminus \{\mathbf{v}_j \,|\, \exists\, \mathbf{v}_k \in (\mathcal{V}_{\text{emb}} \setminus \mathcal{V}_{\text{cell}}),\, \text{NN}(\mathbf{v}_k) = \mathbf{v}_j\}.$$

2. Triangular faces are considered apical if all three of their vertices are apical. Let $T_j = (\mathbf{v}_{j1}, \mathbf{v}_{j2}, \mathbf{v}_{j3})$, then:

$$T_j \in \mathcal{T}_{\text{apical}} \quad \text{if} \quad \mathbf{v}_{j1}, \mathbf{v}_{j2}, \mathbf{v}_{j3} \in \mathcal{V}_{\text{apical}}.$$

3. The apical area is computed as the sum of the areas of all triangles in $\mathcal{T}_{\text{apical}}$:

$$A_{\text{apical}} = \sum_{T_j \in \mathcal{T}_{\text{apical}}} \text{Area}(T_j)$$

##### B.3.3 Curvatures

The curvature metric quantifies the mean curvature of an E-cell's surface, normalized to a reference value.

1. **Curvature definition**: The local curvature, $H$, at each mesh triangle is computed as the mean of the principal curvatures:
$$H = \frac{\kappa_1 + \kappa_2}{2}.$$

2. **Filtering cell-cell interfaces**: Within a cell-cell interface, only triangles close to the center of the contact area are selected for curvature calculation.

   - The center triangle is determined based on the pairwise distances between all triangles in the interface. The triangle with the minimum maximum distance to all other triangles is chosen as the center:
   $$T_{\text{center}} = \arg \min_{T_i \in \mathcal{T}_{\text{interface}}} \max_{T_j \in \mathcal{T}_{\text{interface}}} \|\mathbf{x}_i - \mathbf{x}_j\|,$$
   where $\mathbf{x}_i$ and $\mathbf{x}_j$ are the centroid positions of triangles $T_i$ and $T_j$.
   - A heuristic threshold (e.g., 40% of triangles closest to $T_{\text{center}}$) is applied to exclude peripheral triangles. The selected triangles, $\mathcal{T}_{\text{selected}}$, are used for further calculations.

3. **Centering curvatures at the interface level**: The curvature of each cell-cell interface is "centered" relative to the mean curvature of the contacting cell at that interface:
$$H_{\text{centered}} = H - \frac{H + H_{\text{contact}}}{2},$$
where $H_{\text{contact}}$ is the mean curvature of the interface triangles belonging to the contacting cell.

4. **Area-weighted average for cell-cell interfaces**: For each cell-cell interface, the centered curvature is averaged using an area-weighted approach:
$$H_{\text{interface}} = \frac{\sum_{T_i \in \mathcal{T}_{\text{selected}}} H_{\text{centered},i} \cdot A_i}{\sum_{T_i \in \mathcal{T}_{\text{selected}}} A_i},$$
where $H_{\text{centered},i}$ is the centered curvature of triangle $T_i$, and $A_i$ is its area.

5. **Averaging for E-cells**: The curvature values for all interfaces associated with a given E-cell are averaged to produce a single value for each E-cell:
$$H_{\text{avg}} = \frac{1}{|\mathcal{I}_c|} \sum_{I \in \mathcal{I}_c} H_{\text{interface}},$$
where $\mathcal{I}_c$ is the set of interfaces involving the E-cell.

6. **Normalization**: The average curvature is normalized by a reference value ($H_0 = 10^5$), which corresponds to the curvature of a sphere with a radius of 10 micron:
$$H_{\text{norm}} = \frac{H_{\text{avg}}}{H_0}.$$

##### B.3.4 Sphericity

The sphericity metric quantifies the geometric similarity of a cell's shape to that of a perfect sphere. It is defined as:

$$\Psi = \frac{\pi^{\frac{1}{3}} \cdot (6 \cdot V)^{\frac{2}{3}}}{A},$$

where:

- $V$: The volume of the cell.
- $A$: The surface area of the cell.

The sphericity ranges from 0 to 1, where a value of 1 corresponds to a perfect sphere.

##### B.3.5 Avg centroid delta

The centroid delta metric quantifies the positional deviation of cells in simulated meshes relative to their corresponding positions in target meshes. It is computed as follows:

1. For each cell in the simulated mesh, the Euclidean distance ($\Delta$) between the cell's position ($\mathbf{c}_{\text{sim}}$) and its position in the target mesh ($\mathbf{c}_{\text{target}}$) is calculated:

$$\Delta = \|\mathbf{c}_{\text{sim}} - \mathbf{c}_{\text{target}}\|.$$

2. The target position, $\mathbf{c}_{\text{target}}$, is interpolated from the target meshes at the query time from E-division ($t_E$). Specifically:

   - The target meshes at timepoints immediately before and after $t_E$ are identified.
   - Linear interpolation is performed between these timepoints to estimate the cell's position at $t_E$:

$$\mathbf{c}_{\text{target}}(t_E) = \frac{t_{\text{after}} - t_E}{t_{\text{after}} - t_{\text{before}}}\mathbf{c}_{\text{before}} + \frac{t_E - t_{\text{before}}}{t_{\text{after}} - t_{\text{before}}}\mathbf{c}_{\text{after}},$$

   where $t_{\text{before}}$ and $t_{\text{after}}$ are the times of the surrounding target meshes.

3. The delta centroid for the embryo mesh from simulation is calculated as the average delta distance across all cells:

$$\Delta_{\text{avg}} = \frac{1}{N_c}\sum_{i=1}^{N_c}\Delta_i,$$

   where $N_c$ is the total number of cells in the embryo.

##### B.3.6 Visualisation of ingression metrics

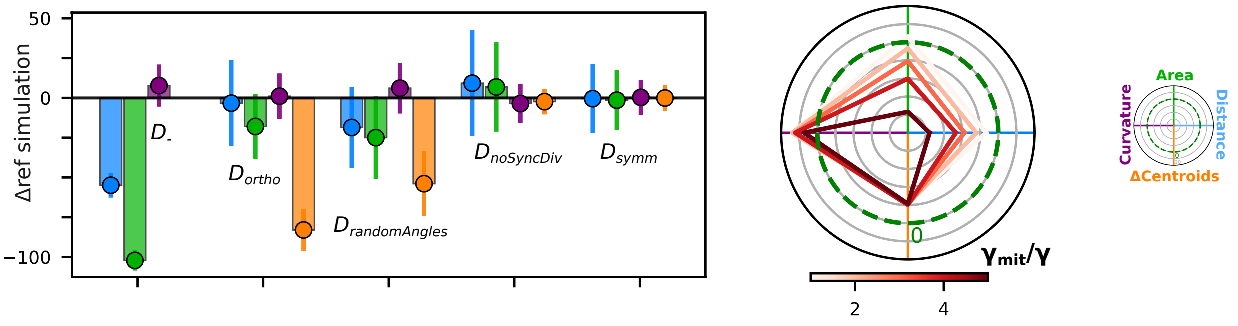


Figure B2: Bar chart and radar plot example

The primary methods for visualizing simulation metrics involve bar charts and radar plots (fig. B2), both designed to represent deviations from target values, reference ranges, or reference simulations. A core representation for several metrics, including `ingression dist`, `free apical area`, and `avg centroid delta`, is a percentage based on their progression through a defined range, expressed as $Y = P_v - 100\%$ where $Y$ is the value on the bar chart's y-axis or the radial value on the radar plot. The percentage $P_v$ is calculated as:

$$P_v = \frac{v_{sim} - v_{first}}{v_{last} - v_{first}} \times 100\%$$

where $v_{sim}$ is the simulated metric value. The terms $v_{first}$ and $v_{last}$ define the start and end points of the target reference range, specific to each metric:

- For `ingression dist` and `free apical area`: $v_{first}$ is the target value of the metric at the initial simulation timepoint for a given replicate, and $v_{last}$ is the target value at the final simulation timepoint for that replicate.

- For `avg centroid delta`: The reference range is determined for each cell and is the AB-distance from start to end position. Daughter cells are traced back to their ancestral cells to do so (for example, in case of P4 the delta centroid is the difference between the final position of P4 to the position of P3 at the start of the simulation). $v_{last}$ is set to 0 per definition. $v_{sim}$ is the delta centroid as an absolute value. In other words the remaining displacement error is expressed relative to the AB-trajectory distance of the cell. This value is then averaged over all cells to the final result.

For these metrics, a plotted value of 0 (i.e., $P_v = 100\%$) signifies that the simulated metric value $v$ has reached its target value perfectly. For other metrics, which do not have such a defined target start and end value, we need other representations:

- **Average E-cell curvature**: This metric quantifies the deviation of the simulated mean curvature ($H_{sim}$) from the target mean curvature ($H_{target}$). It is calculated as the scaled difference: $(H_{sim} - H_{target})/s$. A plotted value of 0 indicates that $H_{sim} = H_{target}$. The scaling factor used for this metric is $s = 5 \times 10^{-5}$.
- **Apical tension**: Only shown in radar plots in context of parameter sweeps, these metrics are visualized as a percentage difference of the simulated value $v$ from the value of a reference run within that sweep: $P = \frac{v - v_{ref}}{v_{ref}} \times 100\%$. For visual clarity, the radial y-bars represent 5%. A radial value $R = 0$ means $v = v_{ref}$.

For the bar chart representation:

- Error bars typically show the standard deviation. For metrics visualized as $P_v - 100\%$, this is the standard deviation of the $P_v$ values. For `avg E-cell curvature`, it's the scaled standard deviation of the $(H_{sim} - H_{target})$ difference.
- If a reference scenario is specified, all plotted $Y$ values are further adjusted by subtracting the $Y$ value of the designated reference scenario. This allows for direct visual comparison of deviations relative to this reference scenario.

##### B.3.7 Ingression volume flux

The ingression volume flux quantifies the rate of change in occupancy by E-cells (Ea and Ep) of a pre-defined target volume, corresponding to the space occupied by Ea and Ep at the end of ingression, right before they divide. This metric provides an alternative way to describe the ingression dynamics of the E-cells, expressed as a percentage of volume change per minute (%vol/min).

1. **Defining the reference E-cell target volume**:
   - A reference timepoint, $t_{ref}$, is chosen, typically corresponding to the last imaging stack where both Ea and Ep cells are present and have reached their approximate final positions post-ingression. The combined mesh of Ea and Ep cells at this $t_{ref}$ defines the target volume.
   - A large number of sample points, $\mathcal{P}_{Ex} = \{\mathbf{p}_k\}$, are uniformly distributed within this target volume derived from Ea and Ep at $t_{ref}$. Let $N_{total\_points} = |\mathcal{P}_{Ex}|$.
2. **Tracking occupancy of the target volume over time**: For each timepoint $t_j$:
   - For every cell $c$ present in the embryo at time $t_j$, its 3D mesh $\mathcal{M}(c, t_j)$ is obtained.
   - The number of sample points from $\mathcal{P}_{Ex}$ that fall within the mesh $\mathcal{M}(c, t_j)$ is determined. This is denoted as $N_{inside}(c, t_j)$.
   - The fractional overlap for cell $c$ with the target volume is calculated as:

$$O(c, t_j) = \frac{N_{inside}(c, t_j)}{N_{total\_points}}.$$

- This fractional overlap is then normalized by the sum of overlaps of all cells at that specific timepoint $t_j$, to account for slight discretization errors:

$$O_{norm}(c, t_j) = \frac{O(c, t_j)}{\sum_{all\_cells\, i} O(i, t_j)}.$$

3. **Calculating volumetric flux for E-cells**:

- At each timepoint $t_j$, the total normalized overlap by the E-cells (Ea and Ep) is the sum of their individual normalized overlaps:

$$O_{E,tot}(t_j) = O_{norm}(Ea, t_j) + O_{norm}(Ep, t_j).$$

This value represents the fraction of the initial target E-cell space that is occupied by the current Ea and Ep cells at time $t_j$.

- The volumetric flux, $\text{Flux}_E(t_j)$, at time $t_j$ is computed using a central difference method, considering the timepoints $t_{j-1}$ (stack before $t_j$) and $t_{j+1}$ (stack after $t_j$). Let $\Delta t_{stack}$ be the time interval between consecutive imaging stacks (e.g. 1.5 minutes).
- The change in total normalized E-cell overlap over the time period $2 \cdot \Delta t_{stack}$ is:

$$\Delta O_{E,tot}(t_j) = O_{E,tot}(t_{j+1}) - O_{E,tot}(t_{j-1}).$$

- The ingression volume flux is then calculated as the rate of this change, expressed as a percentage per minute:

$$\text{Flux}_E(t_j) = \frac{\Delta O_{E,tot}(t_j)}{2 \cdot \Delta t_{stack}} \times 100\%.$$

#### B.4 Angle calculations

##### B.4.1 Division angles

Two primary angles are computed: the apical angle (fig. B3a) and the AP-angle (fig. B3b). The calculation proceeds as follows:

1. **Reference geometry and division vector**:

- A reference ellipsoid is computed to approximate the embryo's surface, based on the bounding box of cell centroids. Let its center be $\mathbf{c}_{ellipsoid}$ and its semi-axes be $a, b, c$ corresponding to the AP, DV, and LR directions respectively.
- For a given dividing parent cell, its centroid $\mathbf{c}_{parent}$ (normalized AP, DV, LR coordinates) is identified (fig. B3a).
- This centroid is projected onto the surface of the reference ellipsoid by finding the closest point, resulting in $\mathbf{p}_{proj}$.
- The outward surface normal $\mathbf{N}$ of the ellipsoid is calculated at $\mathbf{p}_{proj}$.
- The division direction $\mathbf{d}$ is determined from the angles of the division normal (normal to the division plane) with respect to the global embryonic axes (AP, DV, LR degrees).

2. **Apical angle ($\alpha_{\text{apical}}$)**: The apical angle is defined as the angle between $\mathbf{d}$ and $\mathbf{N}$:

$$\alpha_{\text{apical}} = \arccos\left(\frac{\mathbf{d} \cdot \mathbf{N}}{\|\mathbf{d}\| \|\mathbf{N}\|}\right).$$

The orientation of $\mathbf{d}$ is chosen so that $\alpha_{\text{apical}}$ falls in the range $[0°, 90°]$.

- An $\alpha_{\text{apical}} \approx 0°$ indicates a division that is largely perpendicular to the surface (i.e., radial).
- An $\alpha_{\text{apical}} \approx 90°$ indicates a division that is largely parallel to the surface (i.e., tangential).

3. **AP angle (derived from $\phi_{\mathbf{azimuth}}$)**: This angle describes the orientation of the tangential component of the division vector relative to the embryo's AP axis (fig. B3b).

   - The division vector $\mathbf{d}$ is projected onto the tangent plane defined by the normal $\mathbf{N}$ at $\mathbf{p}_{proj}$. Let this tangential component be $\mathbf{d}_{\text{tan}}$.
   - A reference direction (the 0°-line) in the tangent plane, $\mathbf{AP}_{\text{tan}}$, is established, corresponding to the projection of the AP axis onto this tangent plane.
   - The azimuthal angle, $\phi_{\text{azimuth}}$, is the angle between $\mathbf{AP}_{\text{tan}}$ and $\mathbf{d}_{\text{tan}}$, measured within the tangent plane, defined in a range $[-180°, 180°]$. A value of 0° means $\mathbf{d}_{\text{tan}}$ aligns with the posterior direction, ±90° means it aligns with the DV/LR direction in the tangent plane, and ±180° means it aligns with the anterior direction.
   - The AP angle is a folded version of this azimuthal angle:

$$\phi_{\text{AP}} = \left| \phi_{\text{azimuth}} - 180° \cdot \text{round}\left(\frac{\phi_{\text{azimuth}}}{180°}\right) \right|.$$

     This transformation maps the $\phi_{\text{AP}}$ range to $[0°, 90°]$:
     - $\phi_{\text{AP}} \approx 0°$ indicates that $\mathbf{d}_{\text{tan}}$ is primarily aligned along the AP axis (either anteriorly or posteriorly).
     - $\phi_{\text{AP}} \approx 90°$ indicates that $\mathbf{d}_{\text{tan}}$ is primarily aligned orthogonally to the AP axis (i.e., along the DV or LR direction within the tangent plane).

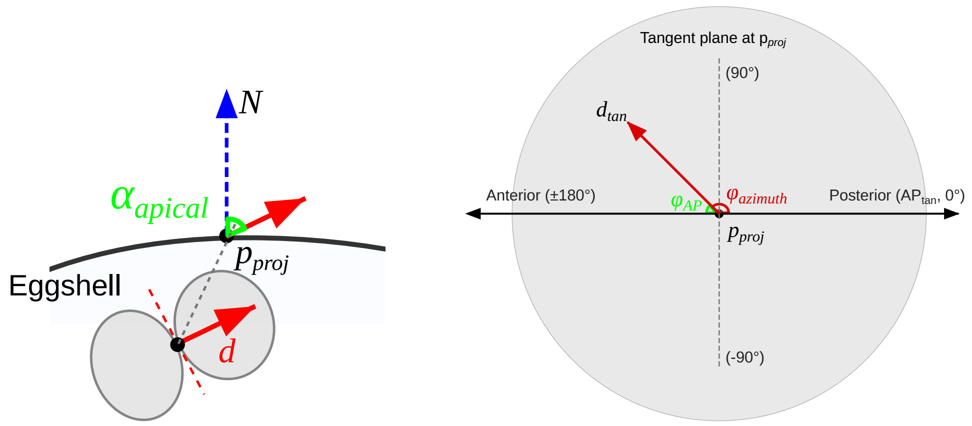


(a) Apical angle

(b) Azimuth angle

Figure B3: Division angles illustration

##### B.4.2 Triple junction angles

Triple junction angle calculations are performed as in Vanslambrouck et al. (2024) which is based on the algorithm described in Xu et al. (2018). Where three cells meet, or two cells and the exterior, a triple junction is defined (fig. B4). To measure the angles of a triple junction, the triangles from the cell meshes are first reassigned to three distinct surfaces. At several points along the junction, we then measure the angle between the surfaces in a plane normal to the contact line. The triple junction angles are averaged along the junction to give us the three final angles $\theta_{ij}$. Below, we explain this in more detail, step-by-step:

1. **Definitions**:

   - **Trijunction**: Trijunctions are points where three cells meet. One of the "cells" can also be the outside environment, in that case the trijunction is referred to as an exterior trijunction, as opposed to interior.

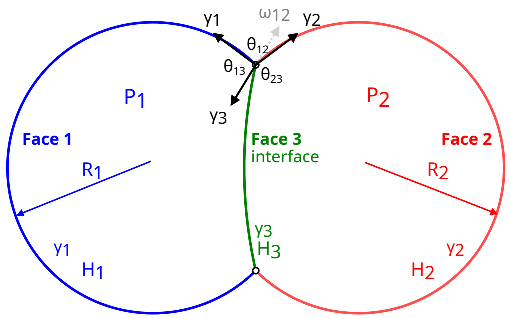


Figure B4: A diagram explaining the mechanical principles at a triple junction, the point where three interfaces meet. For the system to be in mechanical equilibrium, the tensional forces acting along each interface must balance. The contact angles ($\theta$), such as $\theta_{13}$ and $\theta_{23}$, describe the geometry of the force balance and provide information on the relative surface tensions, shown as $\gamma_1$, $\gamma_2$, and $\gamma_3$. An adhesive tension ($\omega_{12}$) can exist between two cells, which reduces the tension at their shared interface. The shape of the interface is governed by the Young-Laplace equation ($P_1 - P_2 = 2H_3\gamma_3$). For a spherical surface, the mean curvature ($H$) is the reciprocal of its radius ($R$), such that $H = 1/R$ (*adapted from* Vanslambrouck et al. (2024))

- **Trijunction angle**: A trijunction contains three angles, which add up to 360 degrees. The angle which is outlined by cell A, will be reffered to as "angle A"

2. **Determination of the trace line**: The trace line is the curve where the three surfaces meet, and a set of trijunctions is selected along this line. This trace line also provides the projection plane used for reading out the trijunction angles. The final trijunction angles for a given trijunction will be the averaged angles of this set. To compute the trace line:

   (a) Candidate points are sampled from the centroids of the triangles at the trijunction and their midpoints. These points are collected into a set:

   $$\mathbf{x}_{\text{trace}} = \{\mathbf{x}_i\} \quad \forall i \in \mathcal{T}_{\text{junction}}.$$

   (b) A k-d tree is used to find neighboring points within a characteristic length $L_c$, defined as:

   $$L_c = \sqrt{\frac{\sum A_i}{N}}, \quad A_i = \text{area of triangle } i, \quad N = \text{number of triangles}.$$

   (c) Points are retained for the trace line if they are near all three surfaces at the junction, ensuring proximity to all parent cells. The neighborhood radius for validation is set to $1.5L_c$.

3. **Calculating trijunction angles**: The angles at the trijunction are calculated by projecting the triangle normals onto the normal plane of the trace line and measuring the angles between the tangents:

   $$\theta_{ij} = \arccos\left(\frac{\mathbf{t}_i \cdot \mathbf{t}_j}{\|\mathbf{t}_i\|\|\mathbf{t}_j\|}\right),$$

   where $\mathbf{t}_i$ and $\mathbf{t}_j$ are the tangents of the surfaces corresponding to two of the meeting cells. These tangents are derived by projecting the surface normals onto normal plane of the trace line.

4. **Validation and filtering**: Several filtering criteria are applied to ensure robust and accurate trijunction angle calculations:

   - **Minimum triangle count**: A trijunction must involve at least 50 triangles ($N_{\text{triangles}} \geq 50$) for validation.
   - **Trace line sampling**: Angles are sampled along the trace line, requiring at least 10 points ($N_{\text{samples}} \geq 10$).

- **Exclusion of faulty triangles**: Triangles classified as "exterior" but with fewer than 10 connected neighbors are excluded, as they likely represent isolated artifacts.
- **Geometric properties**: Triangles with areas below 0.01 times the average contact area between cells are excluded:

$$A_i < 0.01 \cdot \overline{A}_{\text{contact}}, \quad \overline{A}_{\text{contact}} = \frac{\sum A_{\text{contact}}}{N_{\text{contacts}}}.$$

5. **Output**: For each trijunction, the method provides the three interior angles $(\theta_1, \theta_2, \theta_3)$ corresponding to the three cells meeting at the junction.

#### B.5 Protein distribution

##### B.5.1 Center of mass distance ($\mathbf{prot_{com}}$)

The center of mass distance ($prot_{com}$) quantifies the spatial offset between the centroid of a surface and its intensity-weighted center of mass. It is defined as:

$$prot_{com} = \left\| \frac{\sum_{i=1}^{N} \mathbf{x}_i \cdot I_i}{\sum_{i=1}^{N} I_i} - \frac{\sum_{i=1}^{N} \mathbf{x}_i}{N} \right\|,$$

where:

- $\mathbf{x}_i = (x_i, y_i, z_i)$ are the mesh node coordinates,
- $I_i$: The normalized protein signal intensity at point $i$.
- $N$: The total number of points on the surface.

A high $prot_{com}$ indicates a significant asymmetry in the protein distribution, while a low $prot_{com}$ suggests a more uniform distribution.

**Normalization:**

The normalization of $prot_{com}$ (fig. B6) is a two-step process designed to handle the distribution's skewness and scale. First, a logarithmic transformation is applied to the raw values. Subsequently, the resulting log-transformed values are standardized.

1. **Logarithmic transform**: A small constant, $\epsilon = 10^{-2}$, is added to each value to prevent issues with non-positive numbers before applying the natural logarithm:

$$\mathbf{prot}_{\text{com, log}} = \log(\mathbf{prot}_{\text{com, raw}} + \epsilon)$$

2. **Standardization**: The log-transformed values are then centered and scaled:

$$\mathbf{prot}_{\text{com}} = \frac{\mathbf{prot}_{\text{com, log}} - \mu_{\text{log}}}{\sigma_{\text{log}}}$$

##### B.5.2 Moment of inertia ($\mathbf{prot_{moi}}$)

The function $prot_{moi}$ indicates how the observed protein intensities ($I_i$) are spatially distributed around their centroid, relative to a uniform (average-intensity) distribution.

$$prot_{moi} = \frac{\sum_{i=1}^{N} \left\| \mathbf{x}_i - \overline{\mathbf{x}} \right\|^2 I_i}{\sum_{i=1}^{N} \left\| \mathbf{x}_i - \overline{\mathbf{x}} \right\|^2 \overline{I}} \tag{4}$$

**Where:**

- $\mathbf{x}_i = (x_i, y_i, z_i)$ are the mesh node coordinates,
- $\overline{\mathbf{x}} = \frac{1}{N} \sum_{i=1}^{N} \mathbf{x}_i$ is the centroid of the cell-cell interface,

- $I_i$ is the normalized protein intensity at point $i$,
- $\overline{I} = \frac{1}{N}\sum_{i=1}^{N} I_i$ is the average intensity,
- $N$ is the total number of points on the surface.

**Interpretation:**

$prot_{moi} < 1 \implies$ intensities are more concentrated toward the center.
$prot_{moi} > 1 \implies$ intensities are more spread out from the center.

**Normalization:**

The normalization of $prot_{moi}$ (fig. B6) is performed using a quantile transformation, which maps the distribution of the data to a standard normal (Gaussian) distribution. This non-linear transformation is robust to outliers and effectively reshapes the data distribution. The procedure is implemented using the 'QuantileTransformer' method from the 'scikit-learn' library.

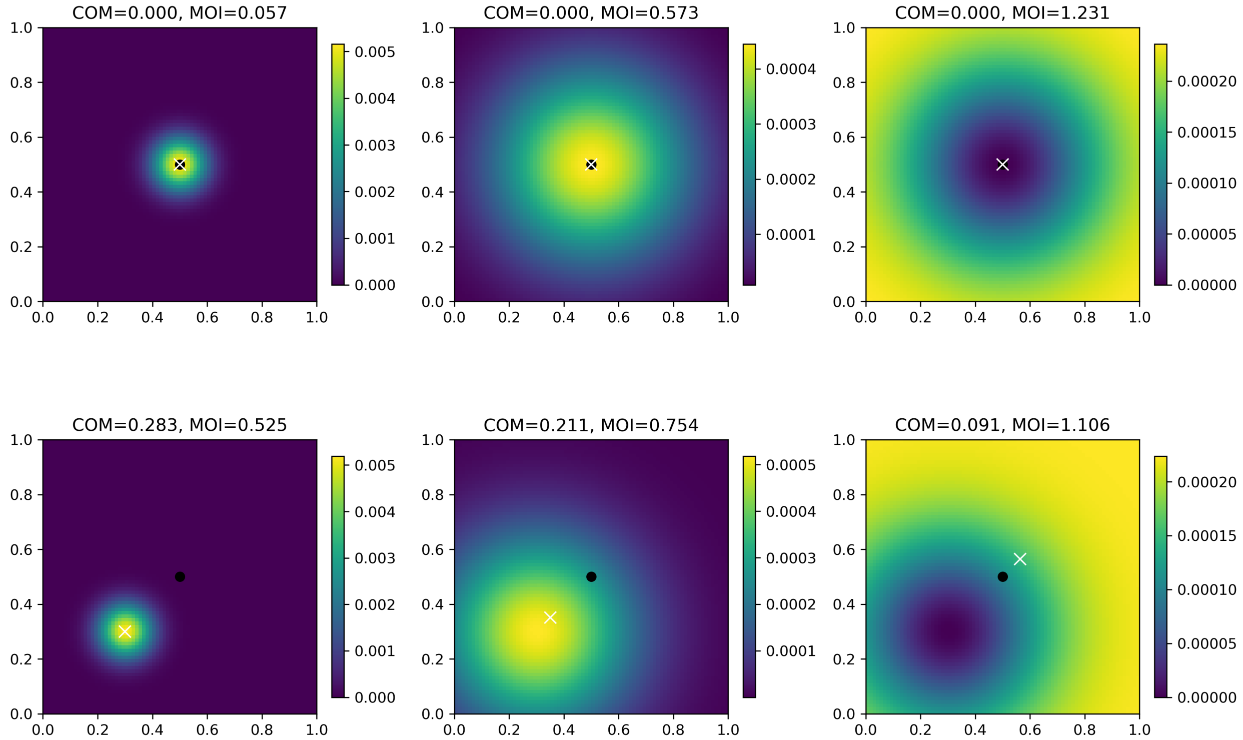


Figure B5: Illustration of $\mathbf{prot}_{\mathrm{com}}$ and $\mathbf{prot}_{\mathrm{moi}}$ scenarios. All surfaces have equal amount of protein signal, only the distribution changes.

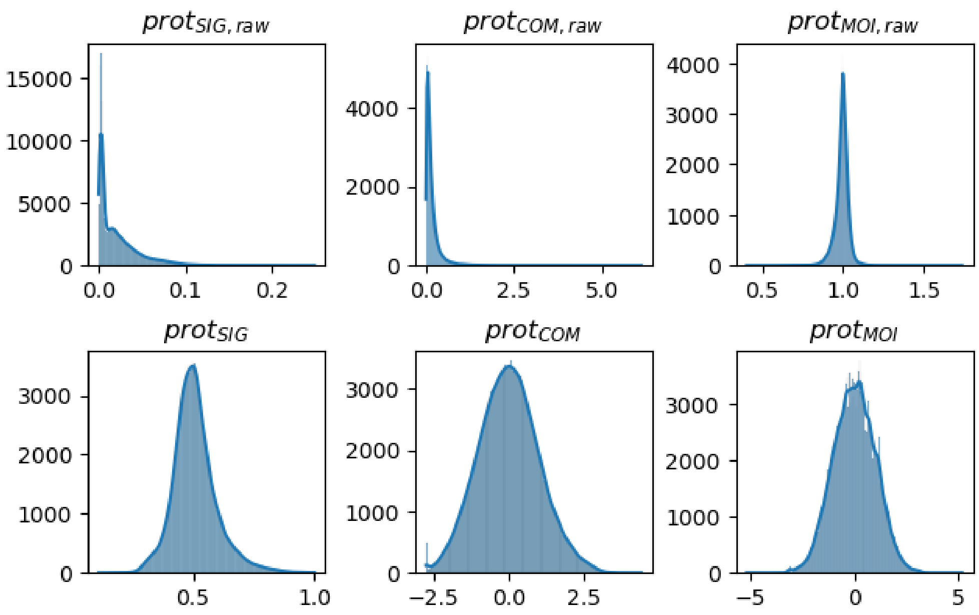


Figure B6: Distribution of raw and normalized protein surface metrics. The top row displays the raw distributions for signal ($prot_{sig,raw}$), center of mass ($prot_{com,raw}$), and moment of inertia ($prot_{moi,raw}$). The bottom row shows the corresponding normalized metrics.

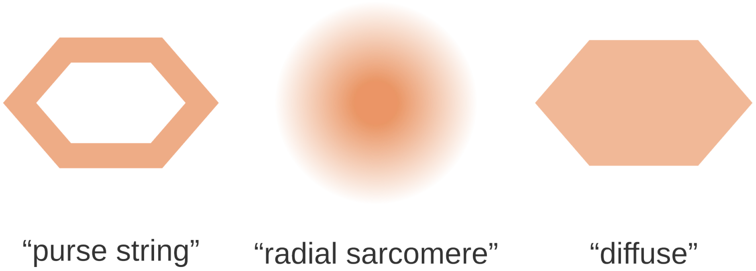


Figure B7: Models of actomyosin architecture that drive apical constriction in cells, with the orange shading representing the regions where myosin-based force generation is localized. The "purse string" model consists of an apicolateral or circumferential actomyosin belt that generates pulling forces at the periphery of the cell apex. The "radial sarcomere" model shows a medioapical network with centrally enriched non-muscle myosin II, as observed in the *Drosophila* ventral furrow. The "diffuse" model represents a medioapical network where non-muscle myosin II is distributed throughout the apical cortex without a central bias, as seen during *C. elegans* gastrulation. *Adapted from* Zhang et al. (2023).

##### B.5.3 Protein profiles

A protein profile describes the spatial distribution of a cortical protein (E-cadherin, myosin, or actin) across a specific cell surface. This surface may correspond to a cell-cell interface or a free apical area of a cell. The profile is computed by projecting protein signal intensities associated with the mesh nodes along a specific directional axis, binning the distances along that axis, and averaging the protein signal within each bin.

1. **Calculation of node distances**: The protein profile is computed from node distances extracted from the triangulated mesh of the cell or cell-cell interface. It is computed along specific directional axes, such as:
   - **Apical distance**: From the basal to the apical region of the cell or cell-cell interface.

- **Eap distance**: Toward the apical region of the Ea-Ep contact area.

For apical distances, the following steps are performed:

- The convex hull of the embryo mesh is computed.
- The distance ($d_i$) from each mesh node to the convex hull is calculated:

$$d_i = \|\mathbf{x}_i - \mathbf{p}_i\|,$$

where $\mathbf{x}_i$ is the position of node $i$, and $\mathbf{p}_i$ is the closest point on the convex hull.

After distances are computed, per cell-cell interface, they are shifted such that the most apical node corresponds to a distance of zero:

$$d_i \to d_i - d_{\text{min}},$$

where $d_{\text{min}} = \min(d_i)$.

2. **Binning and averaging**: The node distances and their corresponding protein signal intensities are binned along the chosen axis, into 100 bins. For each bin $B_j$, the average protein signal is computed as:

$$P_j = \frac{1}{|B_j|} \sum_{i \in B_j} I_i,$$

where $P_j$ is the average protein signal in bin $j$, $I_i$ is the protein signal at node $i$, $B_j$ is the set of nodes in bin $j$, and $|B_j|$ is the number of nodes in $B_j$. Empty bins can occur and will not have any protein signal assigned to it.

3. **Applications**: Protein profiles are used to visualize the spatial polarization of proteins on cell surfaces. When computed over time, and pooled across replicates, these profiles can be visualized as kymographs. appendix B.5.4

###### B.5.4 Kymograph

Kymographs are used to visualize the spatial distribution of cortical proteins over time. They display the corrected protein signal (appendix A.9.1) along a chosen axis (e.g., apical distance or Eap distance) as a function of time. They are generated using protein profiles (appendix B.5.3) as input. Each kymograph is visualized as a heatmap, where:

- The $x$-axis represents the spatial distance along the chosen axis (e.g. apical distance).
- The $y$-axis represents the time relative to E-division ($t_E$) or another reference time.
- The protein signal data is smoothed using a two-dimensional B-spline, fitted with cubic splines ($k_x = k_y = 3$) and with the default smoothing factor($s$) using the `scipy.interpolate.bisplrep` function.
- The color intensity represents the protein signal, with enrichment (red) and depletion (blue) highlighted using a diverging colormap. The center of the colormap is set at the mean protein intensity, while maximum and minimum color intensities are set at one standard deviation from the mean.
- Regions with no data points are masked by determining the distance ranges in smoothing intervals of 2.5 minutes, and then fitting a smoothing spline through both minima and maxima of these intervals, which outlines the region where datapoints are present. This to avoid displaying regions which are purely based on extrapolated data.

##### B.5.5 Quantification of flow vorticity

To quantify the magnitude of the embryo-wide rotational motion, we calculated the vorticity field $\boldsymbol{\omega}(\mathbf{x}, t)$, defined as the curl of the instantaneous velocity field $\mathbf{u}(\mathbf{x}, t)$:

$$\boldsymbol{\omega} = \nabla \times \mathbf{u} \tag{5}$$

The vorticity vectors were computed numerically on the mesh using the discrete derivative operator (PyVista `compute_derivative` filter). To derive a single global metric for rotation at each time step, we calculated the spatially averaged magnitude of the vorticity across all $N$ nodes in the mesh:

$$\bar{\omega}(t) = \frac{1}{N} \sum_{i=1}^{N} \|\boldsymbol{\omega}_i(t)\| \tag{6}$$

This scalar metric, expressed in $\text{min}^{-1}$, captures the global intensity of the curling motion of the tissue, independent of the specific axis of rotation.
