## Appendix C for "Integrated quantitative imaging and biomechanical modeling of early gastrulation in *C. elegans*"

### C Appendix-force models

#### C.1 Deformable cell model

##### C.1.1 Model parameters Mpacts

table C1 provides a list of all parameters used in the simulations. These parameters include 1) Mechanical parameters associated with the viscous deformable cell model, 2) parameters that govern the timing and mechanical behavior of the cytokinesis model, and 3) parameters associated with the numerical solution.

Table C1: Reference table for model parameters.

| **Parameter** | **Symbol** | **Value** |
|---|---|---|
| **Deformable cell model** | | |
| Cell radius (reference) | $R_0$ | 6.89985 $\mu m$ |
| (Interphase) surface tension | $\gamma_I$ | 0.4 nN $\mu m^{-1}$ |
| (Interphase) cell-cell adhesion density | $\omega_I$ | $\gamma_I/2$ |
| Effective range of adhesion | $h_0$ | 75 nm |
| Cytosolic bulk modulus | $K'$ | 2 kPa |
| Effective fluid viscosity | $\eta_f$ | 0.09 kPas |
| Effective cortex viscosity | $\eta$ | 11 kPas |
| Effective cortex thickness | $t$ | 300 nm |
| cell-cell contact friction | $\xi$ | 0.12 kPas $\mu m^{-1}$ |
| **Cytokinesis model** | | |
| Transition radius | $R_t$ | 1.5 $\mu m$ |
| Abscission radius | $R_f$ | 0.75 $\mu m$ |
| Mitotic surface tension | $\gamma_M$ | $3\gamma_I$ nN $\mu m^{-1}$ |
| Mitotic cell-cell adhesion density | $\omega_M$ | $\gamma_M/4$ nN $\mu m^{-1}$ |
| Daughter-half bulk modulus | $K_{div}$ | 0.25 kPa |
| Furrow ring stiffness | $k$ | 165 nN |
| Initial sub-unit length | $l_0$ | 0.6 $\mu m$ |
| Mitotic rounding duration | $\tau_m$ | 90 s |
| Division duration | $\tau_d$ | 180 s |
| **Simulation parameters** | | |
| Time step | $dt$ | 0.5 s |
| Minimal triangle area | $A_m$ | 0.25 $\mu m^2$ |
| Maximum triangle area | $A_M$ | 1.50 $\mu m^2$ |
| Conjugate gradient solver tolerance | $\epsilon_{CG}$ | $2 \cdot 10^{-5}$ |

##### C.1.2 Cell division model

*This subsection is taken from* Cuvelier et al. (2023)

We present a novel computational model of cytokinesis based on an existing deformable cell model (DCM) (Cuvelier et al., 2023). In the DCM, the cell cortex is approximated as a thin, viscous shell under surface tension, and cell-cell interactions are considered through adhesion forces. For this work, the molecular mechanisms underlying furrow ring mechanics were simplified, and a coarse-grained model was used. Here, the furrow ring is represented by a contracting ring that shrinks progressively during cytokinesis, enforcing cell division (Carvalho et al., 2009). Furrow ingression follows from tension buildup in the contractile ring and the local deformation of viscous cortical material. The simulation of cell division was performed for unconfined, spherical cells of different sizes. Constriction rates and cytokinesis durations were analyzed and compared with experimental work by Carvalho et al. (Carvalho et al., 2009), as shown in Figure S7. The experimental fit of the total duration of cytokinesis, $\tau_t$, was reproduced for stiff furrows. The model suggests that tension buildup is controlled by the balance of contractile and viscous forces in the furrow ring. Failure of furrow ingression occurs below a critical threshold when furrow stiffness is too low. The model also predicts that cell abscission occurs when the furrow perimeter reaches a critical value. During this phase, furrow ring components are disassembled, and the mother cell is split into two daughter cells. The study highlights that although furrow ring stiffness plays a dominant role, cortical viscosity and surface tension significantly influence cytokinesis dynamics.

**Furrow ring line tension** We calculated the line tension in the furrow ring of dividing spherical cells by assuming a force balance between cortex tension ($\gamma_M$) and furrow ring line tension ($F_f$) (Turlier et al., 2014). The resulting force balance equation was solved using geometrical and physical parameters from C. elegans development. For a high contractile ring stiffness, furrow radius dynamics were linear, and tension buildup was effectively controlled by furrow ring mechanics. However, as stiffness decreased, tension failed to drive ingression, leading to cytokinesis failure. The sensitivity analysis of mechanical parameters further indicated that increasing cortical viscosity delayed furrow ingression due to viscous stress buildup. A low contractile stiffness also contributed to delayed ingression, confirming experimental observations by Carvalho et al. (Carvalho et al., 2009).

##### C.1.3 Apical tension model

**Selection of triangles with increased apical tension** Apical tension is modeled by increasing the area tension of specific mesh triangles located on the apical side of the cell. The selection of these triangles involves a two-step process:

**Step 1: Initial apical triangle selection** First, the triangles belonging to a specific cell (Ea or Ep) ($T_{\text{cell}}$) are considered. The distance $\text{dist}(t)$ between the centroid of each triangle $t \in T_{\text{cell}}$ and the nearest centroid of any triangle in other cells ($T_{\text{notCell}}$) is computed. Using a threshold of contactrange $= 1\,\mu$m, the apical triangles are selected as:

$$T_{\text{apical}} = \{t \in T_{\text{cell}} \mid \text{dist}(t) > \text{contactrange}\}.$$

Here:

- $T_{\text{cell}}$: Set of triangles belonging to the queried cell.
- $T_{\text{notCell}}$: Set of triangles belonging to all other cells in the embryo.
- $\text{dist}(t)$: Distance from triangle $t$ to its nearest neighbor in $T_{\text{notCell}}$.
- contactrange: Threshold distance for excluding triangles that are close to other cells.

**Step 2: Expansion of apical triangle selection** The initial set of apical triangles ($T_{\text{apical}}$) is expanded to include neighboring triangles on the apical side. The distance $\text{distApical}(t)$ between the centroid of each triangle $t \in T$ and the nearest centroid in $T_{\text{apical}}$ is computed. Using a threshold $1.5\,\mu$m, the expanded set of apical triangles is defined as:

$$T_{\text{apicalExt}} = \{t \in T \mid \text{distApical}(t) \leq 1.5\,\mu\text{m and } t \in T_{\text{cell}}\}.$$

Here:

- $T$: Set of all triangles in the mesh.
- $\text{distApical}(t)$: Distance from triangle $t$ to its nearest neighbor in $T_{\text{apical}}$.
- $1.5\,\mu$m: Threshold for selecting triangles close to $T_{\text{apical}}$.

The final set of triangles used for increased apical tension is $T_{\text{apicalExt}}$, which includes both the initial apical patch and neighboring triangles close to it.

**Calculation of apical tension** The apical tension applied to $\gamma_{\text{apicalExt}}$ can be a constant value, or determined dynamically using a proportional-integral (PI) controller. In case of the latter, the tension is calculated based on the error between the current and target value of a specific ingression metric, such as apical area, ingression distance, or curvature:

$$\gamma_{\text{apical}} = K_p \cdot e(t) + K_i \sum_{i=0}^{N-1} e(t_i),$$

where:

- $e(t)$: The error at time $t$, defined as the difference between the target and current values of the metric.
- $K_p$: Proportional gain, which scales the immediate response to the error.
- $K_i$: Integral gain, which accounts for the accumulated error over time.
- $N = 5$: Number of previous time points used in the calculation (history buffer size).

The values of $K_p$ depend on the ingression metric used:

- For **apical area**, $K_p = 4 \times 10^7$.
- For **ingression distance**, $K_p = 0.002$.
- For **curvature**, $K_p = 2 \times 10^{-7}$.

The integral gain is set as $K_i = \frac{K_p}{10}$.

##### C.1.4 Interfacial contact model

*This subsection is taken from the Mpacts manual. Some variable names were changed for consistency with the rest of the appendix.*

This contact model generates an adhesive interfacial tension between two viscous surfaces under tension. The model relies on the following parameters:

- $\omega$ (`attrConst`): The net work of adhesion between the two surfaces.
- $h_0$ (`contact_range`): The effective range of adhesive interaction.
- $c_n$: Viscous damping constant in the normal (to the surface) direction.
- $c_t$: Viscous damping constant in the tangential (to the surface) direction.

The contact model performs two main tasks:

1. Adds a negative surface tension contribution to the triangles of interacting surfaces to reflect adhesive energy ($\omega/2$), where $\omega$ represents the net work of adhesion. This is specific to shell-like surfaces that cannot sustain in-plane stress at equilibrium.
2. Computes a localized linear pressure that scales with the overlap distance $\delta$, representing the adhesive and repulsive pressure for the (un)binding energy between the surfaces.

The adhesive energy $E_{AB}^{\text{adh}}$ for a pair of surfaces $AB$, with contact area $S_{AB}$, is defined as:

$$E_{AB}^{\text{adh}} = \left( \frac{\delta_{AB}}{h_0} - \frac{\delta_{AB}^2}{2h_0^2} \right) \omega S_{AB}.$$

The effective range of adhesion, $h_0$ (`contact_range`), is used to virtually translate the surfaces along their normal directions. The overlap distance $\delta_{AB}$ is calculated, and the contact pressure $P_{AB}^{\text{adh}}$ is given by:

$$P_{AB}^{\text{adh}}(\mathbf{x}) = k_{AB}\delta_{AB}(\mathbf{x}) - P_{AB}^0,$$

where $k_{AB} = P^0_{AB}/h_0$ is the contact stiffness and $P^0_{AB} = \omega/h_0$ ensures equilibrium ($P^{\text{adh}}_{AB} = 0$). The overlap distance $\delta_{AB}$ at a point $\mathbf{x}$ is defined as:

$$\delta_{AB}(\mathbf{x}) = \max\left(0, 2(\mathbf{x} - \mathbf{x}_{AB}) \cdot \hat{\mathbf{n}}_{AB}\right),$$

where $\hat{\mathbf{n}}_{AB}$ is the contact normal and $\mathbf{x}_{AB}$ represents the geometric center of the contact point. The resulting contact forces are computed as:

$$\mathbf{F}^{\text{adh}}_{AB} = \int_{S_{AB}} dS\, P^{\text{adh}}_{AB}(\mathbf{x}),$$

and moments are derived by integrating over the oriented contact area $S_{AB}$. To distribute the contact forces to the nodal forces, it is assumed they are collinear with the contact normal $\hat{\mathbf{n}}_{AB}$. This results in the following system of linear equations:

$$\sum_{i \in A} \mathbf{F}^{\text{adh}}_{AB,i} = -\int_{S_{AB}} dS\, P^{\text{adh}}_{AB}(\mathbf{x}),$$

$$\sum_{i \in A} \left[(\mathbf{I} - \hat{\mathbf{n}}_{AB} \otimes \hat{\mathbf{n}}_{AB}) \cdot (\mathbf{x}_i - \mathbf{x}_{AB})\right] \times \mathbf{F}^{\text{adh}}_{AB,i} = \int_{S_{AB}} dS\, (\mathbf{x} - \mathbf{x}_{AB}) \times P^{\text{adh}}_{AB}(\mathbf{x}),$$

where $A$ is the set of nodes of the interacting triangle. The drag force acting on a node $i$ of triangle $A$ is given by:

$$\mathbf{F}^{\text{fric}}_{AB,i} = \Gamma^{\text{fric}}_{AB} \sum_{k \in B} w^{AB}_{ik} (\mathbf{v}_k - \mathbf{v}_i),$$

where $\Gamma^{\text{fric}}_{AB}$ is the friction coefficient, and weights $w^{AB}_{ik}$ scale with the nodal contact forces:

$$w^{AB}_{ik} = \frac{(\mathbf{F}^{\text{adh}}_{AB,i} + \mathbf{F}^{\text{adh}}_{AB,k}) \cdot \hat{\mathbf{n}}_{AB}}{6 \sum_{k \in B} \mathbf{F}^{\text{adh}}_{AB,k} \cdot \hat{\mathbf{n}}_{AB}}.$$

The friction coefficient is computed as:

$$\Gamma^{\text{fric}}_{AB} = S_{AB}\left[c_n \hat{\mathbf{n}}_{AB} \otimes \hat{\mathbf{n}}_{AB} + c_t(\mathbf{I} - \hat{\mathbf{n}}_{AB} \otimes \hat{\mathbf{n}}_{AB})\right],$$

where $c_n$ (`c_n`) and $c_t$ (`c_t`) are the normal and tangential friction coefficients.

**References** This contact model is based on:

- Smeets et al. (2015)
- Smeets et al. (2019)
- Cuvelier et al. (2021)
- Cuvelier et al. (2023)

##### C.1.5 Eggshell

*This section is taken from Cuvelier et al. (2023). Some variable names were changed for consistency with the rest of the appendix.*

The eggshell is modeled as a convex hull surrounding the embryo.

**Contact model** The interaction between the embryo and the eggshell is described using a linear force model:

$$\mathbf{F}_c = \begin{cases} (k\delta + c_n\dot{\delta})\hat{\mathbf{n}} - c_t\mathbf{v}_t & \text{if } \delta \geq 0, \\ \mathbf{0} & \text{else,} \end{cases}$$

where:

- $\delta$: Overlap distance between the embryo and the eggshell.
- $\hat{\mathbf{n}}$: Normal unit vector at the contact surface.
- $\mathbf{v}_t$: Tangential relative velocity at the contact surface.
- $k_{\text{shell}}$: Repulsive stiffness, controlling the linear force response to overlap.
- $c_n$: Normal damping coefficient, representing resistance proportional to the normal velocity.
- $c_t$: Tangential damping coefficient, representing resistance to sliding along the contact plane.

**Parameterization**

- $k_{\text{shell}} = 10\,\text{nN}/\mu\text{m}$: Repulsive stiffness.
- $c_n = 0.00001\,\text{nN} \cdot \text{s}/\mu\text{m}$: Normal damping coefficient.
- $c_t = 0.00001\,\text{nN} \cdot \text{s}/\mu\text{m}$: Tangential damping coefficient.
